# Extreme climatic events shape adaptive potential by altering genetic variance for adult survival in a wild population

**DOI:** 10.64898/2026.10.02.755590

**Authors:** Paul Acker, Mark A. Newell, Sarah Wanless, Sarah J. Burthe, Michael P. Harris, Ella Benninghaus, Ruth E. Dunn, Carrie Gunn, Rita Fortuna, Francis Daunt, Jane M. Reid

## Abstract

Extreme climatic events (ECEs) could potentially induce episodes of rapid adaptive evolution, altering the evolutionary trajectories and viability of wild populations experiencing increasingly stochastic environments. Adaptive potential stems substantially from additive genetic variance (*V*_A_) in survival. However, the degree to which ECEs impact this variance, and the resulting evolutionary implications, have not been fully conceptualised or quantified in nature. These aims are highly challenging because underlying genetic and environmental variances have fundamentally non-linear effects on the dichotomous phenotypic outcome of survival versus death, while substantial data are required for estimation. We present a conceptual and analytical framework that formalises biological hypotheses for how *V*_A_ depends on environmental severity, how such dependences are reshaped by non-linear genotype-environment-phenotype relationships and demographic structure, and the implications for evolutionary responses. We then use 25 years of survival and pedigree data from European shags (*Gulosus aristotelis*) to decompose variance in adult survival across diverse environmental conditions including ECEs. Using a capture-recapture animal model, we demonstrate non-negligible *V*_A_ underlying adult survival across all years combined, implying notable adaptive potential. Yet, partitioning by environmental severity revealed substantial *V*_A_ in good years with high survival but negligible *V*_A_ in bad years with ECE-induced low survival, implying a genotype-by-environment interaction that constrains adaptive potential under severe conditions. However, scale transformations showed how the non-linear genotype-environment-phenotype relationship can distort or even reverse such patterns, by reducing the extent to which *V*_A_ is phenotypically expressed in good conditions with high mean survival and amplifying it in more severe conditions with lower mean survival. Evolutionary responses to severe conditions could consequently be non-negligible even given small underlying *V*_A_. We thereby demonstrate how ECEs can reshape adaptive potential, highlighting the need to integrate impacts of environmental severity on genetic variation in fitness components into predictions of population resilience under climate change.

**TEASER TEXT:** Extreme climatic events (ECEs) could substantively reshape wild populations’ capacities for adaptive micro-evolution, but such effects have not yet been rigorously conceptualised or quantified in nature. We derive a general quantitative genetic framework that formalises how environmental severity could modulate adaptive potential by altering additive genetic variance in survival. We apply this framework to adult annual survival in a wild population of European shags, considering differing levels of environmental severity reflecting occurrences of ECEs. Notably, underlying additive genetic variance was substantial in good conditions, but clearly smaller in bad conditions. Yet, among-genotype variance manifested on the expressed phenotypic scale of survival versus death can be magnified by low survival in bad conditions, potentially enhancing the scope for evolutionary adaptation. We thereby show how ECEs can simultaneously constrain and restructure adaptive potential, implying that quantitative genetic impacts of environmental severity must be considered when predicting population resilience to climate change.

## INTRODUCTION

Quantifying the extent to which extreme climatic events (ECEs) can trigger rapid adaptive evolution, and how such responses differ from those evident in typical environments, is necessary to predict whether evolution can enable population persistence as climate change increases the frequency of ECEs (van de Pol et al., 2017; Maxwell et al., 2019; Baeckens & Donihue, 2025; Heinen et al., 2025). In principle, a population’s capacity for rapid adaptive evolution (its ‘adaptive potential’) is determined by the magnitude of standing additive genetic variance (*V*_A_) for key traits that contribute to fitness, because this variation induces natural selection driving heritable phenotypic change (Lande & Arnold, 1983; Hansen & Houle, 2008; Hendry et al., 2018). Since survival is a core fitness component and a primary demographic route by which ECEs threaten population viability (Bailey & van de Pol, 2016; Baeckens & Donihue, 2025), quantifying *V*_A_ in survival, and quantifying the impact of ECEs on such *V*_A_, is prerequisite for predicting evolutionary responses to climate change. Yet, despite extensive evidence of profound physiological and ecological impacts of ECEs (Parmesan et al., 2000; Reyer et al., 2013; van de Pol et al., 2017), rigorous conceptualisations and empirical estimates of how ECEs alter *V*_A_ in survival, and hence shape adaptive potential, are still lacking (Baeckens & Donihue, 2025; Heinen et al., 2025).

Wider theory and empirical results imply that environmental changes could substantively affect adaptive potential, thereby determining evolutionary responses to increasingly severe and/or stochastic environments. Indeed, *V*_A_ in diverse phenotypic traits (spanning morphology, physiology, behaviour, and life-history) frequently differs between favourable and unfavourable conditions, reflecting genotype-by-environment interactions (G×E; Hoffmann & Merila, 1999; Charmantier & Garant, 2005; Rowin ski & Rogell, 2017). *V*_A_ may increase with environmental severity (Figure 1ai), representing release of previously ‘cryptic’ genetic variation by stressful conditions that compromise robustness of developmental and self-maintenance systems (Gibson & Dworkin, 2004; Masel & Trotter, 2010; Paaby & Rockman, 2014). Yet, such releases may be curtailed under even more severe conditions where developmental and self-maintenance systems reach hard biophysical limits independent of standing genetic variation (Blows & Hoffmann, 2005; Angilletta, 2009). *V*_A_ could then decrease, generating an overall ‘hump-shaped’ relationship with environmental severity (Fig 1aii). Conversely, *V*_A_ could be minimal at intermediate severities (Fig 1aiii) if genetic variation was previously eroded under such environmental conditions given their relatively common occurrence and strength of selection (Hoffmann & Merila, 1999). Alternatively, genetic effects and resulting *V*_A_ could be unaffected by environmental severity (Fig 1aiv). However, beyond these general scenarios, rationalising how environmental deteriorations (including ECEs) affect adaptive potential by altering *V*_A_ in the key fitness component of survival additionally requires formalising the implications of the inherently non-linear relationship that links genetic and environmental variation to the discrete phenotypic outcome of survival versus death.

**Figure 1.**
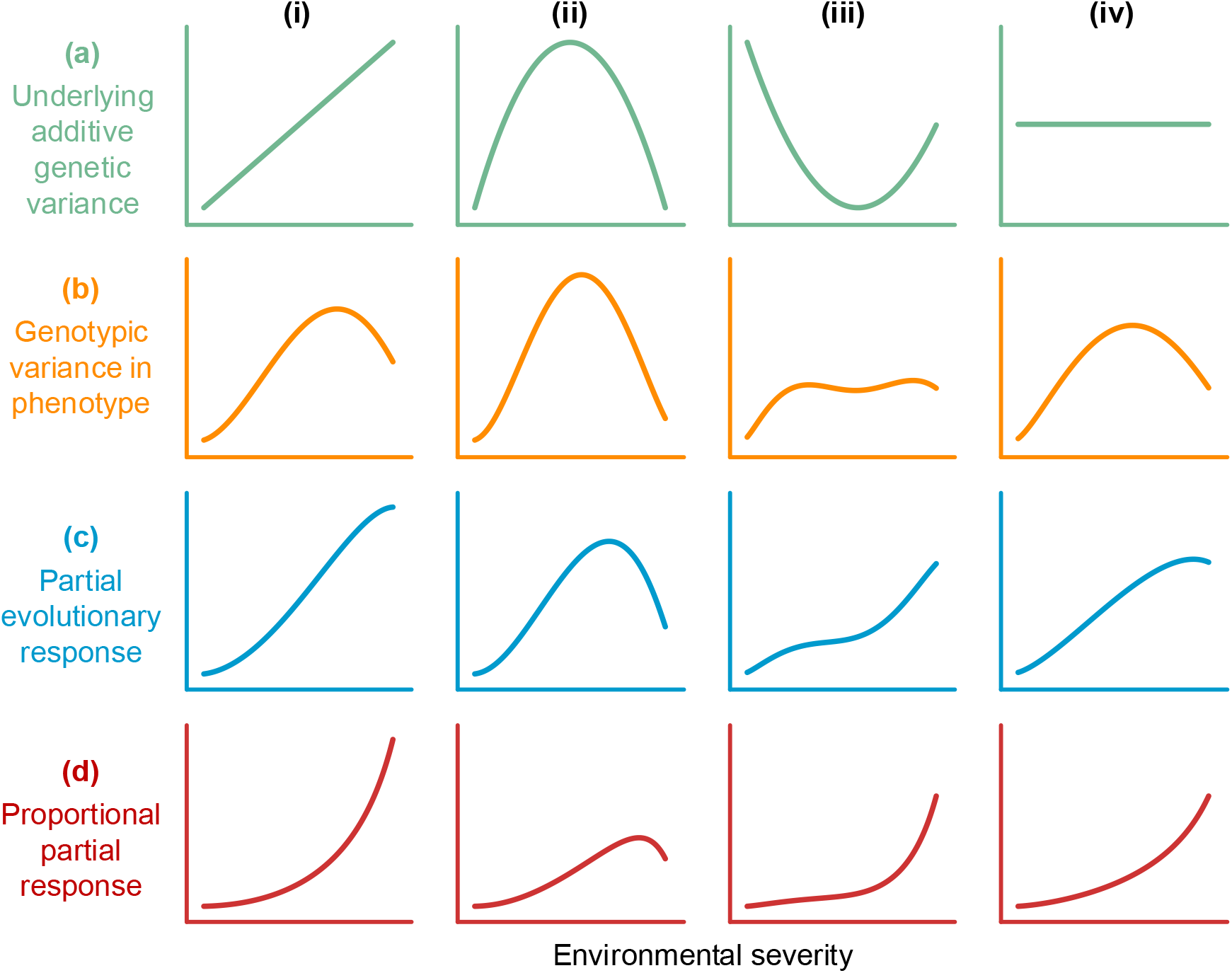
Hypothetical relationships between environmental severity (defined as 1 minus mean survival, i.e. 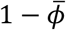, giving the mortality rate) and (a) latent-scale additive genetic variance (*V*_A_) underlying survival, (b) phenotypic-scale genotypic variance in survival (*V*_*ϕ*/_*C*_;A_), and the partial evolutionary responses measured as the (c) absolute and (d) relative change in mean survival (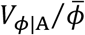 and 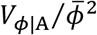 respectively). Y-axis scales are identical across the four panels within each row. For illustration, all graphs were drawn using environmental severity spanning 0.02–0.82, with *V*_A_ spanning 0.075–0.30 (probit-scale values). Axis graduations are omitted to emphasise qualitative transformation patterns from (a) to (d), and because similar functional forms in (b-d) can arise from the same starting relationships in (a) but with alternative parameter values (Appendix S1).

To derive an appropriate quantitative framework, survival can be conceptualised within the established quantitative genetic paradigm for dichotomous traits. Here, genetic effects of numerous loci are postulated to act alongside environmental effects on a latent unbounded continuous scale and translate non-linearly onto the phenotypic scale in the form of probabilities, *ϕ*, of the discrete outcome (Figure 2, Appendix S1; Dempster & Lerner 1950; Falconer & McKay, 1996; Acker & Reid, 2025). *V*_A_ underlying survival variation is therefore defined on a latent scale. Due to the intrinsic sigmoidal genotype-environment-phenotype relationship, the expressed among-genotype variance in survival arising from *V*_A_, termed ‘phenotypic-scale genotypic variance’ *V*_*ϕ*/_*C*_;A_ (Appendix S1), can change substantially with the population mean even given constant *V*_A_. Specifically, *V*_*ϕ*/_*C*_;A_ will increase (or decrease) as mean survival, 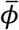, moves towards (or away from) 0.5 (Figure 2; Acker & Reid, 2025). Hence, decreases in 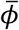 caused by ECEs will inevitably modulate how underlying *V*_A_ is expressed as survival variation, thereby inherently reshaping the amount of adaptive potential effectively exposed to direct selection irrespective of whether underlying *V*_A_ also changes. Consequently, linear, hump-shaped, U-shaped, or flat relationships between *V*_A_ and environmental severity on the latent scale (Figure 1a) will transform into different relationships on the phenotypic scale. For example, linear can become hump-shaped, U-shaped can be flattened, and flat can become hump-shaped (Figure 1b; Appendix S1). Explicitly quantifying ECE-induced changes in latent-scale *V*_A_, and its phenotypic-scale expression *V*_*ϕ*/_*C*_;A_, is therefore essential for understanding how adaptative potential is intrinsically underpinned and functionally manifested in differing environmental conditions.

**Figure 2.**
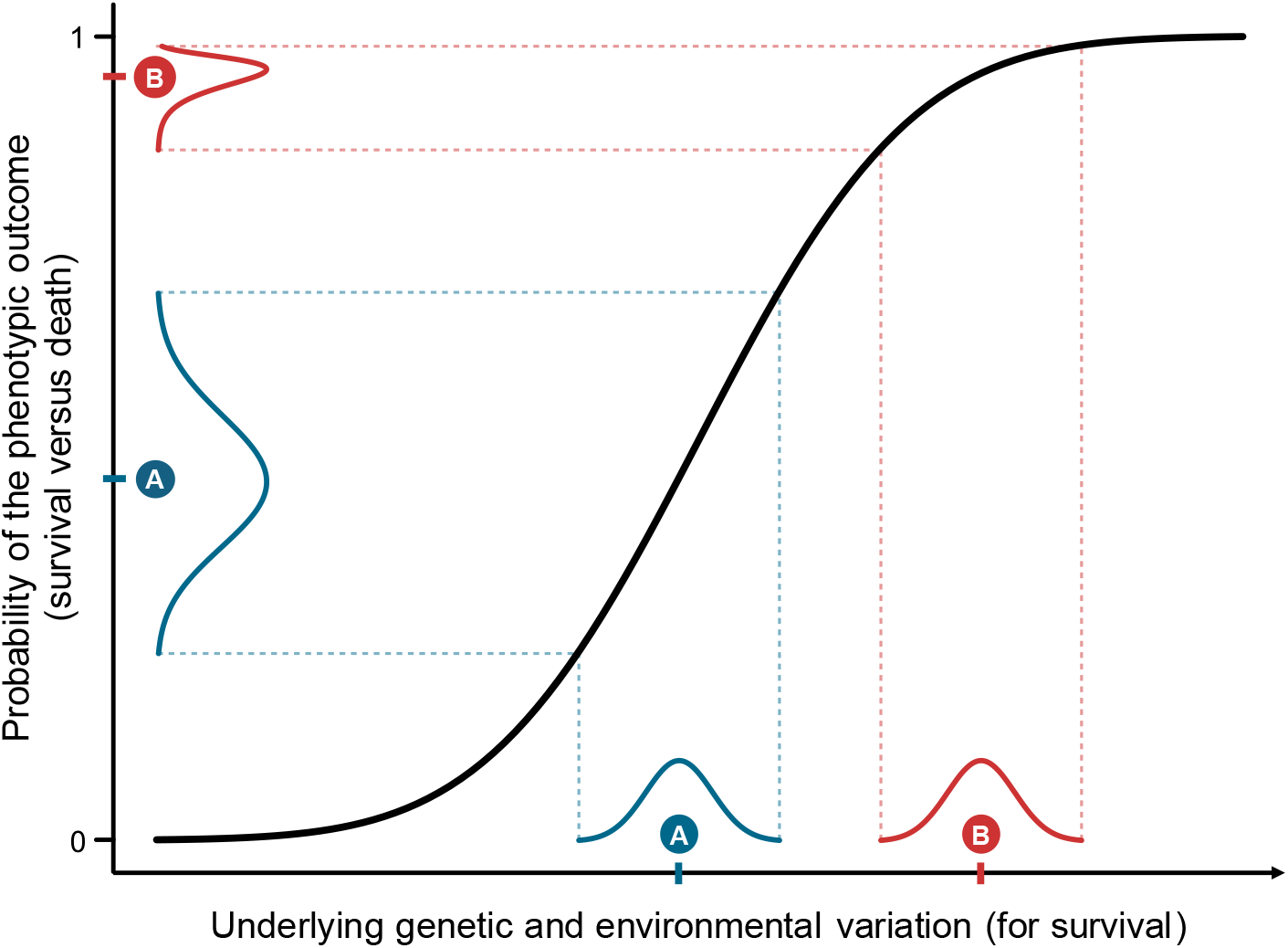
The non-linear (sigmoidal) genotype-environment-phenotype relationship (black line) typically postulated to link the probability, *ϕ*, of any dichotomous phenotypic outcome to a latent continuous predictor conceived as the linear combination of underlying genetic and environmental effects. The density curves on the x-axis (A and B) represent hypothetical distributions of underlying additive genetic effects with equal variances (i.e. equal latent-scale *V*_A_) but different means (ticks), exemplifying the same population placed under bad vs good environmental conditions. The arising phenotypic-scale genotypic variance (i.e. *V*_*ϕ*/_*C*_;A_, the variance in phenotypic expression among genotypes represented by density curves A and B on the y-axis) is maximised given population mean phenotype (i.e. mean outcome probability, 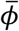) close to 0.5 (e.g. A) and decreases with mean phenotype close to 0 or 1 (e.g. B). Such sigmoidal relationships are enacted by any binary generalised linear mixed model (‘GLMM’; illustrated with a probit link, but other links such as logit or robit imply similar relationships). Note that on the underlying latent scale, variation is depicted in units with constant proportionality of the residual standard deviation for any focal genotype and/or environment, and our simple illustration assumes no underlying G×E (see Appendix S1 for mathematical explanations).

Evolutionary implications can then be quantified, and comparative analyses potentially facilitated, using metrics grounded in the mathematical reasoning of the Price equation (Appendix S1; Price, 1970). Specifically, noting that 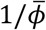 is the gradient of selection on survival (Lande & Arnold 1983), 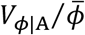 quantifies the ‘partial’ evolutionary response, defined as the increase in 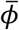 expected from the underlying change in allele frequencies due to survival selection, conditional on the phenotypic (i.e. survival) effects of the alleles under the focal pre-selection conditions (Price, 1972; Ewens, 1989; Walsh & Lynch, 2018). This evolutionary response is ‘partial’ because it excludes any possible additional shift in 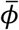 given concurrent genetic and environmental changes that define post-selection conditions (e.g. gene flow, further selection, or population dynamics) and will alter the cross-generational transmission of phenotypic effects (Appendix S1). Further, the quantity 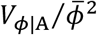 measures the same partial evolutionary response but as a proportional increase, as often advocated for cross-system comparisons (Hendry et al., 2018; Bonnet et al., 2022). These two metrics capture the fact that, independently of any change in *V*_*ϕ*/_*C*_;A_, more severe conditions directly promote greater adaptive evolution by reducing 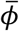 and thus increasing the selection gradient. Accordingly, relationships with environmental severity are again reshaped, making flat or hump-shaped curves tend towards essentially monotonic increases (Figure 1c-d, Appendix S1).

Further, beyond quantifying multi-scale responses of *V*_A_ to environmental severity (Figure 1), any comprehensive assessment of adaptive potential in varying environments should also quantify cross-environment correlations in additive genetic effects (Sgro & Hoffmann, 2004; Kruuk et al., 2008). Such correlations could be strongly positive or negative, respectively implying that alleles that increase *ϕ* under good conditions increase or decrease *ϕ* under bad conditions. If the magnitudes of environment-specific *V*_A_ and the non-linear genotype-environment-phenotype relationship translate such correlations into substantial cross-environment covariances in *ϕ*, then evolution induced by bad conditions could enhance or impede adaptation to good conditions (and vice versa). Conversely, correlations could be weak if largely distinct traits determine survival under contrasting conditions, or effectively null if *V*_A_ is negligible in one or multiple environments. Adaptation to environments where *V*_A_ is non-negligible would then proceed unconstrained. Moreover, *V*_A_ and 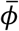 could also be sex-specific, to degrees that could change with environmental conditions (Poissant et al., 2010; Han & Dingemanse, 2017). Coupled with any cross-sex genetic correlations, resulting non-linear effects on *ϕ* could generate strongly sex-specific adaptive potentials, further affecting population outcomes (Kruuk et al., 2008; Wolak et al., 2018). Yet, how environmental conditions and sex shape *V*_A_ underlying survival, and affect its phenotypic expression (generating environment and sex-specific *V*_*ϕ*/_*C*_;A_), remains largely unquantified in wild populations.

This gap likely remains because quantifying environmentally-induced dynamics of *V*_A_ in survival entails major data and analytical challenges. Estimation of latent variances underlying dichotomous phenotypes requires particularly large phenotypic and relatedness datasets spanning many individuals, especially when phenotypic means (here survival probabilities) can be high or low, implying that most underlying genetic variation is cryptic (Acker et al., 2023a; Figure 2). Further, datasets must span multiple years including environmental conditions that are severe enough to overcome demographic buffering (i.e. canalisation of *ϕ*; Hilde et al., 2020) and reduce [inlne], such as ECEs (Frederiksen et al., 2018; Bailey & van de Pol, 2016). Advanced non-linear quantitative genetic ‘animal models’ and complex scale transformations are then required to fully partition latent and phenotypic variation in survival (Appendix S1; de Villemereuil et al., 2016; Acker & Reid, 2025). Partly due to these challenges, few wild population studies have estimated *V*_A_ in survival, and none have quantified effects of environmental severity (Papaï x et al., 2010; Morrissey et al., 2012; Stoffel et al., 2017; Wolak et al., 2018; Moiron et al., 2022). Available estimates therefore encompass mixtures of environmental conditions, which could obscure the adaptive potentials specific to particular degrees of severity, and hence mislead on adaptive potential given environmental change (Fry 1992). Consequently, no studies have conceptualised or quantified the full hierarchical sequence of metrics informing on environmentally-induced variation in adaptive potential (Figure 1).

Accordingly, we applied our general conceptual framework (Figure 1, Appendix S1) to test whether environmental severity shapes *V*_A_ in survival, and resulting adaptive potential, using advanced multi-scale quantitative genetic analyses of 25 years of survival and pedigree data from a wild population of European shags (*Gulosus aristotelis*, hereafter ‘shags’). We focused on annual adult survival, which is typically the predominant cause of fitness variation in long-lived species, constituting the primary vital rate expected to drive or constrain adaptive evolution and population persistence (Sæther & Bakke, 2000; Hilde et al., 2020). Moreover, the population’s dramatic among-year variation in adult survival due to occasional ECEs (Frederiksen et al, 2008; Acker et al., 2021a) provided a valuable opportunity to quantify *V*_A_ across a broad range of environmental severity. To achieve this, we first estimated annual sex-specific survival probabilities and used these to define a biologically-relevant axis of environmental severity. We then estimated *V*_A_ in survival across the full dataset and within three categories of severity comprising good, bad and very bad years, alongside cross-environment genetic correlations. By transforming resulting estimates onto the phenotypic scale and deriving partial evolutionary responses, we reveal how adaptive potential stemming from *V*_A_ in adult survival can be reshaped by ECEs, thereby constraining population responses to climate change.

## MATERIAL & METHODS

### Field system and data collection

The required data were collected during breeding seasons (typically April–July) on the Isle of May National Nature Reserve (hereafter ‘IoM’), Scotland (56°11′5.40″ N, 2°33′16.19″ W). Since 1997, most chicks and breeding adults (typically ≥3 years old) have been marked with uniquely coded colour rings, which are field-readable at distances up to ∼150m with a telescope, enhancing preceding standard ringing that commenced in 1961 (Appendix S2; Harris et al. 1994, 1998; Frederiksen et al., 2008; Acker et al., 2021a,b). Consequently, increasingly large proportions of breeding adults were individually identifiable (reaching ∼95% in 2009–2018; Acker et al., 2021b). Systematic checks of nest and roost sites yielded high, though not perfect, annual adult detection probabilities (Acker et al., 2021a). Most adults were sexed through vocalisations and/or genotyping sex-linked markers. Ringing was licensed by the British Trust for Ornithology (permits A400 and A4607). Work on IoM was annually licensed by NatureScot (permit MON/RP24/2 and its predecessors).

We focused on biological years (defined between consecutive breeding seasons) 2000–2024, which provided sufficient resightings of sexed adults spanning all breeding ages to accurately quantify temporal and individual variation in survival, and parental identity (i.e. pedigree) data for estimating *V*_A_ (Appendix S2). The dataset comprised 12,820 breeding season sightings of 1,475 females and 1,498 males of known age (first ringed as chicks, mean breeding age 7 years, range 2–25), and 17,587 sightings of 2,096 females and 2,016 males including individuals of unknown age (first ringed as adults).

### Population-level variation in adult annual survival

We first fitted a capture-recapture model to quantify sex-specific annual survival probabilities from the full ring-resighting dataset, thereby revealing the survival impacts of annual environmental conditions (while handling imperfect detection; Lebreton et al., 1992; Gimenez et al., 2008). Specifically, we defined a Bayesian discrete-time Cormack-Jolly-Seber model where each individual starts as ‘alive’ in its first observed breeding season during 2000–2023, then remains alive or transitions to ‘dead’ between subsequent breeding seasons given survival probability *ϕ*_*s,t*_, and is observed or not given resighting probability *p*_*s,t,r*_ (where s, *t*, and *r* index year, sex, and ring type, respectively; Appendix S2). Since there is virtually no breeding dispersal from IoM, *ϕ*_*s,t*_represents true adult survival with no confounding permanent emigration (Barlow et al., 2013; Acker et al., 2021a). To ensure independent estimation of *ϕ*_*s,t*_and *p*_*s,t,r*_ in the final year, we modelled an additional occasion in autumn 2024 (for which we had numerous resightings; Appendix S2). From the primary model estimates, we derived the posterior distributions of the year-specific population mean survival 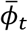 across both sexes (Appendix S3) and measured environmental severity as 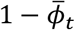. This metric is biologically meaningful because it effectively quantifies the degree of maladaptation of the population across all (unmeasured) dimensions of environmental variation (Figure 1; Maynard Smith, 1976; Chevin, 2013; Brady et al., 2019).

These analyses revealed notable temporal variation in 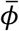 (ranging ∼0.18–0.98; Figure 3a), in line with previously documented long-term variation in the focal population (Harris et al., 1998; Frederiksen et al., 2008; Acker et al., 2021a,b, 2023b). Most years were characterised by high 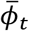 typically around ∼0.9, with little or no sex difference. Yet, multiple years showed much lower 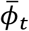 coinciding with rough winter weather including ECEs comprising storms where strong winds, rough seas and/or heavy rainfall combined with low temperatures and short daylength heighten thermoregulatory costs while restricting foraging, causing mass mortality (Appendix S2; Daunt et al., 2006; Frederiksen et al., 2008; Acker et al., 2021a). Further, *ϕ*_*s,t*_was clearly lower in females than males in several low-survival years, and the sex difference increased substantially with environmental severity across all years except the most severe (Figure 3a-b; Appendix S4). This reveals notable sex-specific environmental tolerance, implying sex-specific severity of environmental conditions and potentially generating sex-specific adaptive potential on the phenotypic scale of survival (due to the mean-dependence; Figures 1-2, Appendix S1).

**Figure 3.**
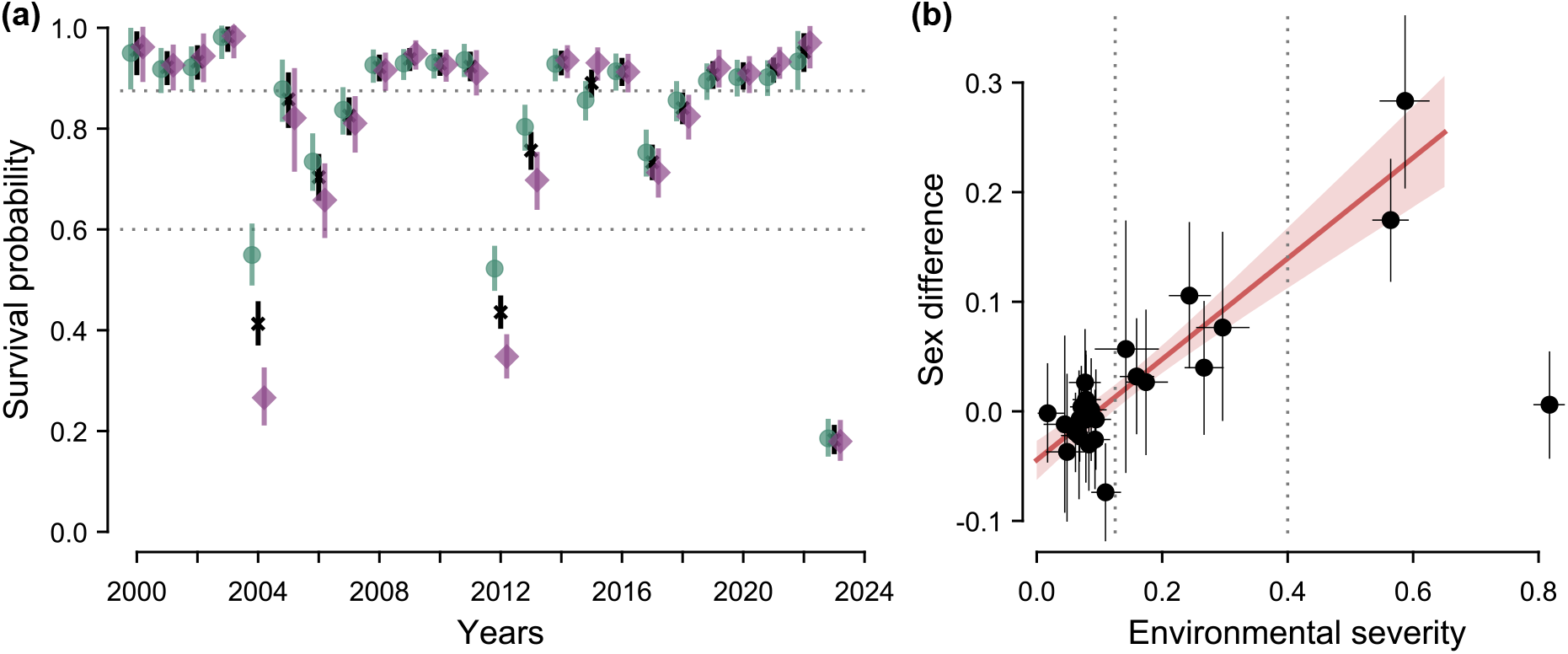
Estimates of (a) annual sex-specific and population mean survival probabilities (*ϕ*_*s,t*_and 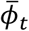), and (b) the sex difference in survival probability in adult shags during 2000–2024. Dotted lines indicate the defined boundaries between years with good, bad, and very bad environmental conditions (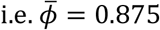 and 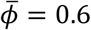). Points and error bars indicate posterior means and 95% credible intervals. On (a), green circles and purple diamonds represent males and females, respectively, while the black crosses show the population mean. On (b), the sex difference is defined as males minus females, and environmental severity is defined as 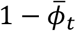. For illustration, the red line and pink shading indicate the posterior mean and 95% credible intervals of the ordinary least square regression including all years except 2023 (when the most extreme environmental conditions occurred, bottom-right corner point).

### Overall quantitative genetic decomposition

To estimate *V*_A_ underlying survival across environmental conditions, we integrated a quantitative genetic ‘animal model’ into the capture-recapture framework. We thereby decomposed variation in *ϕ*_*i*.*t*_ (where _*i*_ indicates individual identity) onto the underlying continuous latent scale, effectively by embedding a binary generalised linear mixed model (‘GLMM’) with a probit link (Appendix S1; Papaï x et al., 2010; de Villemereuil et al., 2016; Acker & Reid, 2025). Specifically, we formulated probit(*ϕ*_*i,t*_) as a linear combination of key sources of variation, first by including a ‘random’ effect *c*_*i*_ representing consistent among-individual heterogeneity spanning diverse environmental conditions:

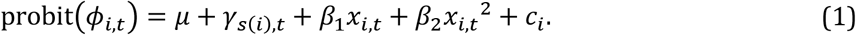

Here, _*μ*_ is the overall intercept, _*s*_(_*i*_) is the sex of individual _*i*_, *γ*_*s(i),t*_ is the ‘random’ year×sex effect distributed with sex-specific variance (given Figure 3) such that *γ*_*s(i),t*_∼N(0,*V*_*γ,s(i)*_), *x* is age, and *β*_1_ and *β*_2_ are linear and quadratic effects of age (thereby accounting for actuarial senescence previously documented in shags; Harris et al., 1994), and *c*_*i*_∼N(0,*V*_*C*_) where *V*_*C*_ is the among-individual variance, representing the magnitude of individual heterogeneity in ‘frailty’ (Cam et al., 2002; Gimenez et al., 2017). Since Equation 1 formulates a phenotypic expectation (the probability *ϕ*_*i,t*_), with the residual variance effectively defined as unit variance on the probit scale (Acker & Reid, 2025), no additional residual is included. Notably, fitting this initial model to the dataset for all known-age adults revealed among-individual variance underlying adult annual survival (posterior distribution of *V*_*C*_ concentrated away from zero, with mean [95% credible interval]: 0.058 [0.001, 0.138]--; Appendix S4).

We then further decomposed this heterogeneity into an additive genetic effect a_*i*_ (‘breeding value’) and a permanent individual effect *b*_*i*_ (which could comprise non-additive genetic and/or environmental components):

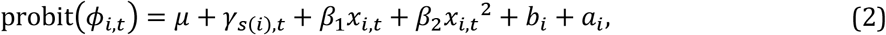

where b_*i*_∼N(0,*V*_B_), ***a***∼N(0,***A****V*_A_), and *V*_*B*_ is the permanent individual variance, a_*i*_ are the elements of vector ***a***, and ***A*** is the additive genetic relationship matrix. Since shags are socially monogamous with biparental care and little extra-pair paternity, putative parents were directly assigned to chicks from field observations, generating a dense pedigree (full details in Appendix S2; Acker et al., 2023a). This provided the relatedness data required to define ***A***, which allows estimating *V*_A_ and *V*_B_ by formalising a covariance structure for ***a*** corresponding to how polygenic inheritance generates trait resemblance among individuals (Kruuk, 2004). We implemented this covariance structure through an algorithm that formulates a directed acyclic graph, implicitly defining ***A*** by explicitly representing parent–offspring transmission of additive genetic effects (Appendix S2).

### Environment-specific quantitative genetic decomposition

We then extended the above model (Equation 2) to achieve our core objective of quantifying *V*_A_ given different levels of environmental severity:

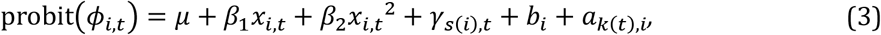

where *k(t)* indicates the environmental conditions in year *t*, and 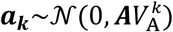. In principle, *V*_A_ could change as a continuous function of environmental severity (Figure 1). However, in practice, unconstrained estimation of 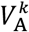, and particularly of any cross-environment covariances, becomes achievable by considering discrete environments (Appendix S4; e.g. McAdam & Boutin, 2003; Pelletier et al., 2007; Fortuna et al., 2024). Accordingly, we defined three categories of environmental severity comprising fifteen ‘good’ years with 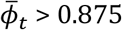 (where also one or both sexes had 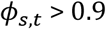, six ‘bad’ years with 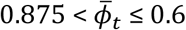, and three ‘very bad’ years (abbreviated *vbad* for indexing) with 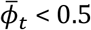 (Figure 3), and hence estimated three distinct 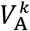 parameters.

Here, we assumed that *b*_*i*_ is constant, representing individual differences in frailty that arise from early development and persist through all future environmental conditions (Lindstro m 1999; Cam et al., 2002; Gimenez et al., 2017). However, models with environment-specific permanent individual effects *b*_*i,k*_ (and variances 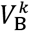) yielded quantitatively similar conclusions (Appendix S3). Moreover, models that additionally included random effects of brood, maternal, or paternal identity returned similar values of 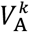 to models without these effects (and their estimated variances were small). 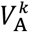 was therefore not discernibly inflated by shared early-life conditions among siblings (Appendix S3).

We fitted further models that included explicit pairwise genetic correlations among the good, bad and very bad environments, or that estimated sex-specific *V*_A_ and 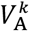and the cross-sex genetic correlations (Appendix S4). Finally, we tested alternative two-level environmental categorisations, where the bad years were grouped with the good or the very bad years (Appendix S4).

### Model inference and checks

All models were coded in Stan (Stan Development Team, 2026), which enacts Bayesian inference through Hamiltonian Monte Carlo, run using package rstan (Carpenter et al., 2017) in R (R Core Team, 2025). We specified weakly informative priors for all parameters (Appendix S3). Details of posterior sampling procedures, MCMC diagnostics (showing good convergence), and parameter estimates are in Appendices S3-S4. Full numerical summaries and posterior samples of all parameters are available from Zenodo alongside Stan and R scripts (see Data and code availability; Acker et al., 2026). Estimates are presented as posterior means along with 95% credible intervals; modes, medians, and 50% credible intervals are also shown on graphical representations of skewed posterior distributions.

We implemented two major checks to evaluate the robustness of our analyses and extent to which the dataset could effectively support our objectives. First, we fitted a ‘control’ model after randomising individual identities which, as expected, yielded negligible 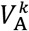 estimates in all three defined environments (Appendix S3). This confirms that estimates are not spuriously inflated, and provides benchmark posterior distributions given null 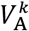. Second, we used simulations to assess our capability to detect non-negligible 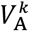 given limited pedigree informativeness in bad and very bad years, as is inevitable due to restricted within-family exposure to rare environmental conditions. Here, we simulated a heritable dichotomous trait using the same pedigree and number of observations per individual in each defined environment as in our real dataset, and three values of 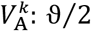, ϑ, and 2ϑ, where ϑ was the highest posterior mean obtained with the real dataset (Appendix S3). Fitting animal models analogous to Equation 3 revealed high chances of detecting non-negligible 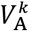 in all three environments given 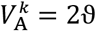, still high chances in the good and bad years for 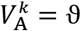, but restricted chances in all three environments given 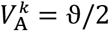 (Appendix S3).

### Derived parameters and phenotypic-scale inference

To implement our full conceptual framework and further quantify how genetic and environmental variation underlying survival shape adaptive potential, we calculated and summarised posterior distributions of key metrics derived from the primary estimated model parameters (Appendix S3).

First, we calculated the environment-specific heritability comprising the proportion of latent-scale variance explained by additive genetic effects (conditional on sex, age, and year variation): 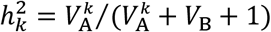 (de Villemereuil et al., 2016). To explicitly test for differences in 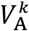 (and hence in 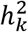) between environments we calculated the statistical support for pairwise differences between defined environments {*m,n*} as the posterior probability 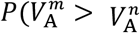; Appendix S3). Values near 0 or 1 indicate clear evidence for a difference (of either sign), while values near 0.5 indicate no evidence (Gelman et al., 2013).

Second, we calculated how adaptive potential is reshaped given the non-linear translation of underlying additive genetic variation into survival outcomes (see full derivations and explanations in Appendix S1; Figure 1b). We back-transformed the probit-scale estimates of 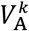 onto the dichotomous phenotypic scale of survival, yielding the phenotypic-scale genotypic variance *V*_*ϕ*/A_ arising from latent-scale breeding values independently of other probit-scale effects (Appendix S1). Yet, the non-linear genotype-environment-phenotype relationship makes the back-transformation conditional on other probit-scale effects, specifically sex, year and age (Figure 2; Appendix S1; Acker et al., 2023a; Acker & Reid, 2025). For simplicity, we focussed on the sex×year-specific back-transformed variance, 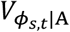, for individuals aged 7 years (the mean age in our dataset; and additionally considered ages 3 and 15 in Appendix S4). Critically, 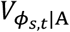 quantifies the expressed variation in a given sex and year which directly undergoes selection to generate a shift in the underlying distribution of additive genetic effects that potentiates evolution (Appendix S1).

Finally, we calculated 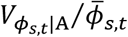 and 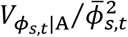 (where 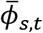 is the back-transformed sex×year-specific mean survival at age 7), representing partial evolutionary responses quantified respectively as absolute and proportional increases in 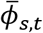 (Figure 1c-d; Appendix S1).

## RESULTS

### Overall quantitative genetic decomposition

Our quantitative genetic decomposition encompassing all years (Equation 2) revealed non-zero *V*_A_ underlying annual adult survival (posterior mean [95% credible interval]: 0.035 [0.001, 0.081]). Despite a modest posterior mean, the posterior distribution peaked away from zero and was distinct from the ‘control’ null expectation (Figure 4a), providing clear evidence of adaptive potential. In contrast, the permanent individual variance (*V*_B_: 0.034 [0.000, 0.110]) had a prominent posterior mode on zero (Appendix S4). Consequently, the overall latent-scale heritability *h*_2_ was small but most likely non-zero (0.03_2_ [0.000, 0.074]). There were also clear age effects implying slightly increasing annual survival probability from age 2 to ∼7 years, then slightly decreasing until ∼11 years, and sharply decreasing subsequently; Appendix S4). This represents notable actuarial senescence and, given the sex differences in survival probability in bad and very bad years (Figure 3), implies sex×age severity of environmental conditions (Figure 2).

**Figure 4.**
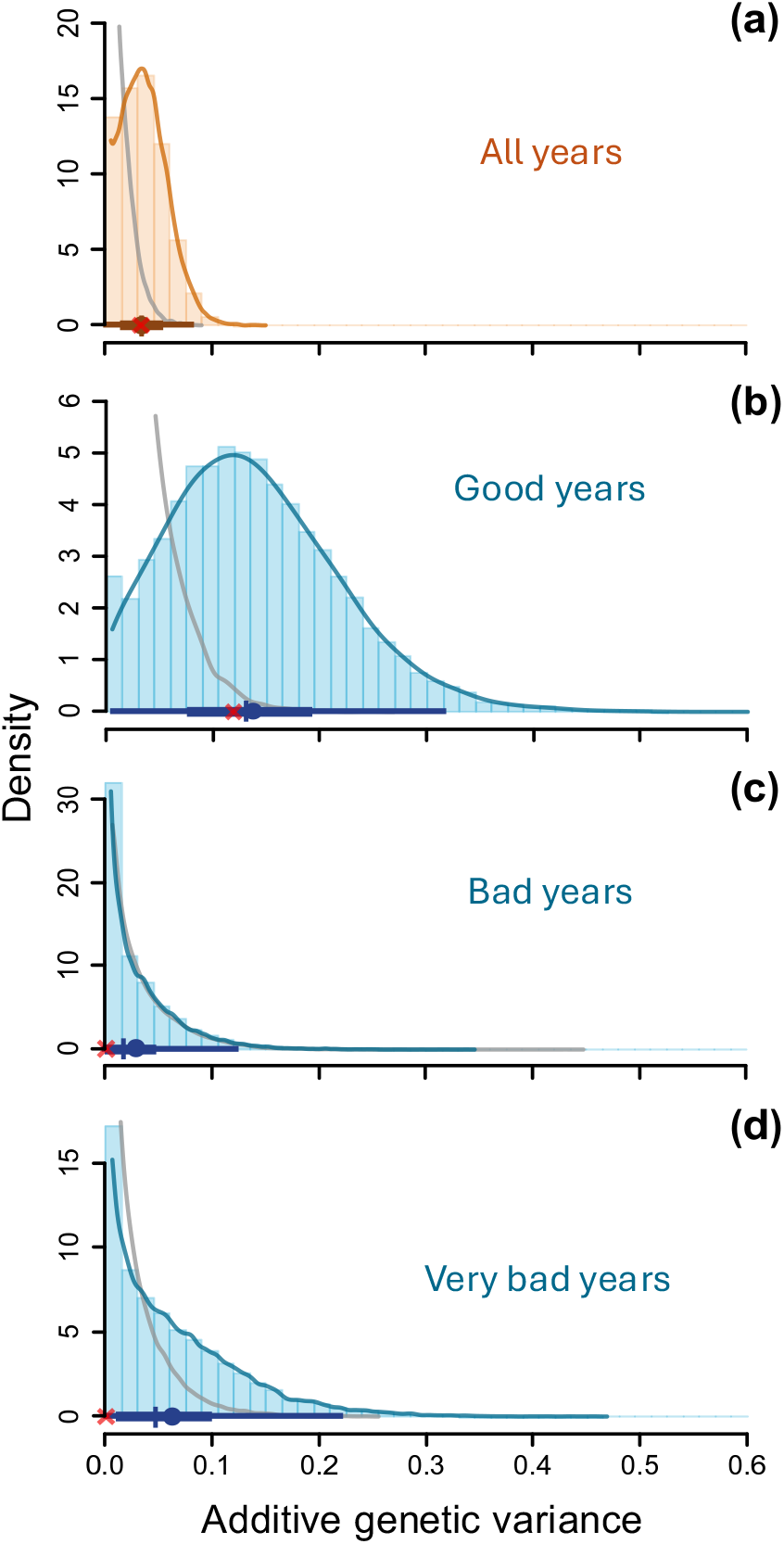
Posterior distributions of additive genetic variance underlying annual adult survival estimated (a) across all years (*V*_A_), and separately in years with (b) good, (c) bad, and (d) very bad environmental conditions 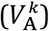. Histogram bars show the distributions of the posterior samples. Solid lines show the corresponding kernel density estimates. The superimposed thin and thick horizontal lines show the 95% and 50% credible intervals, respectively. Along these horizontal lines, the point, vertical tick and red cross show the mean, median, and mode, respectively. The grey curves show the densities of the posterior distributions for the corresponding ‘null expectation’ estimated after randomising individual identities (see also Appendix S4).

### Environment-specific quantitative genetic decompositions

The model that explicitly considered three defined categories of environmental severity (Equation 3) revealed large additive genetic variance underlying annual adult survival in good years 0.138 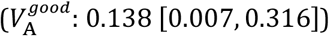, with considerable posterior density far from zero and from the null expectation (Figure 4b). Meanwhile, values for bad and very bad years were small 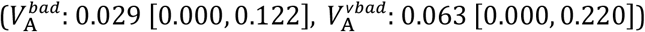 with posterior distributions similar to their null expectations (Figure 4c,d). Consequently, 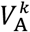 was larger in good years than bad years 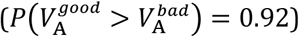, implying a G×E between low and intermediate environmental severities. There was weaker evidence for a difference in 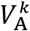 between good and very bad years 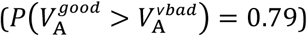, due to the substantial uncertainty for 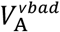. The environment-specific estimates yielded latent-scale heritability that was notably high in good years 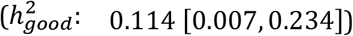, negligible in bad years 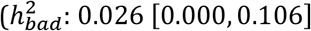 and still small but more uncertain in very bad years 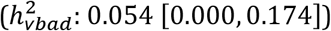

As expected given negligible 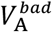 and 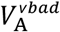, estimates of cross-environment genetic correlations were largely inconclusive (i.e. very uncertain and widely overlapping zero), although the possibility of a strong negative correlation between good and very bad years was largely excluded (Appendix S4). We found no clear difference in additive genetic variance between the sexes, or any cross-sex genetic correlation, as halving the sample sizes of individuals yielded reduced power to estimate sex-specific values given present magnitudes of 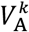. However, there was more evidence for larger 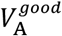 in females (vs males), and for a positive cross-sex correlation in good years (vs a negative one; Appendix S4).

Alternative models with two instead of three environmental categories neither challenged nor fundamentally refined the above conclusions. Specifically, grouping good and bad years yielded small *V*_A_ as in the overall model but with larger uncertainty, while grouping bad and very bad years yielded slightly smaller *V*_A_, with smaller uncertainty, than the main estimate of 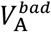 (Appendix S4).

### Phenotypic-scale implications

Because the high survival probabilities in good years imply a flattening genotype-environment-phenotype relationship (Figure 2), the relatively large latent-scale 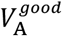 transformed into small phenotypic-scale genotypic variances 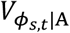 (Figure 5a). The partial evolutionary responses to selection 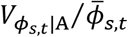 (Figure 5b), and proportional partial responses 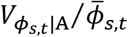 (which entail dividing by values close to one) were consequently similarly small (Figure 5c).

**Figure 5.**
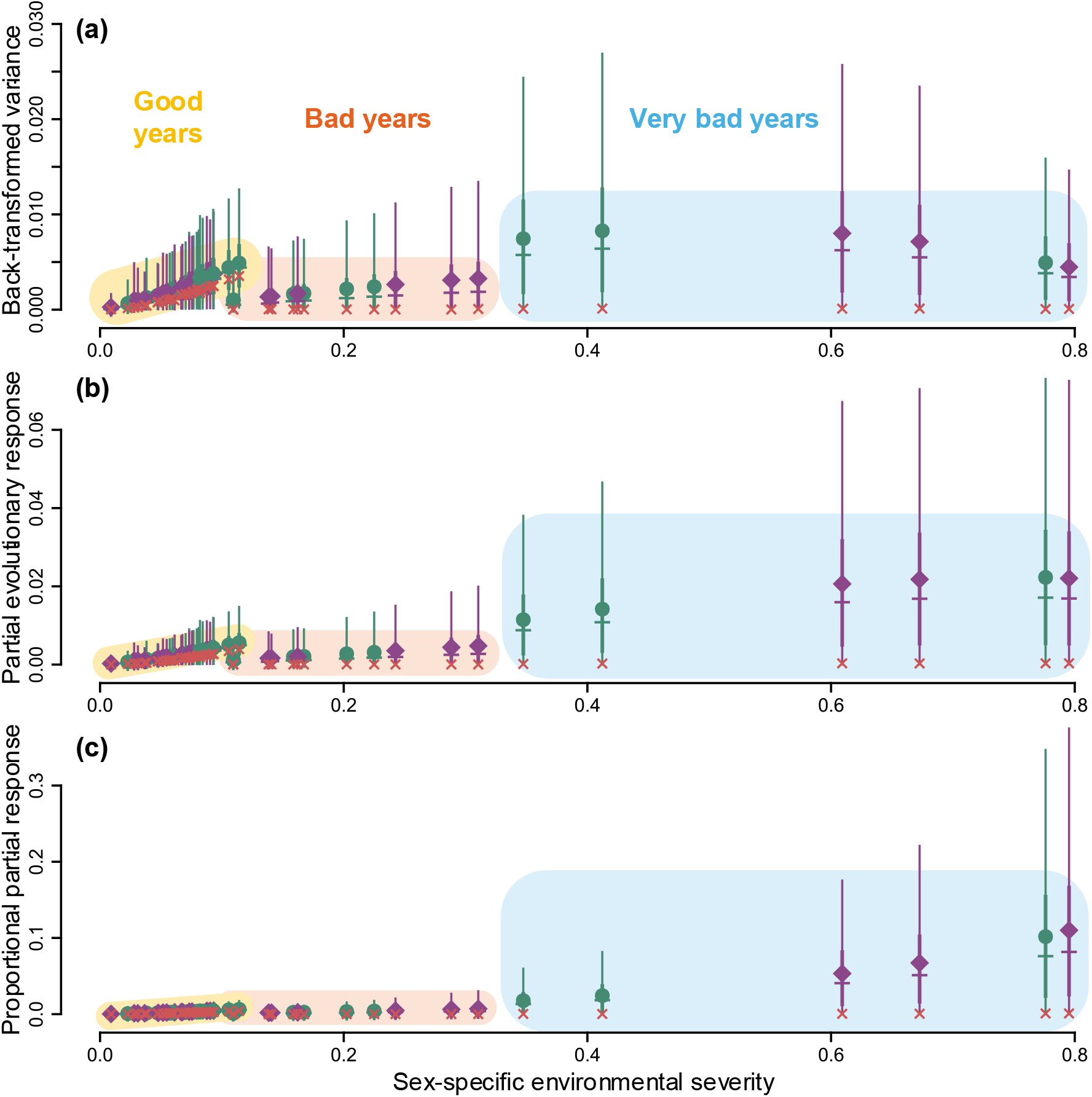
Phenotypic-scale consequences of the additive genetic variance underlying annual adult survival. (a) The sex×year-specific phenotypic-scale genotypic variance in survival 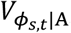, back-transformed given the environment-specific latent-scale additive genetic variance 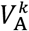 at age 7 years. (b) The partial evolutionary response to selection 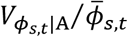 representing the genetic shift in mean survival for the focal sex and annual conditions (and age 7) due strictly to changing allelic frequencies weighted by their initial phenotypic effect. (c) The partial evolutionary response as a proportion of the initial mean survival 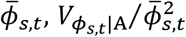. In all panels, environmental severity is defined as 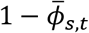; it is therefore sex-specific and differs from the population-level environmental severity (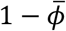; Figure 2) used to define the limits between good, bad and very bad years (delineated here by shaded areas). Green circles and purple diamonds show posterior means of males and females, respectively. Thick and thin error bars show 50% and 95% credible intervals. Horizontal ticks and red crosses show posterior medians and modes. To facilitate readability, uncertainties in environmental severity are not shown (see Appendix S4).

Meanwhile, although the lower survival probabilities in bad and very bad years (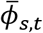 closer to 0.5) imply greater translation of latent variation into phenotypic variation (Figure 2), 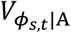 and resulting partial responses to selection had posterior modes on zero (Figure 5). This is expected since the posterior distributions of 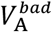 and 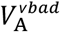 effectively peaked at zero (Figure 4c– d). Yet, uncertainties in 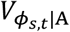 are such that differences with good years are no longer evident and 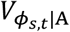 could be greater in very bad years than in the best years (Figure 5a). In turn, since selection is much stronger in severe environments with very low 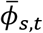 partial evolutionary responses (which entail dividing by values much less than one) could be substantially greater in very bad years than in good and bad years, despite opposing patterns in 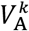 (Figure 5b–c vs Figure 4b-d). Given the posterior modes on zero, these possible differences in the magnitude of adaptation are not strongly supported, but highlight how substantial underlying latent-scale G×E could generate environment-independence, or even reversed G×E, in phenotypic evolutionary responses.

Finally, no major sex differences were apparent in 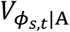 or in the partial evolutionary responses (Figure 5), due to the small sex differences in 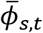 in good years (Figure 3a), and the large uncertainty and posterior modes on zero for 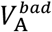 and 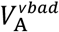. Moreover, large sex differences in 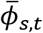 in two out of three very bad years (Figure 3a) placed both sexes at opposite positions of similar steepness on the genotype-environment-phenotype relationship curve (Figure 2), minimising consequent sex differences in phenotypic variances.

## DISCUSSION

Despite its fundamental contribution to adaptive potential, the sensitivity of additive genetic variance (*V*_A_) in survival to differing environmental conditions (including ECEs) has not been quantified in natural populations (Moiron et al., 2022; Baeckens & Donihue, 2025). We detected non-negligible latent-scale *V*_A_ underlying adult annual survival across all observed years in European shags, demonstrating capacity for adaptive micro-evolution through this core fitness component. Yet, partitioning the dataset and analysis by categories of environmental severity (defined by the population mean annual adult survival probability, 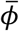) revealed that *V*_A_ was substantial in good years with high 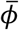, and negligible in bad years with intermediate 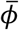. While the estimate of *V*_A_ for very bad years with low 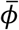 was imprecise, there was no evidence that such conditions released large amounts of previously cryptic genetic variation relative to good conditions, as often postulated (Paaby & Rockman, 2014; Chevin & Hoffmann, 2017). Instead, bad conditions apparently constrained adaptive potential, likely reflecting hard physiological limits independent of genotypes and/or erosion of genetic variation by past selection. Hence, quantitative genetic analyses of population resilience to climate change that ignore environmental severity would underestimate adaptive potential under good conditions and overestimate it under severe conditions such as ECEs, producing biased evolutionary predictions.

These patterns of latent-scale G×E do not, however, directly translate into realised adaptive potential because concurrent environmentally-induced changes in 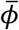 will affect the extent to which underlying variation is manifested as phenotypic variation in survival versus death (Figure 1b; Appendix S1). In shags, good years place the population in a flat region of the sigmoidal genotype-environment-phenotype relationship (Figure 2), meaning that the substantial latent-scale *V*_A_ generates only small phenotypic-scale genotypic variance, *V*_*ϕ*/_*C*_;A_. In contrast, bad and very bad survival years place the population near the steep region (Figure 2), such that very small underlying *V*_A_ could still generate appreciable *V*_*ϕ*/_*C*_;A_ (Figure 5a vs Figure 4b-d). Accordingly, partial evolutionary responses were restricted in good years and might even be exceeded by those in bad and very bad years, because lower survival also implies stronger selection (Figure 5b-c).

However, small partial evolutionary responses can be more consequential than their numerical magnitude may suggest, particularly as a process supporting population persistence in the face of environmental change. For instance, while a partial response of ∼0.003 (the phenotypic-scale posterior median in good years with survival probability ∼0.90; Figure 5b) appears small, it implies a cumulative partial increase of ∼0.1 in survival probability over three decades of similar conditions, representing substantial adaptation. Further, while the partial evolutionary response could be much larger under severe conditions, whether this would imply greater contribution to overall adaptation and population persistence depends on the temporal pattern of environmental severity and associated extinction risk. This contribution will also depend on how adaptive potential interacts with demography. In shags, sex differences in survival probability can sometimes be so substantial in severe environments (Figure 3) that successive bad or very bad years could generate a male-biased adult sex ratio, potentially affecting population fitness and counter-acting any simultaneous adaptation in survival. Sex- and age-specific differences in survival could also imply variation in how much *V*_A_ effectively translates into *V*_*ϕ*/_*C*_;A_ and how strongly selection acts within each sex and age. Our conceptual framework and estimates thereby formalise how genetic, environmental, and phenotypic variances, together with demography, jointly shape the complex effects of ECEs on eco-evolutionary responses.

These findings invite revisiting the few previous estimates of *V*_A_ underlying adult annual survival in wild animals, which were all interpreted as negligible. Estimates in song sparrows (*Melospiza melodia*: Wolak et al., 2018, who quantified sex-specific *V*_A_) and common terns (*Sterna hirundo*: Moiron et al., 2022) were indeed indistinguishable from a null expectation. Meanwhile, estimates in blue tits (*Cyanistes caeruleus*: Papaï x et al., 2010) and Soay sheep (*Ovis aries*: Stoffel et al., 2017, who quantified *V*_A_ across juveniles and adults combined) were just small. Yet, small numerical values are broadly expected on the underlying scale (Acker & Reid, 2025), and phenotypic implications could still be considerable (Appendix S1). Further, none of these previous studies quantified impacts of (sex- and/or age-specific) environmental severity on *V*_A_, even though all reported notable among-year variation in mean survival. The overall estimates might consequently have obscured G×E (Fry, 1992), as illustrated by our current results (Figure 4). More generally, wild population studies that estimated *V*_A_ in any measure of fitness, including lifespan-based measures (i.e. longevity or lifetime reproductive success), have almost all ignored impacts of environmental severity (Hendry et al., 2018; Kulbaba et al., 2019; Bonnet et al., 2022; Moiron et al., 2022; Dumas et al., 2026; but see McFarlane et al., 2014). Targeted (re)analyses of such datasets are now required to reveal any typical patterns of how environmental severity shapes adaptive potential in nature.

Progress in quantifying adaptive potential will also require tighter focus on temporal scales matching the pace of environmental variation. This could imply annual survival and/or reproductive success (Sæther & Engen, 2015), or even shorter seasonal timeframes (Fortuna et al., 2024). Here, our rationale for how environmental severity may reshape adaptive potential across underlying and phenotypic scales (Figure 1), and associated mathematical derivations grounded in the Price equation (Appendix S1; Price, 1970), also apply to other dichotomous fitness outcomes (e.g. breeding vs not, successful vs failed reproduction). Analogous rationales could apply to any form of absolute fitness, which is fundamentally expressed via successive discrete outcomes of survival and reproduction countable in units of individuals (Arnold & Wade, 1984; Sæther & Engen, 2015). This necessarily implies non-linear genotype-environment-phenotype relationships, yielding non-Gaussian distributions and non-independence between fitness means and variances (Shaw et al., 2008; de Villemereuil, 2018; Bonnet et al., 2019). Further, broadly sigmoidal relationships (e.g. Figure 2) are probably reasonable postulates for most definitions of absolute fitness, including annual or lifetime reproductive success, where values are bounded between 0 and some upper limit set by organismal constraints that cannot be readily overcome through plasticity or allelic substitutions. Hence, our formal framework (Figure 1; Appendix S1) provides foundations for dissecting how environmental changes, including ECEs, shape adaptive potential through any fitness component.

Specific metrics tailored for appropriately measuring and comparing adaptive potential across environmental conditions, or contrasted regimes of environmental variation, are then required to enable robust predictions of population resilience to climate change and escalating ECEs. Here, treating the proportional partial evolutionary response as a standardised metric of adaptive potential, as often advocated for fitness (Hendry et al., 2018; Bonnet et al., 2022), almost systematically implies an exponential increase with increasing environmental severity (and resulting lower mean survival or reproduction; Figures 1d, 5c; Appendices S1, S4). Comparing evolutionary implications across environments through proportional responses is consequently less straightforward than comparing across systems in similar environments. Indeed, because fitness is measured as countable outcomes on absolute scales (sensu measurement theory), its information content is hardly preserved by mean-standardisation (Hansen & Houle, 2008; Houle et al., 2011). More generally, under escalating ECEs, the pace and amplitude of environmental fluctuations can become far greater than under relatively stable conditions for which standard evolutionary theory and metrics were primarily developed (Sæther & Engen, 2015; Chevin & Hoffmann, 2017). To address this challenge, standardised quantifications and comparisons of adaptive potential will require coupling scenarios of temporal distributions of environmental severity with predictive models of population adaptation (Chevin et al., 2010; Baeckens & Donihue, 2025). As we have demonstrated, such models will need to incorporate environment-specific quantitative genetic estimates underlying fitness variation and their non-linear phenotypic transforms into demographic projections.

## DATA AND CODE AVAILABILITY

All data used in this study, the code, and primary model outputs (full posterior samples with detailed numerical summaries) are publicly available from the Zenodo repository via https://doi.org/10.5281/zenodo.22274665 (Acker et al., 2026).

## AUTHOR CONTRIBUTIONS

Conceptualization: P.A. (lead), J.M.R.; Formal analysis, Methodology, Software, Validation, Visualization, Writing – original draft: P.A.; Data curation: S.J.B. (lead), P.A., R.F.; Funding acquisition, Supervision: J.M.R.; Investigation: S.W. (lead), F.D. (lead), M.A.N., S.J.B., M.P.H., C.G., R.E.D., E.B.; Project administration: J.M.R., P.A.; Writing – review & editing: P.A. (lead), J.M.R. (co-lead), F.D., R.F., S.J.B., R.E.D., S.W.

## ACKNOWLEDGEMENTS

We thank everyone who contributed to long-term field data collection, particularly Jenny Bull, Linda Wilson, Maggie Hall, Tom Reed, Morten Frederiksen, Hanna Granroth-Wilding, Katherine Herborn, Olivia Hicks, Hannah Ravenswater, Fiona Greco, Fran Ruiz-Raya, Josie Hewitt, and Erin Taylor. We thank NatureScot for support and allowing work on the Isle of May National Nature Reserve, and Isle of May Bird Observatory Trust for supporting the long-term ringing of shags. This work was partially funded by the European Union. Views and opinions expressed are however those of the author(s) only and do not necessarily reflect those of the European Union or the European Research Council Executive Agency. Neither the European Union nor the granting authority can be held responsible for them.

