## Supplementary Information: Appendix S1 to S5 for "Extreme climatic events shape adaptive potential by altering genetic variance for adult survival in a wild population"

#### Contents:

**Appendix S1: Details of the conceptual framework (p. 2)**

- S1.1 Back-transformation of latent-scale $V_{\mathrm{A}}$ onto the phenotypic scale of survival (p. 2)
- S1.2 Formulating evolutionary implications of $V_{\mathrm{A}}$ underlying survival (p. 4)
- S1.3 Application to previous results in other study systems (p. 13)
- S1.4 Hypothetical relationships linking adaptive potential to environmental severity (p. 14)

**Appendix S2: Details of the datasets (p. 18)**

- S2.1 Study system and resighting data (p. 18)
- S2.2 Structure of the pedigree data (p. 22)

**Appendix S3: Details of the statistical modelling (p. 25)**

- S3.1 Implementation of the capture-recapture model analyses (p. 25)
- S3.2 Implementation and key assumptions of the animal model (p. 26)
- S3.3 Prior specifications (p. 27)
- S3.4 Additional model versions and checks (p. 28)
- S3.5 Simulations evaluating our capability to detect $V_{\mathrm{A}}$ (p. 34)
- S3.6 Posterior sampling and derived calculations (p. 36)

**Appendix S4: Additional details of the results (p. 38)**

- S4.1 Sex- and age-specific survival (p. 38)
- S4.2 Detection probabilities (p. 40)
- S4.3 Probit regression parameters (p. 40)
- S4.4 Phenotypic-scale consequences (p. 41)
- S4.5 Alternative quantitative genetic models (p. 44)

**Appendix S5: References cited in Supporting Information (p. 52)**

### Appendix S1

### Details of the conceptual framework

#### S1.1 Back-transformation of latent-scale $V_{\mathrm{A}}$ onto the phenotypic scale of survival

In general, $V_{\mathrm{A}}$ provides a compelling measure of adaptive potential because it gives the expected evolutionary response per unit selection gradient when the genotype-phenotype relationship is linear or approximately so (as typically postulated in standard quantitative genetic analyses of continuously distributed traits; Lande & Arnold, 1983; Houle, 1992; Hansen & Pélabon, 2021). In dichotomous traits, this property holds for underlying $V_{\mathrm{A}}$ defined on the unbounded continuously distributed latent scale. Yet, due to the sigmoidal genotype-environment-phenotype relationship, part of the genetic variation that is additive on the latent scale systematically becomes non-additive on the dichotomous phenotypic scale (Dempster & Lerner 1950; de Villemereuil et al., 2016; Acker & Reid, 2025). Consequently, any $V_{\mathrm{A}}$ directly defined on the phenotypic scale (as the component of phenotypic variance due to genetic effects that are strictly additive on phenotype) would not encompass all the variation stemming from the underlying additive genetic effects, and hence would not fully represent the adaptive potential given latent-scale $V_{\mathrm{A}}$. The full expressed variance stemming from effects that can lead to evolutionary changes in response to selection thus corresponds to the (partly non-additive) among-genotype variance arising on the phenotypic scale from variation in underlying latent-scale additive genetic effects.

Considering the current focal dichotomous phenotype of survival versus death, by definition, the variance in phenotype $z$ (taking values 0 for death and 1 for survival) arising from variation in the latent-scale additive genetic effects $a$ (de Villemereuil et al., 2016; Acker & Reid, 2025) is $V_{\phi|\mathrm{A}} = Var\big(E(z|a)\big)$. Here, since the expectation of an individual's survival outcome is its survival probability $\phi$, $V_{\phi|\mathrm{A}}$ is also the variance in $\phi$ given $a$. Moreover, by the law of total expectation, $E\big(E(z|a)\big) = E(z) = \bar{\phi}$, where $\bar{\phi}$ denotes the population mean survival probability. Under the assumption that the latent-scale additive genetic effects are distributed such that $a \sim \mathcal{N}(0, V_{\mathrm{A}})$ (as is standard in animal models, and applied in all our analyses), $V_{\phi|\mathrm{A}}$ is then given by (de Villemereuil et al., 2016; Acker & Reid, 2025):

$$V_{\phi|\mathrm{A}} = \int_{-\infty}^{+\infty} (E(z|a) - \bar{\phi})^2 f_{\mathcal{N}}\left(\frac{a}{\sqrt{V_A}}\right) d_a, \tag{S1}$$

where $f_{\mathcal{N}}$ is the probability density function of the standard normal distribution.

As illustrated in main text Figure 2, $E(z|a)$ and $\bar{\phi}$, and thus $V_{\phi|\mathrm{A}}$, depend on the latent mean (and hence on any intercept or linear predictor envisaged on the latent scale). To see this, consider a baseline general case given a simple quantitative genetic animal model with an overall intercept $\eta$ and a breeding value $a$ defined on the latent scale using a probit link:

$$\mathrm{probit}(\phi) = \eta + a. \tag{S2}$$

Note that any further variation is included within the random residual around the expectation for survival (i.e. around probit($\phi$), given $z{\sim}Bernoulli(\phi)$). This is enacted here through the probit link which implies a threshold trait model whereby phenotypic expression is underlaid by a latent variable $\ell$ (the 'liability'; Falconer 1965) defined as $\ell = \eta + a + \varepsilon$, where $\varepsilon{\sim}\mathcal{N}(0,1)$ (representing the residual variation), and such that $z = 1$ if $\ell > 0$ or $z = 0$ otherwise. Hence, a binary GLMM using a probit link formulates a linear regression for $E(\ell)$ on a latent scale where the unit is the liability residual standard deviation (Falconer & Mackay 1996; Agresti, 2002; Acker & Reid, 2025). Moreover, given the probit link, $\bar{\phi}$ and $E(z|a)$ have simple expressions (Acker & Reid, 2025):

$$\bar{\phi} = F_{\mathcal{N}}\left(\frac{\eta}{\sqrt{V_A + 1}}\right), \tag{S3}$$

$$E(z|a) = F_{\mathcal{N}}(\eta + a), \tag{S4}$$

and Equation (S1) has the following analytical solution (Acker & Reid, 2025):

$$V_{\phi|\mathrm{A}} = \bar{\phi}(1 - \bar{\phi}) - 2T\left(\frac{\eta}{\sqrt{V_A + 1}}, \sqrt{\frac{1}{2V_A + 1}}\right), \tag{S5}$$

where $F_{\mathcal{N}}$ is the cumulative distribution function of the standard normal distribution (also known as the inverse probit function) and $T(v, w)$ is Owen's T function (Owen, 1956).

In the case of our overall quantitative genetic animal model defined for adult survival in our study population of shags (main text Equation 2), $V_{\phi|\mathrm{A}}$ thus becomes specific to each sex, year, and age (indexed as $s$, $t$, and $x$, see main text). This is made explicit by rewriting Equation (S1) as:

$$V_{\phi_{s,t,x}|\mathrm{A}} = \int_{-\infty}^{+\infty} \left(E(z|s,t,x,a) - \bar{\phi}_{s,t,x}\right)^2 f_{\mathcal{N}}\left(\frac{a}{\sqrt{V_A}}\right) d_a, \tag{S6}$$

where $\bar{\phi}_{s,t,x}$ is the survival expectation across individuals for the focal sex, year and age. We then have (Acker & Reid, 2025):

$$\bar{\phi}_{s,t,x} = \mathrm{probit}^{-1}\left(\frac{\eta(s,t,x)}{\sqrt{V_A + V_B + 1}}\right), \tag{S7}$$

$$E(z|s,t,x,a) = \mathrm{probit}^{-1}\left(\frac{\eta(s,t,x) + a}{\sqrt{V_B + 1}}\right), \tag{S8}$$

where $\eta(s,t,x) = \mu + \gamma_{s,t} + \beta_1 x + \beta_2 x^2$, and (as in Equations (1-2) of the main text) $\gamma_{s,t}$ is the year×sex effect, $\beta_1$ and $\beta_2$ are linear and quadratic effects of age and $V_B$ is the permanent individual variance. Equation (S6) is then solved as (Acker & Reid, 2025):

$$V_{\phi_{s,t,x}|\mathrm{A}} = \bar{\phi}_{s,t,x}\left(1 - \bar{\phi}_{s,t,x}\right) - 2T\left(\frac{\eta(s,t,x)}{\sqrt{V_A + V_B + 1}}, \sqrt{\frac{V_B + 1}{2V_A + V_B + 1}}\right). \tag{S9}$$

#### S1.2 Formulating evolutionary implications of $V_\mathrm{A}$ underlying survival

*Deriving the partial evolutionary response in mean survival as $V_{\phi|A}/\bar{\phi}$*

Because survival is a direct component of absolute fitness, its phenotypic expression through a given timestep effectively constitutes an episode of directional selection. Here, the selection differential, $S$, defined as the difference in mean phenotype across individuals right after selection compared to all individuals initially alive (Lush, 1937), is always $1 - \bar{\phi}$ (since, by definition, survivors have mean phenotypic survival of 1). This result is retrieved by applying the Robertson-Price identity which expresses $S$ as the covariance between relative fitness and phenotype, $Cov(w, z)$ (Robertson 1966; Price 1970). Here, relative fitness for any individual is obtained by dividing its survival by mean survival, giving $w = z/\bar{\phi}$, and thus:

$$S = Cov(z/\bar{\phi}, z)$$
$$= Var(z)/\bar{\phi}. \quad \text{(S10)}$$

And since $z$ is a Bernoulli variable with mean $\bar{\phi}$, so that $Var(z) = \bar{\phi}(1-\bar{\phi})$, Equation (S10) gives:

$$S = 1 - \bar{\phi}. \quad \text{(S11)}$$

Moreover, the selection gradient, i.e. the selection differential divided by the phenotypic variance (Lande & Arnold, 1983), $\beta$, is thus always $1/\bar{\phi}$ when the focal trait is survival.

Accordingly, the immediate (within-generation) change in $\bar{\phi}$ due strictly to direct selection on the underlying additive genetic effects $a$ (given the initial survival values) is then $V_{\phi|\mathrm{A}}/\bar{\phi}$. That is, the product of the phenotypic-scale genotypic variance and the selection gradient. This result can also be obtained by directly calculating the selection differential at the level of the expected phenotype given any underlying additive genetic value $E(z|a)$ (i.e. for the phenotypic-scale genotypic values for survival), $S_{\phi|A}$:

$$S_{\phi|A} = Cov\left(z/\bar{\phi}, E(z|a)\right)$$
$$= Cov(z, E(z|a))/\bar{\phi}. \quad \text{(S12)}$$

Since the covariance between a random variable $Y$ and its conditional expectation given another variable $X$ equals the variance of that conditional expectation, i.e. $Var\big(E(Y|X)\big)$ (see Box S1):

$$S_{\phi|A} = Var\big(E(z|a)\big)/\bar{\phi}$$
$$= V_{\phi|\mathrm{A}}/\bar{\phi}. \quad \text{(S13)}$$

Accordingly, $V_{\phi|\mathrm{A}}/\bar{\phi}$ quantifies how much $\bar{\phi}$ increases due to the underlying change in allele frequencies, given the phenotypic effects of the alleles under the focal pre-selection conditions. Placed in the context of the Price equation (Price, 1970, 1972), $V_{\phi|\mathrm{A}}/\bar{\phi}$ thus represents the *partial evolutionary change* in survival (Ewens, 1989; Frank, 2012; Walsh & Lynch, 2018). The total cross-generation change expected from the survival (and hence selection) episode includes any possible further shift due to how post-selection conditions alter the allelic effects on phenotypes expressed after reproduction (given functional epistasis, dominance, and G×E). Here it is important to note that the post-selection conditions encompass the focal selection-driven changes in allele frequencies, but also possibly any further changes due to mutation and gene flow or other episodes of selection, and any other concurrent environmental change.

To clarify this, consider the general Price equation that decomposes the total cross-generation change $\Delta_{\bar{z}}$ into a term representing selection on the parents and a second term representing post-selection transmission to their offspring (Frank, 2012; Walsh & Lynch, 2018):

$$\Delta_{\bar{z}} = Cov(w, z) + E\left(w\bar{\delta}_z\right). \tag{S14}$$

Here, $w$ and $z$ refer strictly to the parental generation, but $\bar{\delta}_z = z' - z$ is the average difference in phenotype from a parent to its offspring (i.e. in our analyses, $z'$ is the mean adult survival of the offspring from the focal parent). Given our quantitative genetic objective, we formulate $z$ as the sum of a genotypic component (underlaid by the latent scale additive genetic effect $a$, transmissible to the offspring) and a residual, i.e. $z = E(z|a) + \epsilon$. We then have $z' = E(z'|a)$ by definition, and thus:

$$Cov(w, z) = Cov\left(w, E(z|a)\right) + Cov(w, \epsilon), \tag{S15}$$

$$E\left(w\bar{\delta}_z\right) = E\left(w(E(z'|a) - E(z|a)\right) - Cov(w, \epsilon). \tag{S16}$$

Substituting Equations (S15-S16) into Equation (S14), recalling that $w = z/\bar{\phi}$ and $Cov\left(z, E(z|a)\right) = V_{\phi|\mathrm{A}}$ (Equations S12-S13), and letting $T_{\phi|A} = E\left(w\left(E(z'|a) - E(z|a)\right)\right)$, we can rewrite the Price equation to express $\Delta_{\bar{z}}$ conditional on underlying additive genetic variation as:

$$\Delta_{\bar{z}} = V_{\phi|\mathrm{A}}/\bar{\phi} + T_{\phi|A}. \tag{S17}$$

Equation (S17) expresses the total cross-generation change in survival as the partial evolutionary change due to selection on the breeding values (i.e. the heritable genetic effects) plus the partial change due to transmission of those breeding values from selected parents to their offspring.

Note that $V_{\phi|\mathrm{A}}/\bar{\phi}$ is analogous to the additive genetic variance in relative fitness, $V_A(w)$, which is typically used to quantify adaptative potential (e.g. Kulbaba et al., 2019; Bonnet et al., 2022) based on arguments that assume a linear genotype-phenotype relationship for fitness (Ewens, 1989; Walsh & Lynch, 2018). The key distinction is that $V_{\phi|\mathrm{A}}/\bar{\phi}$ accounts for how adaptive evolution occurring through survival selection is shaped by the inherently non-linear expression of underlying heritable genetic variation in survival.

**Box S1: *Proof of the identity $Cov(X, E(X|Y)) = Var(E(X|Y))$***

To prove the covariance identity for conditional expectation, we unpack the definition of the covariance between X and $E(X|Y)$ into the form of the expectation of the product minus the product of the expectations:

$$Cov(X, E(X|Y)) = E(XE(X|Y)) - E(X)E(E(X|Y)). \quad \text{(S18)}$$

Then, since the law of total expectation states $E(X) = E(E(X|Y))$, we can rewrite the second term of the right-hand side of Equation (S18) as:

$$E(X)E(E(X|Y)) = E(E(X|Y))^2. \quad \text{(S19)}$$

Further, still given the law of total expectation, we can rewrite the first term of the right-hand side of Equation (S18) as:

$$E(XE(X|Y)) = E(E(XE(X|Y)|Y)). \quad \text{(S20)}$$

Here, within the outer conditional expectation, $E(X|Y)$ depends only on Y, so it can be treated as a constant and pulled outside, giving:

$$\begin{aligned} E(XE(X|Y)) &= E(E(X|Y)E(X|Y)) \\ &= E(E(X|Y)^2). \end{aligned} \quad \text{(S21)}$$

Therefore, given (S19) and (S21), we can simplify (S18) to:

$$\begin{aligned} Cov(X, E(X|Y)) &= E(E(X|Y)^2) - E(E(X|Y))^2 \\ &= Var(E(X|Y)). \end{aligned} \quad \text{(S22)}$$

*Evaluating $V_{\phi|A}/\bar{\phi}$ as a measure of adaptive potential*

To further clarify the interpretation of $V_{\phi|A}/\bar{\phi}$ as a measure of adaptive potential, we need to examine the remaining component of the evolutionary response (Equation S17), namely the partial change due to cross-generation transmission, $T_{\phi|A}$. We can envisage the focal survival (and hence selection) episode as the only driver of cross-generation change, assuming random mating and no further effective environmental change (including given negligible gene flow and mutations). The transmission term would be zero given a linear genotype-phenotype relationship (Frank, 2012; Walsh & Lynch, 2018). However, for survival this term is not zero because the non-linear genotype-phenotype relationship (Figure 2 of main text) distorts the allelic effects on phenotype as selection changes the mean and variance of $a$. Nonetheless, $T_{\phi|A}$ is still reduced to its minimum potential influence, which is typically negligible, corresponding to where the adaptive potential represented by $V_{\phi|\mathrm{A}}/\bar{\phi}$ can be effectively realised to its maximal extent.

To show this, we derived an analytical expression for $T_{\phi|A}$ (details below; Equations S23-S38). We thereby calculated exact values of $\Delta_{\bar{z}}$ for the baseline general case where survival is formulated as an intercept plus a breeding value on the probit scale (Equation S2), by substituting Equations (S4) and (S38) into (S17). We implemented this solution to calculate $\Delta_{\bar{z}}$ for a full gradient of environmental severity (i.e. $\bar{\phi}$ spanning 0–1) and $V_A$ values of 0.075, 0.15, 0.25, 0.5, 1, and 2, yielding latent scale heritabilities of 0.07, 0.13, 0.2, 0.33, 0.5, and 0.67 (noting that, on the probit scale, $h^2 = V_A/(V_A + 1)$). Here, a nice property is that emerging results on the phenotypic scale are generalisable to any underlying animal model formulation given the same $h^2$ value. We then also calculated $V_{\phi|\mathrm{A}}/\bar{\phi}$ to evaluate its performance in predicting total evolution. We further calculated $V_A(w)$ to evaluate discrepancies with this common metric of adaptive potential. The results (Figure S1) demonstrate that $V_{\phi|\mathrm{A}}/\bar{\phi}$ provides a very close approximation of $\Delta_{\bar{z}}$ across the full range of environmental severity and for all simulated $V_A$ values (though a slight bias becomes apparent for the unrealistic scenario of $h^2 = 0.67$). Meanwhile, $V_A(w)$ also provides a very close approximation given environmental severities <0.6, and more widely given small $h^2$. However, it provides a downwardly-biased approximation of $\Delta_{\bar{z}}$ for environmental severities >0.6 (as is relevant in our context of a typically long-lived species experiencing high annual mortality due to extreme climatic events, or for shorter-lived where mean survival is always low), and the bias increases substantially as $h^2$ increases (Figure S1).

While this highlights the key relevance of using $V_{\phi|\mathrm{A}}/\bar{\phi}$ to quantify phenotypic-scale evolutionary implications of latent-scale $V_{\mathrm{A}}$, it is important to remember that $V_{\phi|\mathrm{A}}/\bar{\phi}$ represents an evolutionary *potential*. It provides a precise measure of expected evolution under assumptions constituting an idealised set of conditions (random mating, no inbreeding, no gene flow, no mutation, no environmental change). Because these assumptions are likely to be violated in nature, the partial evolutionary response measured by $V_{\phi|\mathrm{A}}/\bar{\phi}$ is thus to be interpreted as a useful conceptual metric of adaptive potential but not an accurate prediction of realised evolutionary outcomes. Following this spirit of quantitative abstraction, we can envisage sex-by-year-by-age specific $V_{\phi|\mathrm{A}}/\bar{\phi}$ (given Equations S7 and S9) and hence evolutionary potentials, even though evolution inherently results from an interplay across sexes, ages, and years (shaped by genetic correlations, the dynamic demographic structure, and patterns of stochastic environmental variation; see Introduction and Discussion of main text). In reality, this complex interplay, and any deviations from idealised conditions, will effectively act to distort transmission of (sex-by-year-by-age specific) phenotypic effects, and hence the evolutionary response.

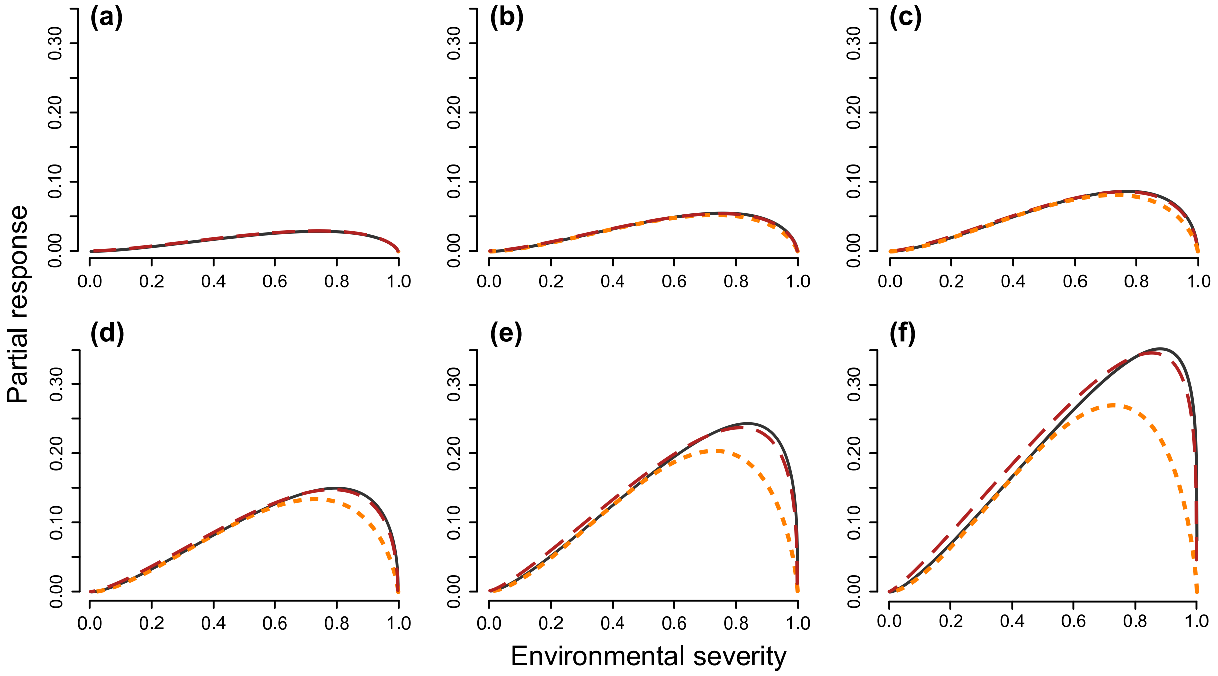


**Figure S1.** Expected change in mean survival ($\bar{\phi}$) in response to selection through survival, as a function of environmental severity ($1 - \bar{\phi}$), derived for latent-scale heritabilities ($h^2$) of (a) 0.07, (b) 0.13, (c) 0.2, (d) 0.33, (e) 0.5, and (f) 0.67. On each panel, the black plain line shows the total evolutionary response, $\Delta_{\bar{z}}$, the red dashed line shows the partial evolutionary response, $V_{\phi|\mathrm{A}}/\bar{\phi}$, and the orange dotted line shows the additive genetic variance in relative survival, $V_A(w)$.

*Calculation of the cross-generation transmission term*

To solve $T_{\phi|A}$, we can start by slightly expanding it as:

$$T_{\phi|A} = E\left(w\big(E(z'|a)\big)\right) - E\left(w\big(E(z|a)\big)\right), \tag{S23}$$

hence making explicit that it is the difference between the overall expectation of the mean offspring phenotype and the overall expectation of the parent phenotype evaluated given the post-selection distribution of $a$. Indeed, $w$ acts here as a probability weight shifting the distribution of breeding values such that, for any function $g$, $E\big(wg(a)\big)$ becomes $E\big(g(a^*)\big)$, with $a^* \sim \mathcal{N}(\bar{a}^*, V_A^*)$ (where $\bar{a}^*$ and $V_A^*$ are mean and variance of the breeding values among surviving parents; readily calculable using the analytical solutions provided in Box S2).

**Box S2:** ***Formulae for the mean and variance of breeding values among selected parents***

*Mean breeding value of selected parents*

Because the mean breeding value before selection is 0, the mean breeding value of selected parents is directly given by the selection differential for the breeding values, i.e. $\bar{a}^* = Cov(w, a)$. Noting again that $w = z/\bar{\phi}$ and given the law of total variance, we thus have:

$$\bar{a}^* = \frac{Cov(E(z|a), a)}{\bar{\phi}}. \tag{S24}$$

Since $a$ is normally distributed, Equation (S24) can be solved by applying Stein's (1973) Lemma. Under the baseline general case defined by Equation (S2), we have $E(z|a) = F_{\mathcal{N}}(\eta + a)$ (Equation S4). Noting that the first derivative of $F_{\mathcal{N}}$ is $f_{\mathcal{N}}$, this gives:

$$\bar{a}^* = V_A \frac{E\big(f_{\mathcal{N}}(\eta + a)\big)}{\bar{\phi}}. \tag{S25}$$

Following the theorem for the expectation of a function of a continuous random variable (Casella & Berger, 2002; Ross, 2014), we have:

$$E\big(f_{\mathcal{N}}(\eta + a)\big) = \int_{-\infty}^{+\infty} f_{\mathcal{N}}(\eta + a) \frac{1}{\sqrt{Va}} f_{\mathcal{N}}(a) da. \tag{S26}$$

Solving the integral equation (S26) (see Owen, 1980) and substituting into Equation (S25) gives the following analytical expression:

$$\bar{a}^* = \frac{V_A}{\sqrt{V_A+1}} \frac{f_{\mathcal{N}}\left(\frac{\eta}{\sqrt{V_A+1}}\right)}{\bar{\phi}}. \tag{S27}$$

*Variance in breeding values of selected parents*

By definition, the variance in breeding values of selected parents, $V_A^*$ is $E(\bar{a}^{*2}) - \bar{a}^{*2}$. As per the same logic used to explain Equation (S24) above, we thus have:

$$\begin{aligned} V_A^* &= E(wa^2) - \bar{a}^{*2} \\ &= \frac{E(za^2)}{\bar{\phi}} - \bar{a}^{*2} \\ &= \frac{E\big(a^2 E(z|a)\big)}{\bar{\phi}} - \bar{a}^{*2}. \end{aligned} \tag{S28}$$

Further, recalling that $E(a) = 0$, Equation (S28) simplifies to:

$$V_A^* = \frac{Cov\big(a, aE(z|a)\big)}{\bar{\phi}} - \bar{a}^{*2}. \tag{S29}$$

We can again use Stein's Lemma to solve Equation (S29). Under the baseline general case defined by Equation (S2), the derivative of $aE(z|a)$ is $F_{\mathcal{N}}(\eta + a) + af_{\mathcal{N}}(\eta + a)$, and thus:

$$V_A^* = V_A \frac{E\big(F_{\mathcal{N}}(\eta + a)\big) + E\big(af_{\mathcal{N}}(\eta + a)\big)}{\bar{\phi}} - \bar{a}^{*2}. \tag{S30}$$

Here, we note that $F_{\mathcal{N}}(\eta + a) = E(z|a)$ (Equation S4), and thus $E\big(F_{\mathcal{N}}(\eta + a)\big) = E(z) = \bar{\phi}$, by the law of total expectation, which gives:

$$V_A^* = V_A \left(1 + \frac{E\big(af_{\mathcal{N}}(\eta + a)\big)}{\bar{\phi}}\right) - \bar{a}^{*2}. \tag{S31}$$

And again given the definition of the expected value of a function of a continuous random variable (Casella & Berger, 2002; Ross, 2014), we have:

$$E\big(af_{\mathcal{N}}(\eta + a)\big) = \int_{-\infty}^{+\infty} af_{\mathcal{N}}(\eta + a) \frac{1}{\sqrt{Va}} f_{\mathcal{N}}(a). \tag{S32}$$

Solving this integral equation (see Owen, 1980) then gives:

$$E\big(af_{\mathcal{N}}(\eta + a)\big) = \frac{{V_A}^2}{V_A + 1} \frac{\eta}{\sqrt{V_A+1}} \frac{f_{\mathcal{N}}\left(\frac{\eta}{\sqrt{V_A+1}}\right)}{\bar{\phi}}. \tag{S33}$$

And finally, substituting Equation (S33) into Equation (S31) and noting that $V_A/(V_A+1) = h^2$ yields the following analytical expression:

$$V_A^* = V_A\left(1 - h^2\left(\frac{\eta}{\sqrt{V_A+1}}\frac{f_{\mathcal{N}}\left(\frac{\eta}{\sqrt{V_A+1}}\right)}{\bar{\phi}} + \frac{f_{\mathcal{N}}\left(\frac{\eta}{\sqrt{V_A+1}}\right)^2}{\bar{\phi}^2}\right)\right). \quad \text{(S34)}$$

Accordingly, we can express $T_{\phi|A}$ as a function of the post-selection distribution of $a$:

$$T_{\phi|A} = E\big(E(z'|a^*)\big) - E\big(E(z|a^*)\big). \quad \text{(S35)}$$

Under the infinitesimal model with random mating, an offspring's breeding value is a normally distributed random deviation from the average of the two parental breeding values, due to Mendelian sampling. Hence, the focal selected parent with breeding value $a^*$ has offspring with breeding values distributed given expectation $E(a') = (a^* + \bar{a}^*)/2$ and variance $V_A' = V_A^*/4 + V_A/2$ (where $\bar{a}^*$and $V_A^*/4$ are the expectation and variance for the breeding value of the random parent, and $V_A/2$ is the segregation variance; Barton et al., 2017). Returning to the baseline general case defined in Equation (S2), following Equations (S3-S4) we then have:

$$E(z'|a^*) = F_{\mathcal{N}}\left(\frac{\eta + \frac{\bar{a}^*}{2} + \frac{a^*}{2}}{\sqrt{\frac{V_A^*}{4} + \frac{V_A}{2} + 1}}\right), \quad \text{(S36)}$$

$$E(z|a^*) = F_{\mathcal{N}}(\eta + a^*). \quad \text{(S37)}$$

Finally, under the approximation that the selection-induced skew in parental breeding values is effectively negligible, given the theorem for the expectation of a function of a continuous random variable (Casella & Berger, 2002; Ross, 2014) and well-known normal integral identities (Owen, 1980), averaging over $a^*$ (Equations S36-S37) and substituting into Equation (S35) gives:

$$T_{\phi|A} = F_{\mathcal{N}}\left(\frac{\eta + \bar{a}^*}{\sqrt{\frac{V_A^*}{2} + \frac{V_A}{2} + 1}}\right) - F_{\mathcal{N}}\left(\frac{\eta + \bar{a}^*}{\sqrt{V_A^* + 1}}\right). \quad \text{(S38)}$$

##### S1.3 Application to previous results in other study systems

In addition to calculating $V_{\phi|A}/\bar{\phi}$ for our own model results (using Equation S9; see main text and Appendix S3), the general results of Equations S3 and S5 allow deriving $V_{\phi|A}/\bar{\phi}$, and hence expected partial evolutionary responses, for any estimate of $V_A$ underlying survival obtained from a binary generalised animal model. This only requires parameter estimates from the original animal model used to obtain the focal $V_A$, followed by rescaling (where necessary) to match the baseline general model formulated in Equation (S2), as detailed in Box S3 below.

As mentioned in the main text, only two studies of wild vertebrates have reported non-zero $V_A$ underlying survival (blue tits: Papaïx et al., 2010; Soay sheep: Stoffel et al., 2017). Using estimates from the blue tit study, after rescaling conditional on average year conditions (Box S3), we get $\bar{\phi} = 0.563$ and $V_{\phi|A} = 0.015$, yielding the partial response $V_{\phi|A}/\bar{\phi} = 0.023$ (and proportional response $V_{\phi|A}/\bar{\phi}^2 = 0.049$). Over 10 iterations of the same selection episode under the same conditions, this would give a cumulative partial evolutionary increase in mean survival of 0.23, which is certainly substantial (and seems unrealistic). Using estimates from the Soay sheep study, after rescaling conditional on no inbreeding, no twin, and average birth year and capture year conditions in midlife age class, we get $\bar{\phi} = 0.87$ in females, $\bar{\phi} = 0.84$ in males, and $V_{\phi|A} = 0.001$ in both sexes, yielding $V_{\phi|A}/\bar{\phi} = 0.001$ in females and $V_{\phi|A}/\bar{\phi} = 0.002$ in females (Box S3). These values would represent a cumulative partial increase in mean survival of ~0.01 or 0.02 after 10 iterations of the same selection episode, which is less notable than in blue tits yet still non-trivial.

**Box S3:** ***Rescaling published animal model estimates to calculate $\bar{\phi}$ and $V_{\phi|A}$***

*Example from the blue tit study of Papaïx et al. (2010)*

In blue tits, Papaïx et al. (2010) fitted a probit model that estimated an intercept ($\eta^{\#} = 0.175$, where '#' denotes their original model), plus a breeding value ($a^{\#}$, with variance ${V_A}^{\#} = 0.122$), a permanent individual effect ($b^{\#}$, with variance ${V_B}^{\#} = 0.105$), and a year effect. Therefore, reformulating their model into the baseline general model (Equation S2), conditional on average year conditions, gives $\eta + a = (\eta^{\#} + a^{\#})/\sqrt{V_B{}^{\#} + 1}$ (as the permanent individual effect is absorbed into the latent residual to form the new unit variance). Rescaling conditional on average year conditions therefore gives $\eta = 0.175/\sqrt{0.105 + 1} = 0.166$ and $V_A = 0.122/(0.105 + 1) = 0.110$. Using these values in Equations (S3) and (S5) then allows calculating $\bar{\phi}$, $V_{\phi|\mathrm{A}}$, and $V_{\phi|\mathrm{A}}/\bar{\phi}$ and $V_{\phi|\mathrm{A}}/\bar{\phi}^2$ (see result above, and archived code: Acker et al., 2026).

*Example from the Soay sheep study of Stoffel et al. (2017)*

In Soay sheep, Stoffel et al. (2017) fitted a logit model that estimated an intercept ($\eta^{\#} = 0.53$), and effects of being a male ($\beta^{\#}_{male} = -0.56$), inbreeding, age class (with $\beta^{\#}_{midlife} = 3.61$), twin, and interaction between inbreeding and age class, plus breeding value ($a^{\#}$, with variance $V_A{}^{\#} = 0.3$), permanent individual ($b^{\#}$, with variance $V_B{}^{\#} = 0.3$), birth year, and capture year effects. Conditional on no inbreeding and no twin, in the midlife age class, in average birth year and capture year conditions, reformulating the original model into the baseline general model gives the following sex-specific expression: $\eta + a = (\eta^{\#} + \beta^{\#}_{sex} + \beta^{\#}_{midlife} + a^{\#})/\sqrt{V_B{}^{\#} + 1}$. Hence, we have $\eta = 0.53 + 3.61$ in females and $\eta = 0.53 + 3.61 - 0.56$ in males, and $V_A = 0.3/(\pi^2/3 + 0.3)$ (noting that $\pi^2/3$ is the latent residual variance implied by the logit model; see also Acker & Reid, 2025). To calculate $\bar{\phi}$, $V_{\phi|\mathrm{A}}$, and $V_{\phi|\mathrm{A}}/\bar{\phi}$, we then assume that the probit provides an adequate approximation for the logit rescaled to a unit residual variance and plug these values into Equations (S3) and (S5) (see results above, and archived code: Acker et al., 2026) or we could use existing exact solutions for the logit (available in Acker & Reid, 2025).

##### S1.4 Hypothetical relationships linking adaptive potential to environmental severity

To draw Figure 1 of the main text (see full details in archived code: Acker et al., 2026), we first envisaged hypothetical relationships between underlying $V_{\mathrm{A}}$ and $\bar{\phi}$. We drew these relationships using the baseline general model defined by Equation (S2). Indeed, any probit-scale animal model can always be reduced to Equation (S2), which implies a simple rescaling of the latent scale (and hence of $V_A$ and the underlying intercept $\eta$). Hence, the latent scale unit (which is the standard deviation of all the non-additive genetic effects under Equation S2, see above) is arbitrary and we omitted it on Figure 1. This has no consequence on the form of the relationship between $V_A$ and $\bar{\phi}$. Yet, we could then calculate $\eta$ as a function of $\bar{\phi}$ and $V_A$, and apply Equation (S5) to get $V_{\phi|A}$, and then calculate $V_{\phi|A}/\bar{\phi}$ and $V_{\phi|A}/\bar{\phi}^2$ (see details in archived code: Acker et al., 2026).

In Figure 1, we scaled all the relationships so that the minimum and maximum values of $V_A$ and $\bar{\phi}$ were identical in all panels. Specifically, we selected values of $\bar{\phi}$ that spanned 0.18–0.98 (and hence environmental severity, i.e. $1 - \bar{\phi}$, spanned 0.02–0.82), which corresponds to our estimated values in our focal shag population (see main text). For $V_A$, we chose values spanning 0.075–0.30, corresponding to an order of magnitude overlapping our empirical estimates in our defined good environment. Hence, these values are arbitrarily anchored into realistic ranges for the shag example. Figure 1 thus corresponds to potential relationships for a long-lived organism experiencing a wide range of environmental severities with plausible $V_A$ values. The qualitative transformation patterns (from $V_A$ to $V_{\phi|A}$, $V_{\phi|A}/\bar{\phi}$ and $V_{\phi|A}/\bar{\phi}^2$) corresponding to this example can become different with alternative parameter values. To illustrate this, Figure S2 shows outcomes given the same $\bar{\phi}$ as main Figure 1 with differing $V_A$, and Figure S3 illustrates outcomes given the same $V_A$ but differing $\bar{\phi}$.

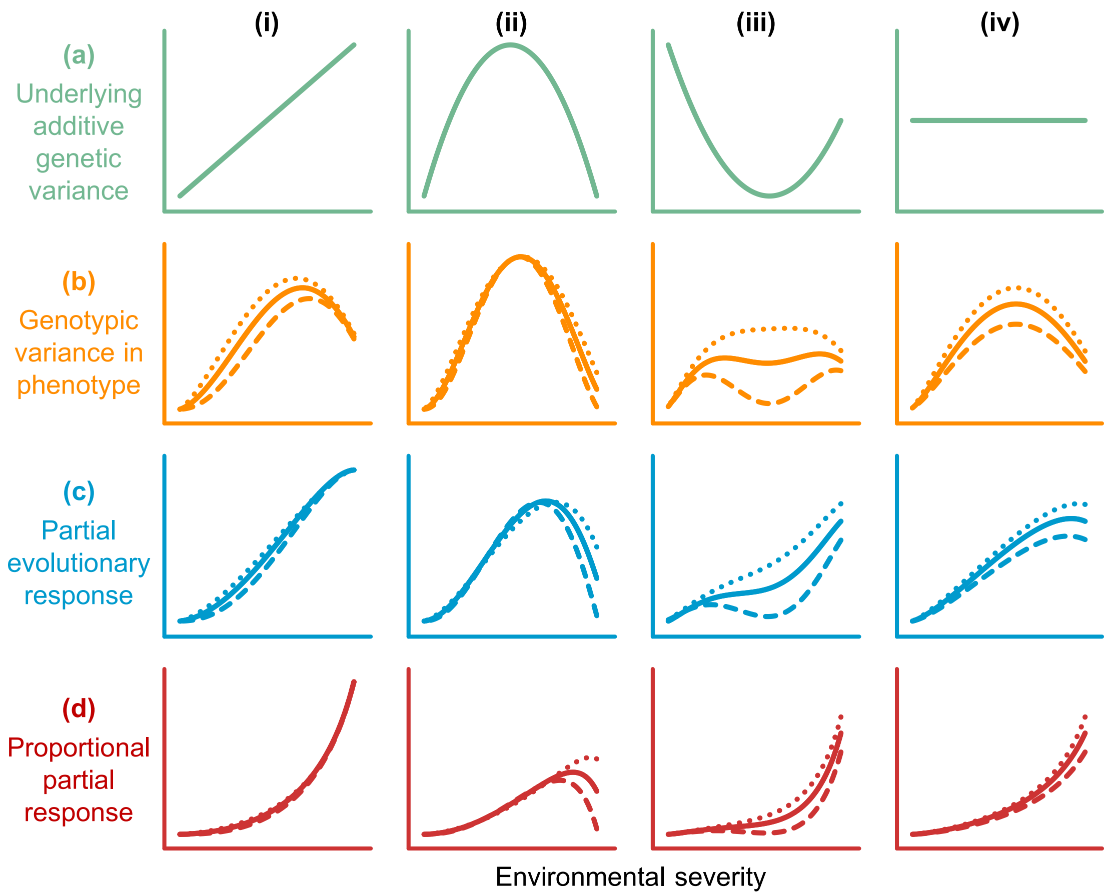


**Figure S2.** Hypothetical relationships between environmental severity and (a) latent-scale additive genetic variance ($V_{\mathrm{A}}$), (b) phenotypic-scale genotypic variance in survival ($V_{\phi|\mathrm{A}}$), and the partial evolutionary responses measured as the (c) absolute and (d) relative change in mean survival ($V_{\phi|\mathrm{A}}/\bar{\phi}$ and $V_{\phi|\mathrm{A}}/\bar{\phi}^2$, respectively). See Figure 1 of main text for further legend details. Here, each line type is a distinct set of parameter values, and the range of the x- and y-axis goes from the minimum to maximum value for the focal parameter set across (i) to (iv). All lines were drawn for $\bar{\phi}$ spanning 0.18–0.98. Solid lines correspond to Figure 1 (i.e. $V_{\mathrm{A}}$ spanning 0.075–0.30), and dotted and dashed lines correspond $V_{\mathrm{A}}$ spanning 0.005–0.15 and 0.30–0.75, respectively.

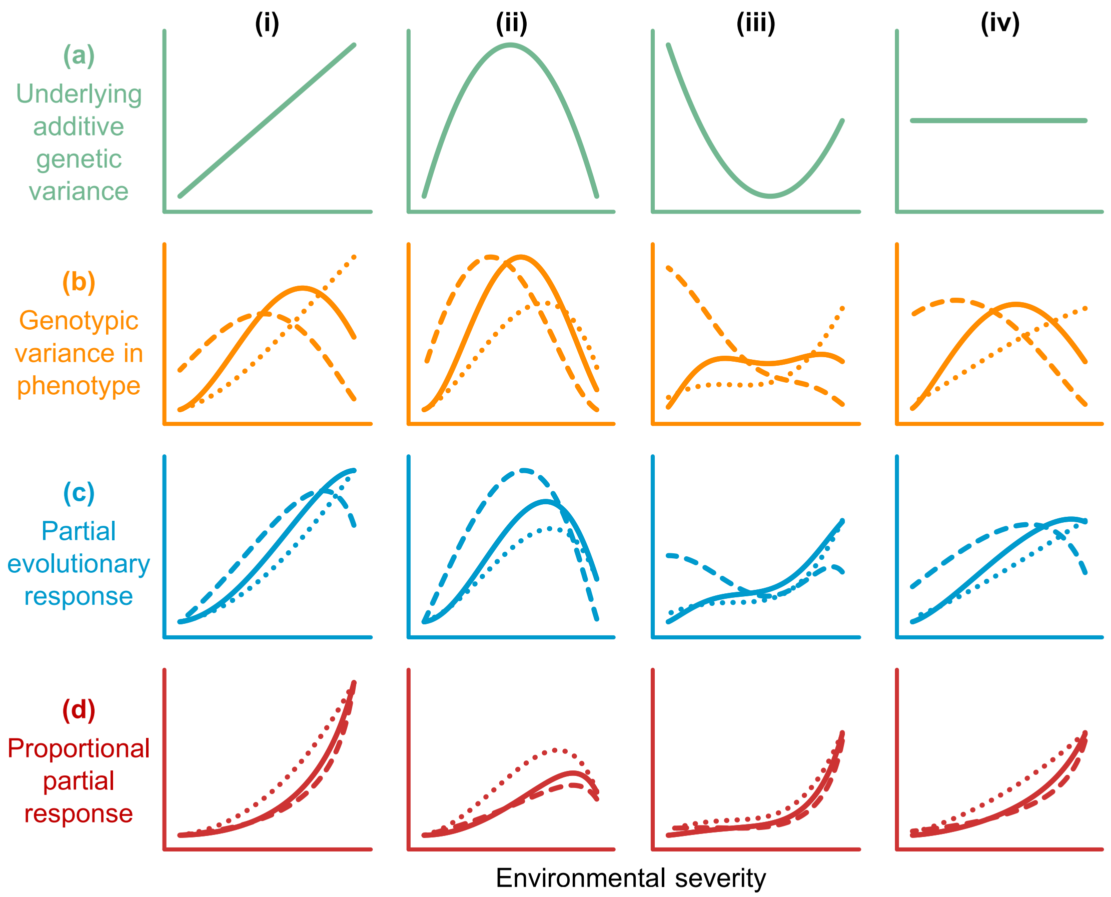


**Figure S3.** Hypothetical relationships between environmental severity and (a) latent-scale additive genetic variance ($V_{\mathrm{A}}$), (b) phenotypic-scale genotypic variance in survival ($V_{\phi|\mathrm{A}}$), and the partial evolutionary responses measured as the (c) absolute and (d) relative change in mean survival ($V_{\phi|\mathrm{A}}/\bar{\phi}$ and $V_{\phi|\mathrm{A}}/\bar{\phi}^2$, respectively). See Figure 1 of main text for further legend details. Here, each line type is a distinct set of parameter values, and the range of the x- and y-axis goes from the minimum to maximum value for the focal parameter set across (i) to (iv). All lines were drawn for $V_{\mathrm{A}}$ spanning 0.075–0.30. Solid lines correspond to Figure 1 of main text (i.e. $\bar{\phi}$ spanning 0.18–0.98), and dotted and dashed lines correspond to $\bar{\phi}$ spanning 0.05–0.65 and 0.65–0.95, respectively.

### Appendix S2

### Details of the datasets

#### S2.1 Study system and resighting data

*Susceptibility of Europeans shags to winter storms*

Previous analyses of resighting or recovery data from shags ringed on Isle of May ('IoM') showed that these birds experience occasional episodes of high mortality due to extreme climatic events, directly observed in their strongest forms through 'wrecks' of numerous dead individuals washed ashore during and immediately after spells of very stormy weather (Harris et al., 1998; Frederiksen et al., 2008; Acker et al., 2021). Shags are pursuit divers and have partially wettable feathers (considered an adaptation to underwater hunting) which they need to dry onshore (Grémillet et al., 1998; Grémillet et al., 2005; White et al., 2008). This appears to make them especially vulnerable to winter storms, as short daylength and low temperatures coinciding with strong winds, rough seas and/or high rainfall create lethal conditions whereby energetic costs of thermoregulation peak while access to fish is the lowest (Frederiksen et al., 2008). In years including varying occurrences, durations and intensities of winter storms, mean adult survival was previously shown to drop to 0.25–0.8 (Harris et al., 1998; Frederiksen et al., 2008; Acker et al., 2021). Detailed descriptions of multiple such events (including those captured by our present analyses) can be found in Harris et al. (1998), Frederiksen et al. (2008), and Acker et al. (2021).

*Details of the resighting data utilised in the present analyses*

Individual marking of IoM adult breeders and chicks using British Trust for Ornithology metal rings, and increasingly also three-letter engraved colour rings, was progressively introduced from 1981 (Harris et al., 1998; Frederiksen et al., 2008) and became systematic and almost comprehensive from 1997. Recapture efforts were then undertaken so that, as much as feasible, adults that were initially marked with only a metal ring have a field readable colour ring added. Accordingly, proportions of individuals still marked only with a metal ring decreased rapidly through the early 2000s. Yet, some individuals were never given a colour ring. While colour rings are readily readable from up to ~150m with a telescope, metal rings require catching individuals or longer sessions of close observations, which introduces heterogeneity in detection (that our capture-recapture model parameterisation handles through resighting probabilities dependent on ring type; Appendix S3). Accordingly, hereafter Tables S1-S2 report on sample sizes available in the resighting data by summarising the absolute frequencies of sighted individuals in breeding seasons 2000–2024 dependent on ring type in each sex and year, in the full dataset with all individuals and the dataset including only known age individuals. Sample sizes are also reported for the additional autumn 2024 occasion which serves to ensure independent estimation of survival probability from breeding season 2023 to 2024 and detection probability in breeding season 2024 (see main text).

From Tables S1-S2, we note that the amount of data clearly differed among the three categories of environmental conditions, as would be expected given the respective temporal distribution of good, bad, and very bad years. Nevertheless, large numbers of individuals were known to be alive at the start of each biological year that then experienced each environment: 2972 in good years, 1464 in bad years, and 1988 in very bad years. Multiple individuals were known to be alive in more than one year of each category. Specifically, the mean number of years in which an individual was known to be alive was 7.36 under good conditions (SD: 1.37, range 1–9), 1.88 under bad conditions (SD 0.97, range 1–6), and 1.07 under very bad conditions (SD: 0.25, range 1–9). These numbers highlight how data availability on individual variation in survival becomes increasingly restricted with environmental severity.

**Table S1.** Numbers of individuals observed for the first time (i.e. when they entered the dataset, at their first recorded breeding attempt during the study period), or subsequently observed (i.e. after they entered the dataset) while wearing only a metal ring or wearing a colour ring, in each sighting occasion in the full dataset (fitted to our initial capture-recapture model to estimate sex-by-year specific survival probabilities).

| Sighting occasion | Females | | | | Males | | | |
|---|---|---|---|---|---|---|---|---|
| | First obs. | Subsequent obs. | | Total | First obs. | Subsequent obs. | | Total |
| | | *Metal* | *Colour* | | | *Metal* | *Colour* | |
| 2000 | 80 | – | – | 80 | 97 | – | – | 97 |
| 2001 | 162 | 8 | 59 | 229 | 169 | 7 | 73 | 249 |
| 2002 | 105 | 34 | 160 | 299 | 100 | 32 | 194 | 326 |
| 2003 | 98 | 21 | 185 | 304 | 104 | 36 | 215 | 355 |
| 2004 ⦿ | 18 | 3 | 209 | 230 | 19 | 15 | 243 | 277 |
| 2005 ⊗ | 51 | 2 | 58 | 111 | 58 | 3 | 152 | 213 |
| 2006 ⊗ | 134 | 3 | 95 | 232 | 126 | 17 | 199 | 342 |
| 2007 ⊗ | 112 | 4 | 128 | 244 | 102 | 10 | 209 | 321 |
| 2008 | 72 | 12 | 199 | 283 | 60 | 6 | 282 | 348 |
| 2009 | 77 | 11 | 264 | 352 | 57 | 7 | 335 | 399 |
| 2010 | 70 | 9 | 321 | 400 | 64 | 7 | 365 | 436 |
| 2011 | 140 | 3 | 365 | 508 | 116 | 7 | 400 | 523 |
| 2012 ⦿ | 157 | 5 | 437 | 599 | 140 | 2 | 473 | 615 |
| 2013 ⊗ | 90 | 2 | 200 | 292 | 54 | 2 | 315 | 371 |
| 2014 | 93 | 3 | 210 | 306 | 73 | 2 | 301 | 376 |
| 2015 | 58 | 3 | 284 | 345 | 55 | 2 | 342 | 399 |
| 2016 | 51 | 2 | 312 | 365 | 59 | 2 | 340 | 401 |
| 2017 ⊗ | 100 | 0 | 325 | 425 | 91 | 0 | 350 | 441 |
| 2018 ⊗ | 81 | 0 | 296 | 377 | 78 | 0 | 333 | 411 |
| 2019 | 92 | 0 | 306 | 398 | 88 | 0 | 356 | 444 |
| 2020 | 45 | 1 | 284 | 330 | 38 | 0 | 349 | 387 |
| 2021 | 57 | 1 | 361 | 419 | 72 | 0 | 375 | 447 |
| 2022 | 65 | 1 | 376 | 442 | 66 | 1 | 382 | 449 |
| 2023 ⦿ | 88 | 2 | 419 | 509 | 130 | 1 | 428 | 559 |
| 2024 | 0 | 0 | 79 | 79 | 0 | 0 | 97 | 97 |
| Autumn 2024 | 0 | 0 | 65 | 65 | 0 | 0 | 81 | 81 |

Note: Symbols '⊗' and '⦿' indicate bad and very bad subsequent year intervals, respectively.

**Table S2.** Numbers of individuals observed for the first time (i.e. when they entered the dataset, at their first recorded breeding attempt during the study period), or subsequently observed (i.e. after they entered the dataset) while wearing only a metal ring or wearing a colour ring, in each sighting occasion in the dataset including only known age individuals (fitted to our joint capture-recapture animal models to conduct quantitative genetic decomposition of variance in survival).

| Sighting occasion | Females | | | | Males | | | |
|---|---|---|---|---|---|---|---|---|
| | First obs. | Subsequent obs. | | Total | First obs. | Subsequent obs. | | Total |
| | | *Metal* | *Colour* | | | *Metal* | *Colour* | |
| 2000 | 52 | – | – | 52 | 71 | – | – | 71 |
| 2001 | 96 | 4 | 39 | 139 | 116 | 5 | 54 | 175 |
| 2002 | 74 | 19 | 106 | 199 | 62 | 23 | 142 | 227 |
| 2003 | 65 | 11 | 126 | 202 | 71 | 20 | 165 | 256 |
| 2004 ⦿ | 15 | 1 | 156 | 172 | 15 | 4 | 193 | 212 |
| 2005 ⊗ | 28 | 1 | 44 | 73 | 31 | 3 | 130 | 164 |
| 2006 ⊗ | 78 | 3 | 59 | 140 | 86 | 12 | 151 | 249 |
| 2007 ⊗ | 50 | 3 | 84 | 137 | 50 | 6 | 157 | 213 |
| 2008 | 42 | 7 | 119 | 168 | 31 | 1 | 192 | 224 |
| 2009 | 41 | 6 | 156 | 203 | 40 | 2 | 214 | 256 |
| 2010 | 48 | 6 | 184 | 238 | 47 | 3 | 235 | 285 |
| 2011 | 105 | 1 | 214 | 320 | 93 | 4 | 261 | 358 |
| 2012 ⦿ | 128 | 3 | 273 | 404 | 121 | 2 | 321 | 444 |
| 2013 ⊗ | 79 | 1 | 139 | 219 | 43 | 2 | 234 | 279 |
| 2014 | 69 | 3 | 159 | 231 | 55 | 2 | 226 | 283 |
| 2015 | 23 | 2 | 212 | 237 | 33 | 2 | 256 | 291 |
| 2016 | 29 | 1 | 213 | 243 | 47 | 2 | 252 | 301 |
| 2017 ⊗ | 90 | 0 | 212 | 302 | 84 | 0 | 264 | 348 |
| 2018 ⊗ | 72 | 0 | 216 | 288 | 71 | 0 | 269 | 340 |
| 2019 | 72 | 0 | 234 | 306 | 68 | 0 | 298 | 366 |
| 2020 | 43 | 1 | 219 | 263 | 36 | 0 | 282 | 318 |
| 2021 | 49 | 1 | 286 | 336 | 63 | 0 | 313 | 376 |
| 2022 | 56 | 1 | 305 | 362 | 56 | 0 | 321 | 377 |
| 2023 ⦿ | 71 | 2 | 339 | 412 | 108 | 0 | 367 | 475 |
| 2024 | 0 | 0 | 67 | 67 | 0 | 0 | 88 | 88 |
| Autumn 2024 | 0 | 0 | 55 | 55 | 0 | 0 | 76 | 76 |

Note: Symbols '⊗' and '⦿' indicate bad and very bad subsequent year intervals, respectively.

The annual age distributions of resighted adult individuals are shown in Figure S4 hereafter. Evidently, the bulk of adult individuals is typically around age 3-9 years, with older individuals around age 10-15 years being present in notable frequencies across all years, and typically only few individuals of age >15 years (Figure S4).

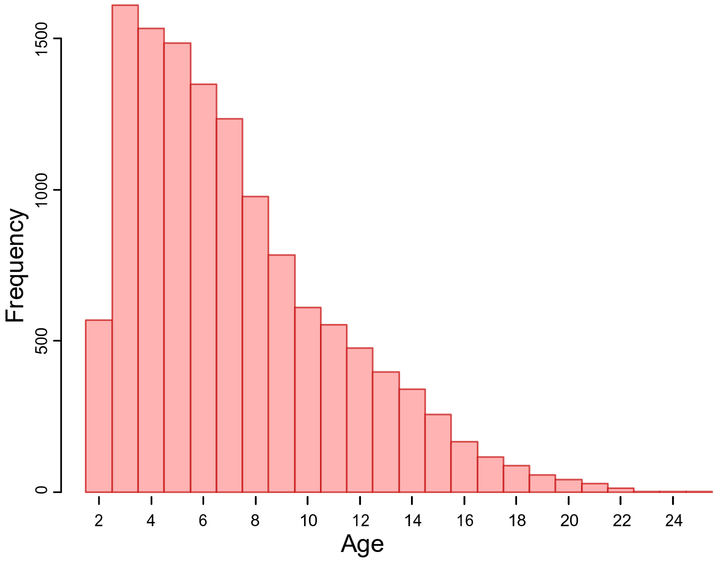


**Figure S4.** Age distribution of resighted individuals across all years. Note that every individual enters the dataset at its first recorded breeding attempt during the study period, and the youngest age at first breeding is 2 years old. The mean age across all annual sightings was 7.246 years old. The oldest individual of known age that was ever resighted was 25 years old.

#### S2.2 Structure of the pedigree data

*Details of the relatedness data*

To conduct our quantitative genetic analyses of the 3455 individuals of known age included in our resighting dataset (hereafter 'focal individuals'), we exploited the relatedness data available given known parental identities linking focal individuals to each other, forming a 'utilised pedigree' that comprised 482 additional ancestors and spanned up to 7 generations. Among those additional ancestors, 158 corresponded to 'dummy' parent identities assigned to link known full siblings of the same brood from an unringed breeding pair. In the utilised pedigree, 891 individuals were founders (i.e. had offspring in the pedigree but no known parents), 208 were half-founders (i.e. had offspring and one unknown parent), 460 were non-founder parents (i.e. had offspring and two known parents), and 1353 were strict offspring (i.e. had at least one known parent but no offspring). Finally, 543 individuals were not related to any other focal individual.

More precisely, the utilised pedigree contained 913 unique pairs with both parents identified, 310 broods with more than one offspring, 388 full sibships (i.e. cases of >1 offspring of the same pair), and 564 half sibships (i.e. cases of >1 offspring of the same parent belonging to >1 pair).

Further, there were 762 identified mothers in total, and 492 maternal sibships (mothers with >1 offspring), including 249 maternal half-sibships (mothers with >1 offspring from >1 fathers). There were 798 identified fathers in total, and 528 paternal sibships, including 315 paternal half-sibships. In total, 900 individuals had at least one full sibling, 652 had at least one same-brood full sibling (556 had two and 96 had three), and 2006 had at least one half-sibling.

As expected, given the above elements of pedigree structure, most of the 18,665 pairwise links between focal individuals corresponded to close kinship (22% ≥0.25, 50% ≥0.125, and 73% ≥0.0625, representing first, second and third-order relatives, respectively; see details in Table S3), and many focal individuals were linked to multiple other focal individuals (Figure S5). These links provided dense data to estimate $V_A$ in survival. The diversity of these links further provided us with relevant information to check whether shared developmental micro-environments associated with the brood, mother, or father identity also caused resemblance among close kin that could have inflated our estimates of additive genetic variance (see also Appendix S3).

**Table S3**. Distribution of kinship (co-ancestry) coefficient values across all non-zero pairwise links between phenotype-informative individuals in the utilized pedigree.

| Coefficient of kinship | Frequency of link | Proportion of all links |
|---|---|---|
| 0.000977 | 10 | 0.05 % |
| 0.001953 | 72 | 0.39 % |
| 0.003906 | 252 | 1.35 % |
| 0.007813 | 609 | 3.26 % |
| 0.015625 | 1417 | 7.59 % |
| 0.03125 | 2676 | 14.34 % |
| 0.039063 | 1 | 0.01 % |
| 0.046875 | 6 | 0.03 % |
| 0.0625 | 4285 | 22.96 % |
| 0.09375 | 15 | 0.08 % |
| 0.125 | 5172 | 27.71 % |
| 0.140625 | 1 | 0.01 % |
| 0.15625 | 2 | 0.01 % |
| 0.1875 | 13 | 0.07 % |
| 0.25 | 4122 | 22.08 % |
| 0.28125 | 5 | 0.03 % |
| 0.3125 | 5 | 0.03 % |
| 0.375 | 2 | 0.01 % |

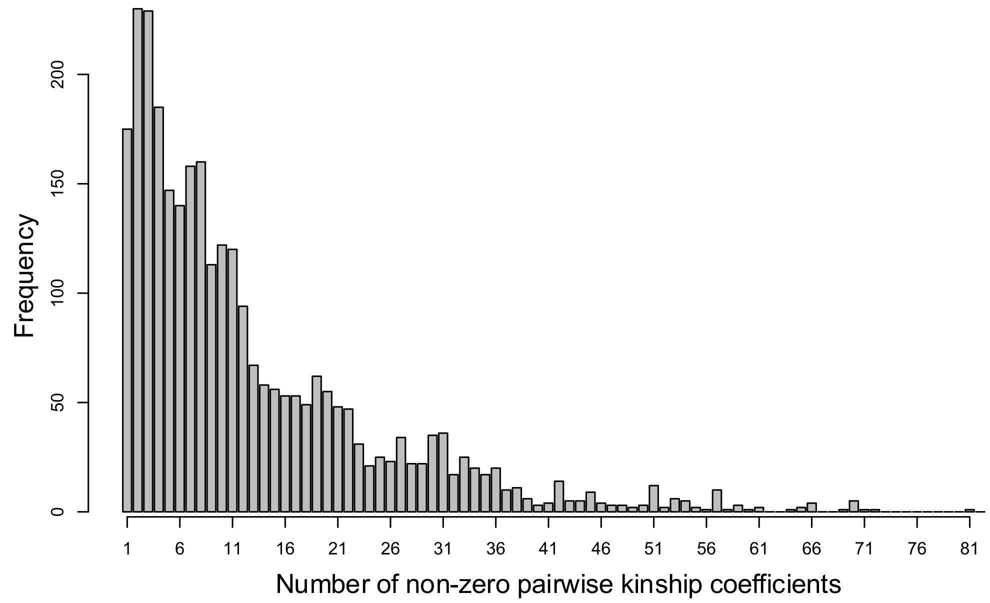


**Figure S5**. Distribution of the per-individual number of non-zero pairwise links between phenotyped individuals.

*Pedigree accuracy*

We used social parentage data, as commonly done in monogamous birds (e.g. Sheldon et al., 2003; Teplitsky et al., 2009; Charmantier et al., 2011; Husby et al., 2011), yet this might induce pedigree errors, due to non-exhaustiveness in individual identification and extra-pair paternity. In general, such errors are expected to lead to slightly downwardly biased estimates of additive genetic variance (since existing resemblance due to inherited variation is not assigned to any relatedness through the observed pedigree, and hence part of the genetic variance is missed). However, we expected few pedigree gaps given the high proportion of known parents (see main text). Further, an early study conducted over 4 years and ~25 pairs followed annually on IoM found an extra-pair paternity rate of only 9.3% (out of 161 chicks from 87 pairs; Graves et al., 1993). These values match the 0–10% extra-pair paternity rates found in other Phalacrocoracidae species (Imperial shag, *Leucocarbo atriceps*: Calderon et al., 2012; great cormorants, *Phalacrocorax carbo*: Minias et al., 2015). This implies that ≥95% of all assigned parental links will be correct (assuming that all mothers are correctly assigned from field observations). Previous simulations have indicated that such low extra-pair paternity rates are likely to cause minimal underestimation of additive genetic variance (Charmantier & Réale, 2005; Firth et al., 2015).

### Appendix S3

### Details of the statistical modelling

#### S3.1 Implementation of the capture-recapture model

To implement our capture-recapture model, we formulated the likelihood of the capture-resighting history $H_i$ of each individual $i$ (i.e. the sequence of 0/1 observations, for 'not seen' vs 'seen alive' on each annual breeding season $t$ until the last one $T$) following well-established principles of Cormack-Jolly-Seber model analyses (Lebreton et al., 1992). More specifically, every history can be divided in two independent parts. First, for each year $t$ after the year individual $i$ entered the dataset, $\zeta_i$, and until the last year $i$ was seen, $\lambda_i$, we know for certainty that $i$ was alive. Therefore, the likelihood for the observation event in year $t$ is given by the probability that individual $i$ survived from the previous breeding season to the focal breeding season times the probability of the focal observation event. That is, $\phi_{s(i),t-1}p_{s(i),t,r(i,t)}$ or $\phi_{s(i),t-1}\left(1-p_{s(i),t,r(i,t)}\right)$, respectively, if the individual was seen or not seen, where $s(i)$ is the sex of $i$ and and $r(i,t)$ is the ring type of $i$ at $t$ (see main text). The second part of $H_i$ corresponds to all years after $\lambda_i$. Here, we do not know whether the individual was still alive or not, which makes the likelihood of each subsequent observation within $H_i$ conditional on the previous one. Yet, we can formulate the likelihood for this full second part of $H_i$ as the probability $\chi_{i,\lambda_i}$ that $i$ was never seen again after $\lambda_i$. This can is achieved through the following backward recursion for calculating $\chi_{i,t}$, defined from $t=T-1$ and until $t=\lambda_i$, noting that $\chi_{i,T}$=1 (since by definition the probability that an individual is never seen after $T$ is 1):

$$\chi_{i,t}=\left(1-\phi_{s(i),t}\right)+\phi_{s(i),t}\left(1-p_{s(i),t,r(i,t)}\right)\chi_{i,t+1}. \qquad \text{(S39)}$$

The likelihood for $H_i$ is then given by the product of the likelihoods for each observation in the first part and the likelihood for the second part. And the likelihood for all the $H_i$ is the product of their likelihoods.

Moreover, to calculate the population mean survival probability in each year, $\bar{\phi}_t$, across both sexes (as a function of the parameters $\phi_{s,t}$), we need to average over all alive individuals. This requires calculating a weighted mean of $\phi_{\text{♀},t}$ and $\phi_{\text{♂},t}$ where the weights are frequencies of males and females representative of their relative proportions at $t$. Yet, because survival and detection are potentially different between both sexes, observed frequencies may be biased. Appropriate frequencies are therefore calculated as expected frequencies of individuals that were alive at $t$ in the dataset, given the observations and the model. These can be calculated as the sum of the probability that $i$ was alive at time $t$ conditional on $H_i$, $\pi_{i,t} = P(z_{i,t} = 1|H_i)$, across all females and all males respectively. Here, we know that $i$ was alive in the adult population from $\zeta_i$ to $\lambda_i$ and, given Bayes' theorem, after $\lambda_i$, $\pi_{i,t}$ is just the probability that $i$ survived from $\lambda_i$ to $t$ but was not resighted then and was never seen again after $t$ divided by $\chi_{i,\lambda_i}$. Accordingly, we have:

$$\begin{cases} \pi_{i,t} = 1 & \text{for } \zeta_i < t \leq \lambda_i \\ \pi_{i,t} = \dfrac{\left(\prod_{u=\lambda_i}^{t-1} \phi_u\right)\left(\prod_{v=\lambda_i+1}^{t} (1 - p_v)\right)\chi_{i,t}}{\chi_{i,\lambda_i}} & \text{for } \lambda_i < t \leq T \end{cases}, \tag{S40}$$

and $\bar{\phi}_t$ is thus given by the following weighted mean:

$$\bar{\phi}_t = \frac{\sum_{i\in\{\text{♀}\}} \pi_{i,t}\phi_{\text{♀},t} + \sum_{i\in\{\text{♂}\}} \pi_{i,t}\phi_{\text{♀},t}}{\sum_i \pi_{i,t}}, \tag{S41}$$

where {♀} is the set of females and {♂} is the set of males.

**S3.2 Implementation and key assumptions of the animal model**

To implement our animal model (integrated within the capture-recapture model), we formulated the structure of additive genetic covariance through an algorithm defining a directed acyclic graph that explicitly represents the process of inheritance (Steinsland & Jensen, 2009; Waldmann, 2009). This method is computationally faster and more flexible than using the full $A$ matrix in lieu of a correlation matrix to express the multivariate normal distribution of breeding values (as would be standard given the general formulation of GLMMs; Kruuk, 2004).

As for any animal model, our analyses assume that each individual $i$ descends from a phantom base population of unrelated individuals under panmixia, including any missing parents in the pedigree. For each individual $k$ in the base population, the breeding value is distributed such that $a_k \sim \mathcal{N}(0, V_\mathrm{A})$. The assumption of panmixia is then relaxed downstream across the pedigree. As inheritance proceeds through sexual reproduction, $a_i$ is composed of the midparent breeding value plus the Mendelian sampling deviation due to recombination and segregation:

$$a_i = \left(a_{p(i)} + a_{q(i)}\right)/2 + m_i, \tag{S42}$$

where $p$ and $q$ denote the two parents. The sampling deviation is distributed such that $m_i \sim \mathcal{N}\left(0,\ V_\mathrm{A}/2\left(1 - \left(F_{p(i)} + F_{q(i)}\right)/2\right)\right)$, where $F_{p(i)}$ and $F_{q(i)}$ are the inbreeding coefficients of the parents of individual *j*. Note that, for any individual *j* (which could $p(i)$ or $q(i)$, or *i* itself) $F_j$ is simply the coefficient of coancestry (or kinship), $\theta$, between its parents: $F_j = \theta_{p(j),q(j)}$. Further, we assume that all individuals $k$ in the base population are non-inbred, i.e. have $F_k = 0$. Full implementation details are available in the Stan code available via the data archive for our study (Acker et al., 2026).

Hence, importantly, $V_\mathrm{A}$ is twice the expected within-family variance in breeding values (i.e. effectively the Mendelian sampling variance) for unrelated outbred individuals under random mating. Because this mating pattern represents a reasonable assumption for IoM shags, $V_\mathrm{A}$ corresponds to approximately twice the within-family variance evident in the population.

##### S3.3 Prior specifications

For the initial capture-recapture model estimating sex-by-year specific survival probabilities, as detailed in the main text, all parameters ($\phi_{s,t}$ and $p_{s,t,r}$) were single probabilities of binary outcomes (survival vs death, and resighting vs detection failure). Here, we used continuous uniform prior distributions over [0,1], $\mathcal{U}(0,1)$, for all parameters, thereby remaining objective regarding our general preconceptions on the values that the parameters could take. The results (main Figure 3) clearly show that the information contained in the data overrode the prior.

For the joint capture-recapture animal models, resighting remained parameterised by single probabilities ($p_{s,t,r}$), and so we retained the $\mathcal{U}(0,1)$ prior. Meanwhile, survival was parameterised through ‘fixed’ effects ($\mu$, $\beta_1$ and $\beta_2$) and ‘random’ effect variances ($V_{\gamma,s(i)}$ and $V_\mathrm{C}$ or $V_B$ and $V_\mathrm{A}$ [or $V_\mathrm{A}^k$ for environment-specific models]) of a probit regression (main Equations 1–3). Here, we chose weakly informative priors for all parameters following typical recommendations for robust (binary) GLMMs (Gelman, 20006; Gelman et al., 2013; Gelman et al., 2017; Ghosh et al., 2018). Specifically, for the fixed effects we used the location-scale Student’s $t$ distribution, with 4 degrees of freedom, location 0, and scale 1, $\ell st(4,0,1)$. For the variance parameters, we used location-scale half-Student’s $t$ distributions (i.e. $\ell st$ truncated to only have nonzero probability density for positive values), $H\text{-}\ell st(4,0,1)$ for the standard deviation of the year effects, $\sqrt{V_{\gamma,s(i)}}$, and $H\text{-}\ell st(4,0,0.5)$ for the standard deviations of the other random effects. Such distributions imply priors on the variance with a prominent mode on zero, and a heavy tail still allowing larger values if the data demands it. Note that the scale of 1 represents a useful starting default in standard probit regressions, because it is the scale of the implicitly-defined residuals (see explanations following Equation S2 above). Yet, the value of 0.5 was chosen to better match the actual scale of $\sqrt{V_\mathrm{C}}$, $\sqrt{V_B}$ and $\sqrt{V_\mathrm{A}}$ (and thereby improve posterior sampling) after preliminary analyses using the $H\text{-}\ell st(4,0,1)$ prior revealed that these parameters were about an order of magnitude smaller than the unit (and MCMC diagnostics indicated sampling difficulties).

Moreover, in models including cross-environment or cross-genetic genetic correlations, we used the Lewandowski-Kurowicka-Joe prior with shape 1, $\mathcal{LKJ}(1)$, on the correlation matrix. This prior implies a uniform density over the space of possible correlation matrices, and it is typically recommended in lieu of the Wishart because it is less informative as it lets the estimation of correlation be independent of the specification of scale (Gelman et al., 2013; McElreath, 2020).

**S3.4 Additional model versions and checks**

*Models with brood, maternal, or paternal effects*

To check whether common early-life environmental effects among close relatives could have inflated our estimates of $V_\mathrm{A}$ and $V_\mathrm{A}^k$, we ran three additional models for both the overall animal model (main Equation 2) and the environment-specific animal model (main Equation 3). In both cases, the three models comprised one with brood effects, one with maternal effects, and one with paternal effects. Here, the animal models were formulated, respectively as:

$$\mathrm{probit}\big(\phi_{i,t}\big) = \mu + \gamma_{s(i),t} + \beta_1 x_{i,t} + \beta_2 {x_{i,t}}^2 + \kappa_{idx(i)} + b_i + a_i, \quad \text{(S43)}$$

$$\mathrm{probit}\big(\phi_{i,t}\big) = \mu + \beta_1 x_{i,t} + \beta_2 {x_{i,t}}^2 + \gamma_{s(i),t} + \kappa_{idx(i)} + b_i + a_{k(t),i}. \quad \text{(S44)}$$

where parameters are as in Equations (2-3) of the main text, $\kappa$ is the focal kin effect (i.e. either brood, maternal, or paternal effect) and $idx(i)$ is the focal kin identity of individual $i$ (i.e. brood, mother, or father identity), and $d_{idx(i)} \sim \mathcal{N}(0, V_\kappa)$.

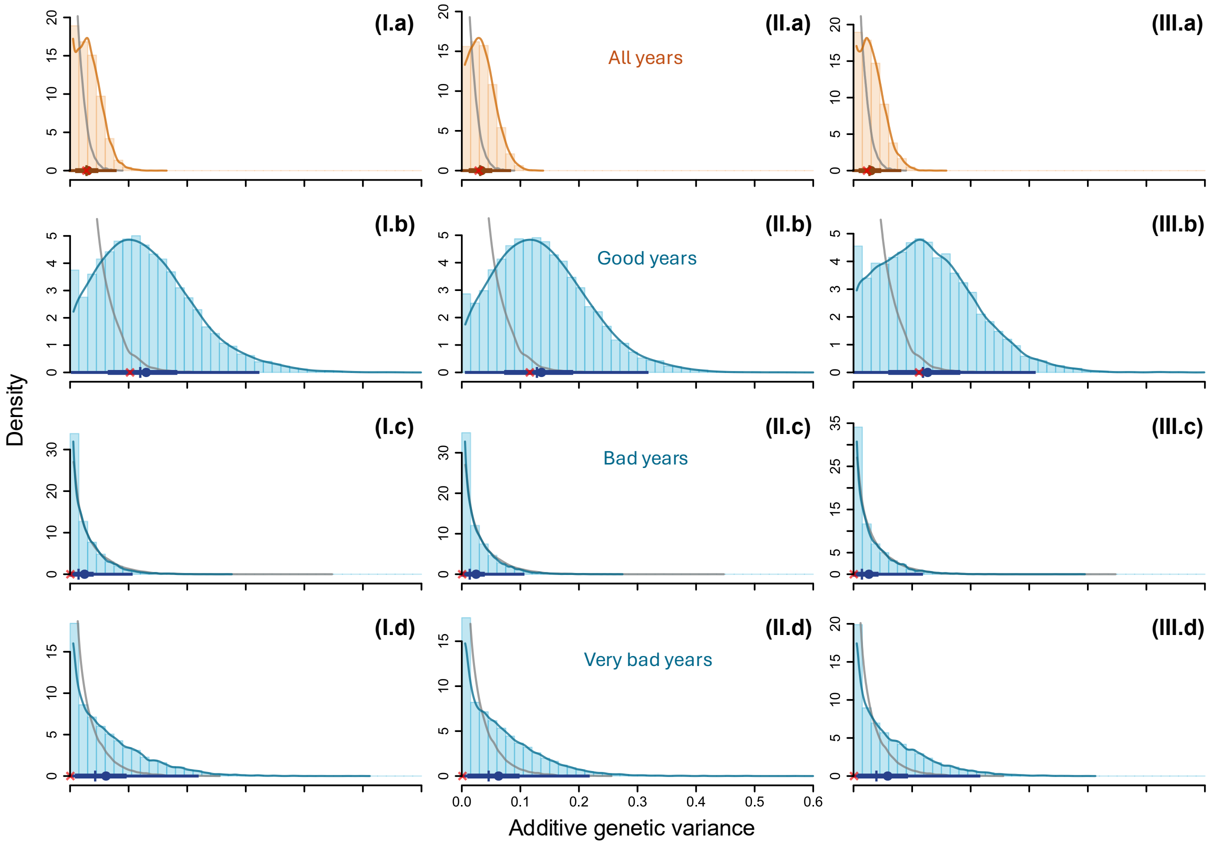


**Figure S6**. Posterior distributions of additive genetic variance underlying annual adult survival estimated in models that included additional (I) brood, (II) maternal, or (III) paternal effects; (I-III.a) across all years ($V_{\mathrm{A}}$), and separately in years with (I-III.b) good, (I-III.c) bad, and (I-III.c) very bad environmental conditions ($V_{\mathrm{A}}^{k}$). Histogram bars show the distribution of the posterior samples, with corresponding kernel density estimation in solid blue line. The superimposed thin and thick horizontal dark-blue lines show the 95% and 50% credible intervals, respectively. Along these horizontal lines, the plain circle, vertical tick and red cross show the mean, median, and mode, respectively. The solid grey curve shows the density of the posterior distribution for the corresponding 'null expectation' (see main text and below).

The posterior distributions of $V_{\mathrm{A}}$ and $V_{\mathrm{A}}^{k}$ in these models were systematically similar to the posterior distributions obtained in the models presented in main text (which did not include any kin effect; Figure S6, vs Figure 4 of the main text; Table S4), and there was no or weak evidence of kin effects (Figure S7, Table S4). Specifically, in all these models the additive genetic variance still clearly peaked away from zero, while the mode of the focal kin variance was always on zero, indicating that kin variance was small or negligible.

**Table S4**. Estimates of the additive genetic variance in survival and kin effect variance from the additional models including either brood, maternal, or paternal effects. In each case, estimates were obtained from a model with one $V_{\mathrm{A}}$ parameter defined across all environments, or three $V_{\mathrm{A}}^{k}$ parameter defined for good, bad, and very bad years.

| Model | | Parameter | Posterior summary | | |
|---|---|---|---|---|---|
| Kin effect | Environments | | Mean | Median | 95% CRI |
| Brood | Together | $V_{\mathrm{A}}$ | 0.030 | 0.028 | [0.000, 0.077] |
| | | $V_{\kappa}$ | 0.028 | 0.021 | [0.000, 0.088] |
| | Separate | $V_{\mathrm{A}}^{good}$ | 0.130 | 0.12 | [0.004, 0.321] |
| | | $V_{\mathrm{A}}^{bad}$ | 0.025 | 0.014 | [0.000, 0.104] |
| | | $V_{\mathrm{A}}^{vbad}$ | 0.061 | 0.043 | [0.000, 0.217] |
| | | $V_{\kappa}$ | 0.028 | 0.021 | [0.000, 0.088] |
| Mother | Together | $V_{\mathrm{A}}$ | 0.033 | 0.031 | [0.001, 0.082] |
| | | $V_{\kappa}$ | 0.011 | 0.006 | [0.000, 0.044] |
| | Separate | $V_{\mathrm{A}}^{good}$ | 0.136 | 0.128 | [0.008, 0.316] |
| | | $V_{\mathrm{A}}^{bad}$ | 0.024 | 0.014 | [0.000, 0.104] |
| | | $V_{\mathrm{A}}^{vbad}$ | 0.063 | 0.046 | [0.000, 0.216] |
| | | $V_{\kappa}$ | 0.011 | 0.006 | [0.000, 0.044] |
| Father | Together | $V_{\mathrm{A}}$ | 0.030 | 0.027 | [0.000, 0.079] |
| | | $V_{\kappa}$ | 0.012 | 0.008 | [0.000, 0.044] |
| | Separate | $V_{\mathrm{A}}^{good}$ | 0.127 | 0.119 | [0.002, 0.309] |
| | | $V_{\mathrm{A}}^{bad}$ | 0.026 | 0.014 | [0.000, 0.116] |
| | | $V_{\mathrm{A}}^{vbad}$ | 0.058 | 0.039 | [0.000, 0.214] |
| | | $V_{\kappa}$ | 0.012 | 0.008 | [0.000, 0.044] |

Note: ‘95% CRI’ is the 95% credible interval.

Note that here, MCMC sampling was more challenging than with the main models, and there were few occasional divergent iterations. This is likely because the inference on liability-scale variances of both additive genetic and kin effects is highly data-demanding, and our dataset probably lies at the lower margin of what is necessary to estimate the additional effects of interest.

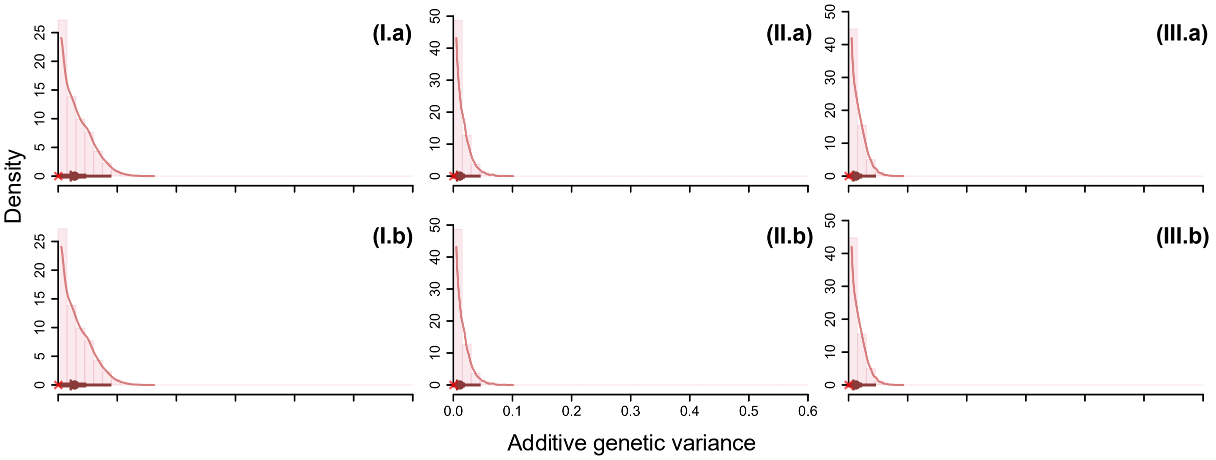


**Figure S7**. Posterior distributions of the variance in the kin effect ($V_{\mathrm{k}}$) in models that included additional (I) brood, (II) maternal, or (III) paternal effects; and additive genetic variance was formulated (I-III.a) across all years ($V_{\mathrm{A}}$; Equation S43), and (I-III.b) as environment-specific ($V_{\mathrm{A}}^{k}$; Equation S44). Histogram bars show the distribution of the posterior samples, with corresponding kernel density estimation in solid line. The superimposed thin and thick horizontal dark lines at the bottom show the 95% and 50% credible intervals, respectively. Along these horizontal lines, the plain circle, vertical tick and red cross show the mean, median, and mode, respectively.

*Models with environment-specific permanent individual effects*

In search of an appropriate specification of individual heterogeneity in our main models (Equations 2-3 of main text), we initially tested an alternative parameterisation where the permanent individual effects were environment-specific, such that:

$$\mathrm{probit}(\phi_{i,t}) = \mu + \beta_1 x_{i,t} + \beta_2 {x_{i,t}}^2 + \gamma_{s(i),t} + \chi_{idx(i)} + b_{k(t),i} + a_{k(t),i}, \qquad \text{(S45)}$$

where $b_{k(t),i} \sim \mathcal{N}(0, V_{\mathrm{B}}^{k})$, and all other parameters are as in Equation (3) of the main text.

As expected, given the limited number of repeated within-individual exposure to bad and very bad environmental conditions (see Appendix S2), we found negligible $V_{\mathrm{B}}^{bad}$ and $V_{\mathrm{B}}^{vbad}$ (Figure S8). We also found negligible $V_{\mathrm{B}}^{good}$ (Figure S8). Accordingly, and since $V_{\mathrm{B}}$ was also negligible in our original main model (see main text and Appendix S4), there was no perceptible difference in $V_{\mathrm{A}}^{k}$ estimates between the two model versions (Figure S8 vs Figure 4 of main text). Specifically, the posterior mean [95% credible interval (or 'CRI')] was $0.084\,[0.000, 0.311]$ for $V_{\mathrm{B}}^{good}$,

0.031 [0.000, 0.258] for $V_B^{bad}$, and 0.059 [0.000, 0.404] for $V_B^{vbad}$ and 0.156 [0.008, 0.364] for $V_A^{good}$, 0.031 [0.000, 0.129] for $V_A^{bad}$, and 0.059 [0.014, 0.211] for $V_A^{vbad}$.

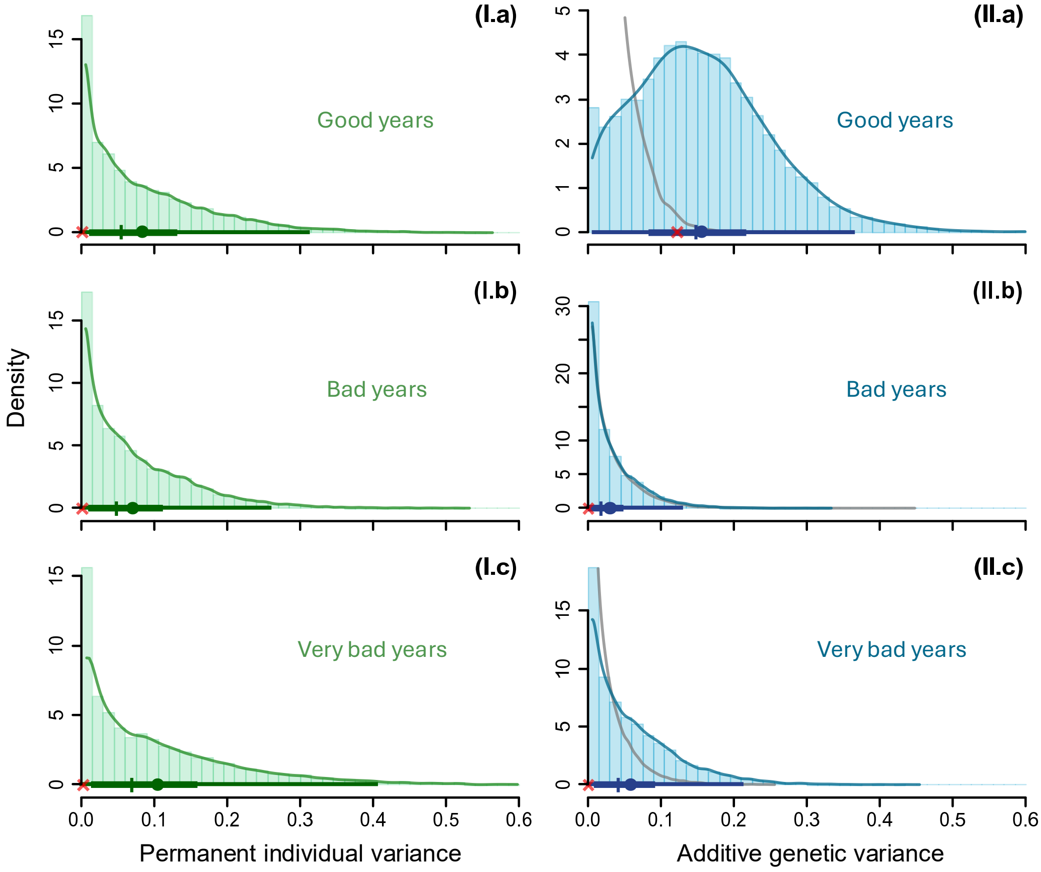


**Figure S8**. Posterior distributions of the environment-specific (I) permanent individual and (II) additive genetic variances ($V_{B,k}$ and $V_{A,k}$) underlying annual adult survival under (a) good, (b) bad, and (c) very bad environmental conditions. Estimates are from the alternative model defined in Equation (S45). Histogram bars show the distribution of the posterior samples, with corresponding kernel density estimation in solid line. The superimposed thin and thick horizontal lines show the 95% and 50% credible intervals, respectively. Along these horizontal lines, the plain circle, vertical tick and red cross show the mean, median, and mode, respectively. The solid grey curve shows the density of the posterior distribution for the 'null expectation' (see below).

*Models with randomised identities*

To obtain posterior distributions of $V_A$ and $V_A^k$ under the expectation of zero additive genetic variance, we ran our main quantitative genetic models (main Equations 2-3) after randomising the individual identities of the capture-resighting histories in the dataset. By doing so, any pattern of resemblance between individuals was therefore occurring purely by chance and not because of inherited genetic variation. As expected, these ‘null expectation’ analyses yielded negligible estimates of $V_{\mathrm{A}}$ and $V_{\mathrm{A}}^{k}$, with posterior distributions that clearly peaked on zero and have low density on large values (Figure S9).

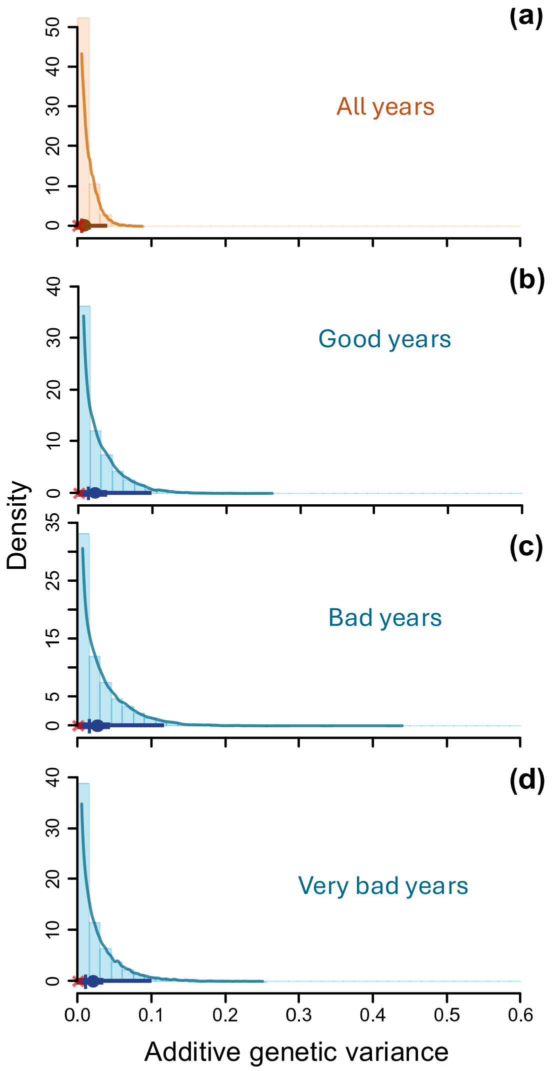


**Figure S9.** Posterior distributions of additive genetic variance underlying annual adult survival estimated (a) across all years ($V_{\mathrm{A}}$), and separately in years with (b) good, (c) bad, and (c) very bad environmental conditions ($V_{\mathrm{A}}^{k}$), after randomising the individual identities of the capture-resighting histories. Histogram bars show the distribution of the posterior samples, with corresponding kernel density estimation in solid line. The superimposed thin and thick horizontal lines show the 95% and 50% credible intervals, respectively. Along these horizontal lines, the plain circle, vertical tick and red cross show the mean, median, and mode, respectively.

##### S3.5 Simulations evaluating our capability to detect $V_\mathrm{A}$

To evaluate our capability to detect $V_\mathrm{A}$, we generated 'replicated' datasets by simulating phenotypic outcomes of a heritable dichotomous trait expressed in three different environmental categories, given the structure of the present resighting and relatedness data analysed in our study. We replicated all our focal individuals, with the same pedigree, and approximately the same amount of informative phenotypic outcomes $z$ in each environmental category $k$ per individual $i$. More precisely, for each $i$, we simulated as many $z$ values in $k$ as the number of years in $k$ when $i$ was known to be alive in the real dataset (see Appendix S1). Given the very high detection probabilities achieved by the shag monitoring on IoM (see main text and Appendix S4), this represents only slightly larger effective sample sizes compared to the real dataset.

To simulate and subsequently analyse $z$, we considered a generative model analogous to Equation (S45) above, i.e. the version of our main model including environment-specific additive genetic and permanent individual effects, but without age effects:

$$\mathrm{probit}\left(\tilde{\phi}_{i,t}\right) = \tilde{\mu} + \tilde{\gamma}_{s(i),t} + \tilde{b}_{k(t),i} + \tilde{a}_{k(t),i}, \qquad \text{(S46)}$$

where specifications are as for Equation (3) of the main text, yet '~' denotes the simulation (to distinguish the parameters from those of our main model fitted to the real data, i.e. Equation 3 of main text). To generate the data, we chose simulation values for $\tilde{\mu}_{s,t} = \tilde{\mu} + \tilde{\gamma}_{s(i),t}$, $\tilde{V}_B$ the variance in $\tilde{b}_{k(t),i}$, and $\tilde{V}_A^k$ the variance in $\tilde{a}_{k,i}$ (see below). We drew $\tilde{b}_{k(t),i}$ randomly from $\mathcal{N}\left(0, \tilde{V}_\mathrm{B}^k\right)$ and $\tilde{\boldsymbol{a}}_{\boldsymbol{k}}$ from $\mathcal{N}\left(0, \boldsymbol{A}\tilde{V}_\mathrm{A}^k\right)$. We then calculated each $\tilde{\phi}_{i,t} = \mathrm{probit}^{-1}\left(\tilde{\mu} + \tilde{\gamma}_{s(i),t} + \tilde{b}_i + \tilde{a}_{k(t),i}\right)$ and drew $z_{i,t}$ from $Bernoulli(\tilde{\phi}_{i,t})$.

We avoided simulating phenotypic expression explicitly as survival *vs* death, since removing individuals after death given simulated parameter values would have necessarily distorted the observation structure and pedigree informativeness relative to our original dataset. The replicated datasets were generated given simulated probit-scale intercepts corresponding to the sex×year specific estimates of survival probability obtained from our analyses of IoM shags. Hence, our simulations replicate the same phenotypic frequencies as for survival (vs death) in the shag dataset. Noting that our animal model inferences are blind to whether phenotypic expression is actually reversible or not, this procedure provided an appropriate approach for replicating datasets with the same structure and expected informativeness as our real data.

Accordingly, we simulated 50 datasets for the three scenarios defined by the simulation value of $\tilde{V}_A^k$ (with identical $\tilde{V}_A^k$ for all three environmental categories): $\tilde{V}_A^k = 0.07$, $\tilde{V}_A^k = 0.14$, and $\tilde{V}_A^k = 0.28$, i.e.$\vartheta/2$, $\vartheta$, and $2\vartheta$, where $\vartheta = 0.14$ is the posterior mean of $V_{\mathrm{A}}^{good}$ from our main model. For $\tilde{V}_B$, we always used the simulation value $\tilde{V}_{\mathrm{B}}^k = 0.04$, i.e. the posterior mean of $V_{\mathrm{B}}$ from our main quantitative genetic model fitted to the real dataset (Appendix S4). And for $\tilde{\mu}_{s,t}$, we always used values such that $E(\tilde{\phi}_{i,t})$ would be equal to the posterior mean $\hat{\phi}_{s,t}$ of the sex×year specific survival probability estimated from our initial capture-recapture analyses (Figure 3 of main text). Accordingly, we calculated simulation values as $\tilde{\mu}_{s,t} = \mathrm{probit}^{-1}(\hat{\phi}_{s,t})\sqrt{\tilde{V}_B + \tilde{V}_{\mathrm{A}}^k + 1}$.

We then fitted a simple binary generalised animal model (de Villemereuil et al., 2016) enacting equation (S42) to each dataset, thereby revealing our capability to detect $\tilde{V}_A^k$ given each scenario (and hence $V_{\mathrm{A}}^k$, by analogy). Graphical representations of all posterior distributions of each $\tilde{V}_A^k$ for all three simulation scenarios and replicated datasets are available via the data archive for our present study (Acker et al., 2026). Overall, across all replicated datasets for all simulation scenarios, the simulated values were most often included within the 95% CRI for the corresponding estimate of $\tilde{V}_A^k$. And no systematic bias was apparent across all cases for which there was evidence of non-zero $\tilde{V}_A^k$ (i.e. a posterior mode away from zero), as the relative frequency of posterior means that were greater vs lower than the focal simulated value was approximately 50%.

Detailed inspection of the posterior distributions showed that our data structure should provide relatively good capability to detect non-negligible $V_{\mathrm{A}}^k$ in each environmental category when the simulated value was $\vartheta = 0.14$. For the fifteen good years, nearly all posterior distributions (47/50) peaked away from zero. Yet, some distributions (8/50) were downward-biased by another (usually lower-density) peak on zero, implying equivocal conclusions. For the six bad years, our capability to detect $V_{\mathrm{A}}^k$ remained decent, with most posterior distributions

(37/50) peaking away from zero, but equivocal conclusions were more frequent. For the three very bad years, uncertainty was much greater and fewer distributions (23/50) peaked away from zero. Meanwhile, when the simulated value was halved ($\vartheta/2 = 0.07$), our simulation analyses still demonstrated capability to detect non-negligible $V_{\mathrm{A}}^{k}$ but it was more restricted overall. Here, posterior distributions peaked away from zero in only 33/50, 18/50, and 6/50 cases for good, bad, and very bad years, respectively. In contrast, doubling the simulated value ($2\vartheta = 0.28$) yielded near-complete capability to detect non-negligible $V_{\mathrm{A}}^{k}$ in good and bad years. Specifically, all posterior distributions peaked away from zero, and the downward bias was limited overall. In very bad years, only 10/50 distributions had a mode on zero, yet distribution had large uncertainty (with 95%CIs widely overlapping small values).

##### S3.6 Posterior sampling and derived calculations

*Posterior sampling, diagnostics and archive*

For each of the primary models presented in the main text (i.e. the CJS capture-recapture model, and the capture-recapture animal model defined by Equations 1-3), posterior sampling was performed using 4 Markov chains with 1,000 warmup iterations followed by 5,000 monitored iterations (with no thinning), yielding 20,000 samples in total (details on specificities of the sampling protocol in Stan are in Stan Development Team, 2026; see also McElreath, 2020). No divergent iterations occurred during sampling of the models. Here, visual inspection of the chains and the Gelman-Rubin-Brooks $\hat{R}$ statistics (Brooks & Gelman, 1998) calculated following Vehtari et al. (2021) indicated good convergence, with $\hat{R}$<1.01 for all model parameters (and most had $\hat{R} \approx 1$). Bulk and tail effective sample sizes ('ESS') were >400 for all parameters (and most had much higher ESS), suggesting no sampling issues and good reliability of calculated $\hat{R}$ values (Vehtari et al., 2021). We therefore achieved estimation with negligible uncertainty due to imperfect (pseudorandom) sampling. More specifically, the Monte Carlo standard error of the mean (calculated following Vehtari et al., 2021) was <0.005 for all parameters and typically smaller, implying that posterior means could be safely reported with 2-3 decimal place precision.

Overall, these diagnostics indicated good sampling efficiency allowing reliable and precise inference (Gelman et al., 2013; McElreath, 2020; Vehtari et al., 2021). Detailed summaries and diagnostics of the posterior distributions of all parameters and derived quantities from the main models are available alongside complete posterior samples via the data archive for our study (Acker et al., 2026).

For all alternative or additional models, we used 4 Markov chains with 1000 warmup iterations followed by 3000 monitored iterations (with no thinning), yielding 12,000 samples in total. The diagnostics were slightly less optimal yet still very satisfactory for current purposes. We achieved good convergence with all $\hat{R}$<1.02 (and $\hat{R} \approx 1$ for most parameters), ESS>200 (and ESS>1000 or much larger for most parameters), and only occasional divergent iterations.

*Derived calculations*

To clarify the evolutionary implications of our main inferences, we calculated quantities derived from the model parameters (which are the primary target of estimation). Note that any derived parameter is calculated from the set of primary model parameter values in each posterior draw (i.e. each iteration of the MCMC chains). The collection of derived parameter values calculated in all posterior draws constitutes the set of posterior samples of the derived parameter, used to approximate its posterior distribution.

More specifically, we derived year-specific population mean survival probabilities $\bar{\phi}_t$ from the CJS model parameters using Equation (S41). We derived sex×year×age-specific phenotypic-scale genotypic variance in survival $V_{\phi_{s,t,x}|\mathrm{A}}$ from the animal model parameters using Equation (S9). And, we used this quantity to derive sex×year×age-specific (proportional) partial responses to selection $V_{\phi_{s,t,x}|\mathrm{A}}/\bar{\phi}_{s,t,x}$ (and $V_{\phi_{s,t,x}|\mathrm{A}}/\bar{\phi}_{s,t,x}{}^{2}$), after deriving $\bar{\phi}_{s,t,x}$ using Equation (S7). The latent-scale heritabilities $h^2$ were calculated using the formula provided in main text. And finally, the probabilities $P(V_{\mathrm{A}}^{m} > V_{\mathrm{A}}^{n})$ for measuring the support for differences in $V_{\mathrm{A}}^{k}$ (and hence in $h_k^2$) between pairs of environments $\{m, n\}$ were calculated as the proportion of posterior draws in which $V_{\mathrm{A}}^{m}$ was greater than $V_{\mathrm{A}}^{n}$ (Gelman et al., 2013).

### Appendix S4

### Additional details of the results

#### S4.1 Sex- and age-specific survival

In the main text, we provided graphical summaries for the adult annual survival probabilities estimated using the sex×year-specific capture-recapture CJS model (Figure 3). Hereafter, we provide the corresponding numerical summaries (Table S5).

**Table S5**. Estimates of the sex×year-specific survival probabilities and year-specific population mean survival obtained from the capture-recapture CJS model.

| Year | Posterior summary | | | | | |
|---|---|---|---|---|---|---|
| | Females | | Males | | Full population | |
| | Mean | 95% CRI | Mean | 95% CRI | Mean | 95% CRI |
| 2000 | 0.962 | [0.897, 0.997] | 0.950 | [0.883, 0.995] | 0.955 | [0.911, 0.988] |
| 2001 | 0.926 | [0.881, 0.962] | 0.918 | [0.876, 0.955] | 0.922 | [0.891, 0.949] |
| 2002 | 0.944 | [0.897, 0.984] | 0.922 | [0.880, 0.958] | 0.933 | [0.902, 0.960] |
| 2003 | 0.984 | [0.944, 1.000] | 0.982 | [0.943, 0.999] | 0.983 | [0.957, 0.997] |
| 2004 ⦿ | 0.266 | [0.216, 0.321] | 0.549 | [0.494, 0.607] | 0.413 | [0.375, 0.453] |
| 2005 ⊗ | 0.821 | [0.719, 0.915] | 0.878 | [0.818, 0.932] | 0.858 | [0.806, 0.907] |
| 2006 ⊗ | 0.658 | [0.588, 0.726] | 0.735 | [0.682, 0.786] | 0.704 | [0.662, 0.745] |
| 2007 ⊗ | 0.811 | [0.757, 0.859] | 0.837 | [0.793, 0.877] | 0.826 | [0.792, 0.857] |
| 2008 | 0.916 | [0.880, 0.946] | 0.926 | [0.896, 0.952] | 0.922 | [0.899, 0.942] |
| 2009 | 0.948 | [0.922, 0.970] | 0.929 | [0.902, 0.953] | 0.938 | [0.919, 0.955] |
| 2010 | 0.927 | [0.898, 0.952] | 0.931 | [0.904, 0.953] | 0.929 | [0.910, 0.946] |
| 2011 | 0.910 | [0.871, 0.951] | 0.936 | [0.907, 0.963] | 0.923 | [0.899, 0.948] |
| 2012 ⦿ | 0.348 | [0.309, 0.387] | 0.522 | [0.483, 0.563] | 0.436 | [0.408, 0.464] |
| 2013 ⊗ | 0.698 | [0.644, 0.749] | 0.803 | [0.761, 0.842] | 0.757 | [0.723, 0.788] |
| 2014 | 0.935 | [0.905, 0.961] | 0.928 | [0.899, 0.954] | 0.932 | [0.911, 0.950] |
| 2015 | 0.930 | [0.900, 0.956] | 0.857 | [0.821, 0.889] | 0.890 | [0.867, 0.912] |
| 2016 | 0.912 | [0.877, 0.943] | 0.914 | [0.880, 0.944] | 0.913 | [0.890, 0.935] |
| 2017 ⊗ | 0.713 | [0.668, 0.756] | 0.753 | [0.710, 0.793] | 0.733 | [0.703, 0.763] |
| 2018 ⊗ | 0.824 | [0.783, 0.862] | 0.856 | [0.819, 0.889] | 0.841 | [0.814, 0.867] |
| 2019 | 0.921 | [0.887, 0.952] | 0.895 | [0.862, 0.924] | 0.907 | [0.884, 0.929] |
| 2020 | 0.910 | [0.876, 0.939] | 0.902 | [0.869, 0.932] | 0.906 | [0.883, 0.927] |
| 2021 | 0.932 | [0.904, 0.957] | 0.902 | [0.870, 0.931] | 0.917 | [0.896, 0.936] |
| 2022 | 0.970 | [0.925, 0.999] | 0.933 | [0.882, 0.989] | 0.952 | [0.917, 0.984] |
| 2023 ⦿ | 0.179 | [0.146, 0.217] | 0.185 | [0.154, 0.219] | 0.183 | [0.159, 0.207] |

Notes: Posterior medians and modes were all virtually identical to the posterior means. Symbols '⊗' and '⦿' indicate bad and very bad years, respectively.

We also provide a graphical representation of the relationship between age and survival in each type of environment (Figure S10), showing predictions from our model decomposing the variance with a random effect of individual identity (main Equation 1). Note that, as expected, this result was virtually identical in the quantitative genetic models (main Equations 2-3; see below). There was clear evidence of strong actuarial senescence, which phenotypic expression depended on environmental conditions because of the non-linear genotype–environment–phenotype relationship (main Figure 2). Under ‘average’ conditions (corresponding to the upper end of bad years), survival probability of an average adult increases from ~0.75 to ~0.8 from age 3 to 7-8 years then slightly decreases to ~0.75 until age 11-12 years and then sharply decreases to ~0.1 at age 25 (the oldest ever observed in our population; Figure S10a). Under very good conditions, survival probability remained high ~0.95 from age 3 to 14 and declined to ~0.4 at age 25 (Figure S10b). Under very bad conditions, since survival probability was consistently lower, it declined from younger ages and approached zero at age 25 (Figure S10c).

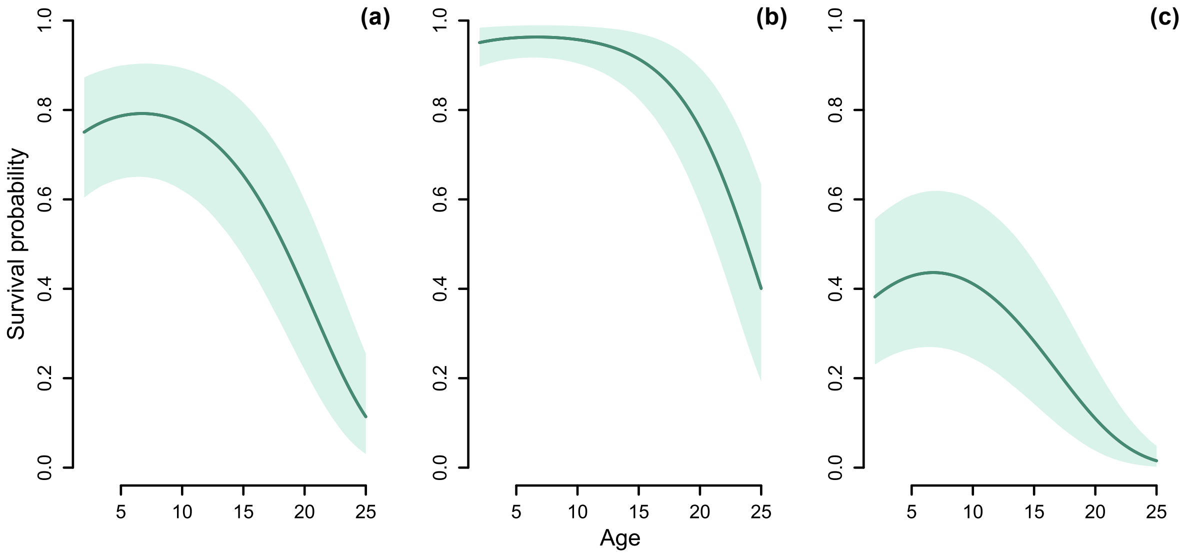


**Figure S10**. Relationship between survival probability and age for average adult shags in (a) an average year, (b) a hypothetical year with very good conditions, and (c) a hypothetical year with very bad conditions. The relationship was calculated for each posterior draw of the model defined in Equation (1) of the main text, given $\phi_{i,x} = \text{probit}^{-1}(\mu + \beta_1 x + \beta_2 x^2 + \gamma)$ (with $c_i = 0$). Values for $\gamma$ were selected to represent the desired environmental severity: $\gamma = 0$, $\gamma = 1$, and $\gamma = -1$ for panels a, b, and c, respectively. See numerical summaries of the parameter values below. On each panel, the solid line indicates the posterior mean, and the shaded area indicates the 95% CRI.

#### S4.2 Detection probabilities

Hereafter we provide graphical summaries of the posterior distribution of the resighting probabilities depending on year, sex, and ring type (Figure S11). The probabilities presented here are those estimated from our initial capture recapture CJS model. Yet, these parameter estimates remained virtually identical across all models. Overall, the probability of resighting individuals that had a colour ring was typically very high and often slightly higher in males (Figure S11a). Meanwhile, the probability of resighting individuals that had only a metal ring was much lower (Figure S11b) and estimates were also less precise (as expected given the low number of individuals that had only a metal ring, notably in recent years; see Appendix S1).

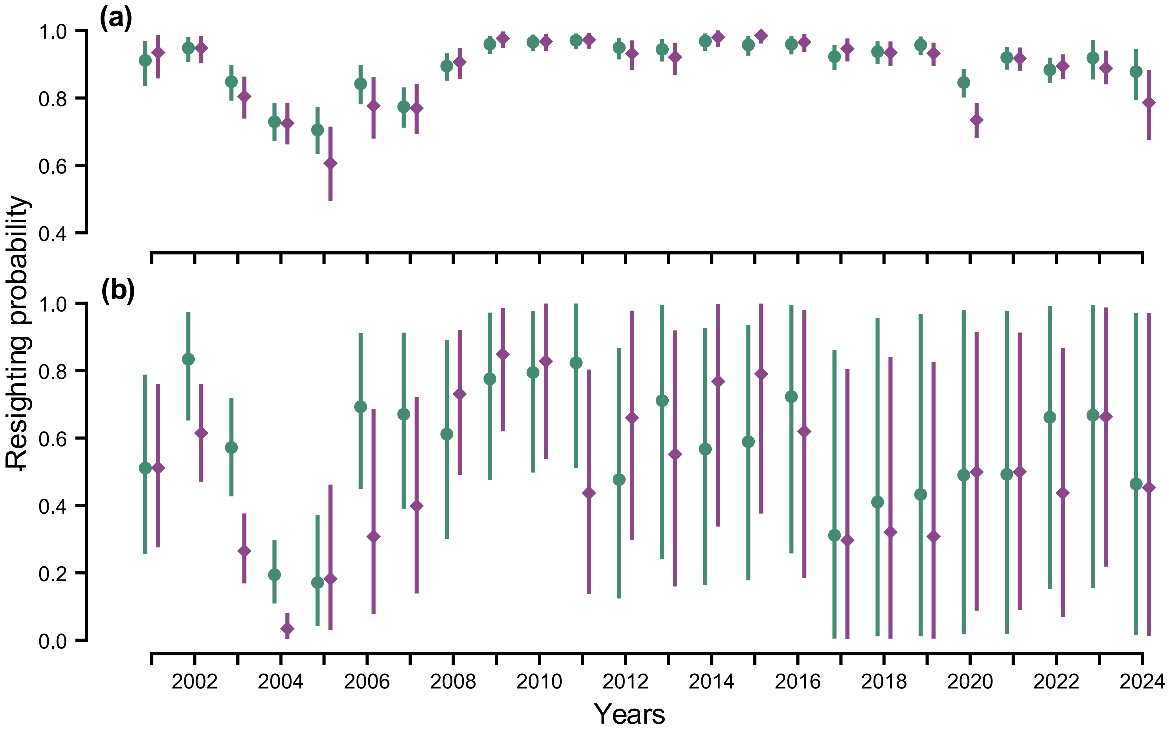


**Figure S11**. Estimates of the sex×year-specific resighting probabilities for individuals with (a) a colour ring or (b) a metal ring only. Points and error bars indicate posterior means and 95% credible intervals. Green circles and purple diamonds represent males and females, respectively.

#### S4.3 Probit regression parameters

Hereafter we provide numerical summaries of the probit-scale parameters of our main quantitative genetic analyses defined by Equations (1-3) of the main text (Table S6-S8).

**Table S6**. Parameter estimates of the individual heterogeneity model (main text Equation 1).

| Parameter | Posterior summary | | |
|---|---|---|---|
| | Mean | Median | 95% CRI |
| $\mu$ | 0.924 | 0.923 | [0.682, 1.175] |
| $\beta_1$ | 0.085 | 0.086 | [0.056, 0.114] |
| $\beta_2$ | -0.006 | -0.006 | [-0.008, -0.005] |
| $V_{\gamma,♀}$ | 0.778 | 0.727 | [0.405, 1.455] |
| $V_{\gamma,♂}$ | 0.450 | 0.421 | [0.238, 0.837] |
| $V_C$ | 0.058 | 0.054 | [0.001, 0.138] |

**Table S7**. Parameter estimates of the overall animal model (main text Equation 2).

| Parameter | Posterior summary | | |
|---|---|---|---|
| | Mean | Median | 95% CRI |
| $\mu$ | 0.947 | 1.945 | [0.698, 1.207] |
| $\beta_1$ | 0.086 | 0.086 | [0.056, 0.114] |
| $\beta_2$ | -0.006 | -0.006 | [-0.008, -0.005] |
| $V_{\gamma,♀}$ | 0.800 | 0.748 | [0.421, 1.467] |
| $V_{\gamma,♂}$ | 0.459 | 0.429 | [0.239, 0.847] |
| $V_A$ | 0.035 | 0.033 | [0.001, 0.081] |
| $V_B$ | 0.034 | 0.025 | [0.000, 0.110] |

**Table S8**. Parameter estimates of the environment-specific animal model (main text Equation 3).

| Parameter | Posterior summary | | |
|---|---|---|---|
| | Mean | Median | 95% CRI |
| $\mu$ | 0.918 | 0.917 | [0.666, 1.177] |
| $\beta_1$ | 0.089 | 0.089 | [0.059, 0.118] |
| $\beta_2$ | -0.007 | -0.007 | [-0.008, -0.005] |
| $V_{\gamma,♀}$ | 0.835 | 0.778 | [0.436, 1.547] |
| $V_{\gamma,♂}$ | 0.473 | 0.442 | [0.245, 0.883] |
| $V_{\mathrm{A}}^{good}$ | 0.138 | 0.13 | [0.007, 0.316] |
| $V_{\mathrm{A}}^{bad}$ | 0.029 | 0.016 | [0.000, 0.122] |
| $V_{\mathrm{A}}^{vbad}$ | 0.063 | 0.046 | [0.000, 0.220] |
| $V_B$ | 0.041 | 0.034 | [0.000, 0.123] |

**S4.4 Phenotypic-scale consequences**

In Figure 5 of the main text, we showed the posterior distributions of $V_{\phi_{s,t,x}|\mathrm{A}}$, $V_{\phi_{s,t,x}|\mathrm{A}}/\bar{\phi}_{s,t,x}$, $V_{\phi_{s,t,x}|A}/\bar{\phi}_{s,t,x}{}^{2}$ as a function of sex-specific environmental severity ($1-\bar{\phi}_{s,t,x}$), for age 7 years (i.e. $x=7$). To facilitate readability, we did not represent the uncertainties in $1-\bar{\phi}_{s,t,x}$ along the y-axis. Hereafter, we show the estimates of the same parameters along with uncertainties in $1-\bar{\phi}_{s,t,x}$, for age 3 (Figure S12), age 7 (Figure S13, i.e. replicate of Figure 5), and age 15 (Figure

S14). Importantly, these figures show that patterns were largely similar across ages, although senescence (and hence lower mean) at older ages resulted in phenotypic-scale genotypic variance that were larger in good years and smaller in very bad years, compared to younger ages. Moreover, uncertainties in $1 - \bar{\phi}_{s,t,x}$ do not challenge the conclusions already visible on Figure 5.

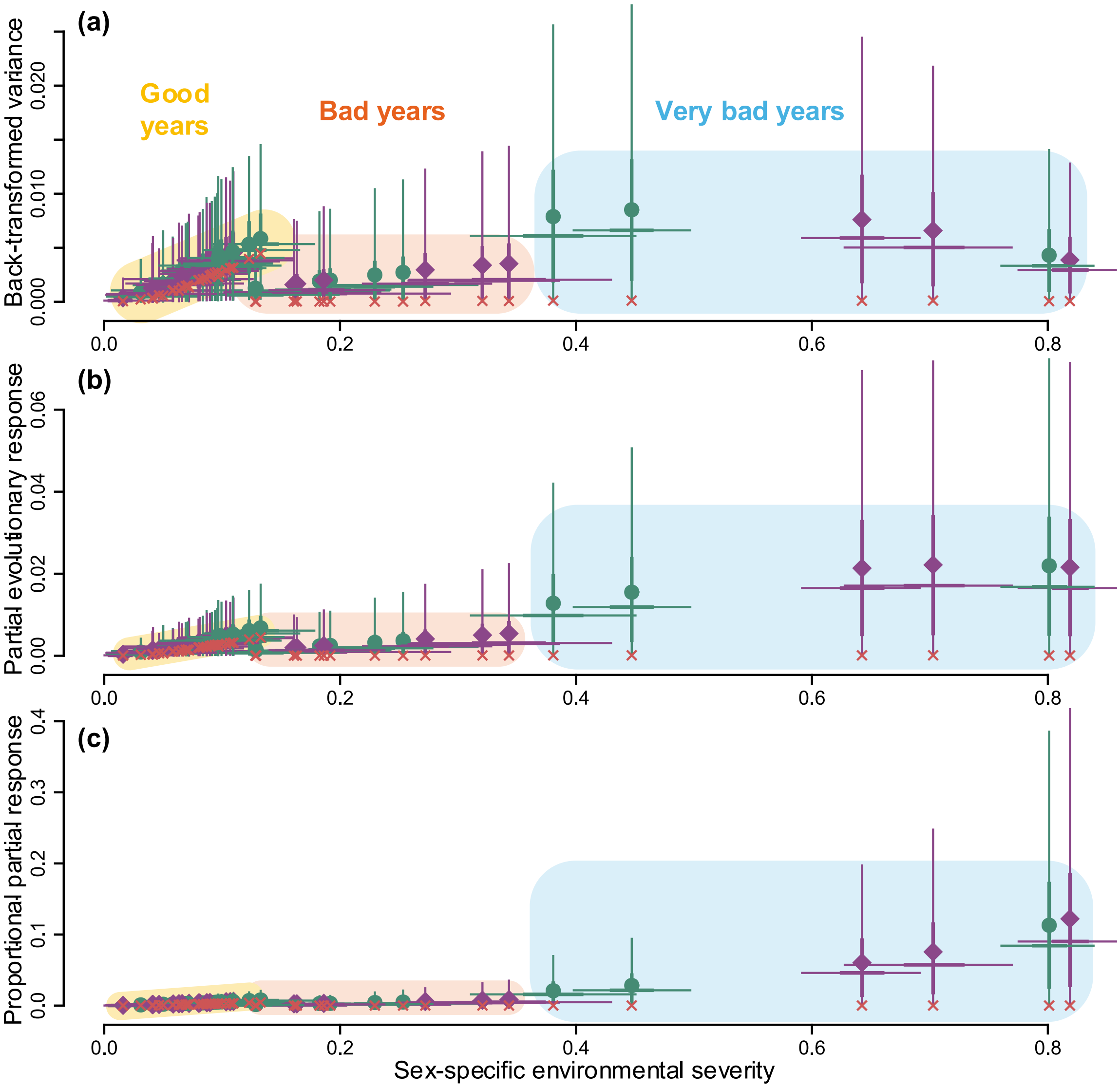


**Figure S12**. Phenotypic-scale consequences of the latent-scale additive genetic variance underlying annual adult survival. (a) The sex×year-specific genotypic variance in survival conditional on age 3 years, $V_{\phi_{s,t,3}|\mathrm{A}}$. (b) The partial evolutionary response to selection, $V_{\phi_{s,t,3}|\mathrm{A}}/\bar{\phi}_{s,t,3}$. (c) The partial evolutionary response, $V_{\phi_{s,t,3}|\mathrm{A}}/\bar{\phi}_{s,t,3}^{\ 2}$. In all panels, environmental severity is defined as $1 - \bar{\phi}_{s,t,3}$. Green circles and purple diamonds indicate posterior means of males and females, respectively. Thick and thin error bars show the 50% and 95% credible intervals for the focal phenotypic-scale parameter and environmental severity (vertical and horizontal bars, crossing at the posterior median). Red crosses indicate posterior modes.

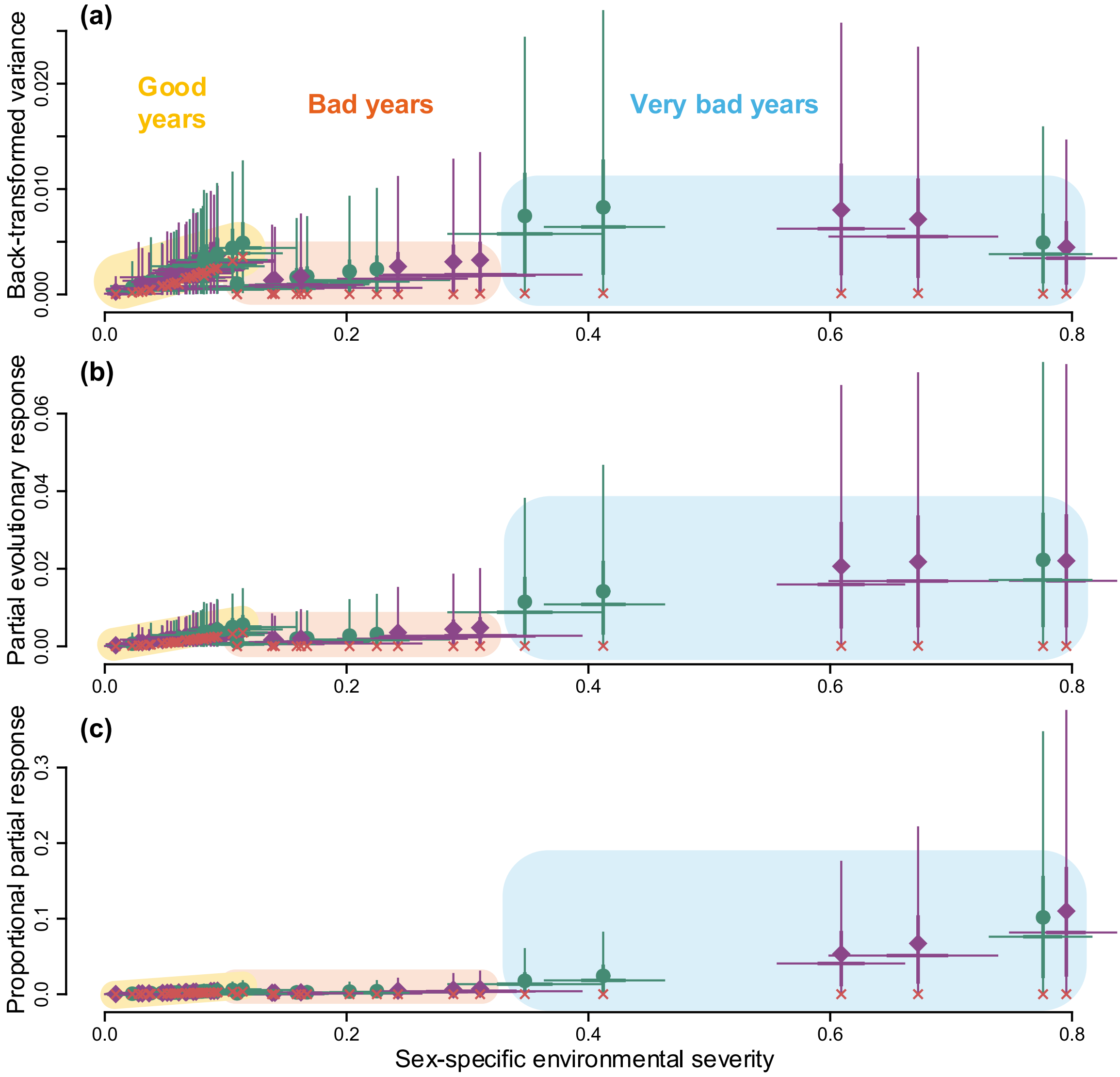


**Figure S13.** Phenotypic-scale consequences of the latent-scale additive genetic variance underlying annual adult survival. (a) The sex×year-specific genotypic variance in survival conditional on age 7 years, $V_{\phi_{s,t,7}|\mathrm{A}}$. (b) The partial evolutionary response to selection, $V_{\phi_{s,t,7}|\mathrm{A}}/\bar{\phi}_{s,t,7}$. (c) The partial evolutionary response, $V_{\phi_{s,t,7}|\mathrm{A}}/\bar{\phi}_{s,t,7}^{\,2}$. In all panels, environmental severity is defined as $1-\bar{\phi}_{s,t,7}$. See Figure S12 for other legend details. Note, however, that the x-axis range is wider in Figure S13 than in Figure S12.

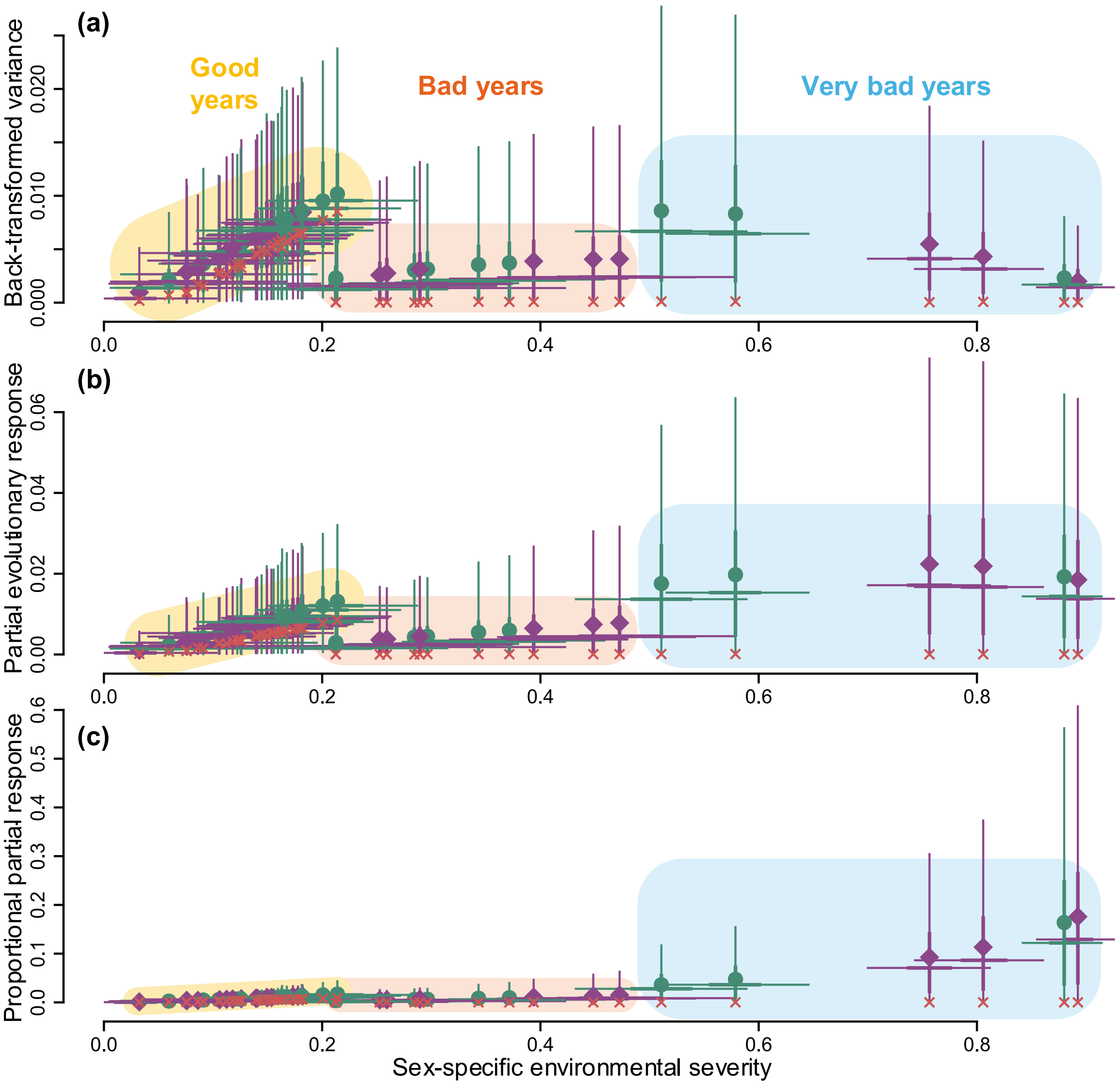


**Figure S14**. Phenotypic-scale consequences of the latent-scale additive genetic variance underlying annual adult survival. (a) The sex×year-specific genotypic variance in survival $V_{\phi_{s,t,15}|\mathrm{A}}$ for age 15 years. (b) The partial evolutionary response to selection $V_{\phi_{s,t,15}|\mathrm{A}}/\bar{\phi}_{s,t,15}$. (c) The partial evolutionary response $V_{\phi_{s,t,15}|\mathrm{A}}/\bar{\phi}_{s,t,15}{}^{2}$. In all panels, environmental severity is defined as $1-\bar{\phi}_{s,t,15}$. See Figure S12 for other legend details. Note, however, that the x-axis range is wider in Figure S14 than in Figure S12.

#### S4.5 Alternative quantitative genetic models

*Models with alternative grouping of environmental categories*

In addition to our main model (main text Equation 3), we tested two models with alternative categories. Results are shown on Figure S15 hereafter.

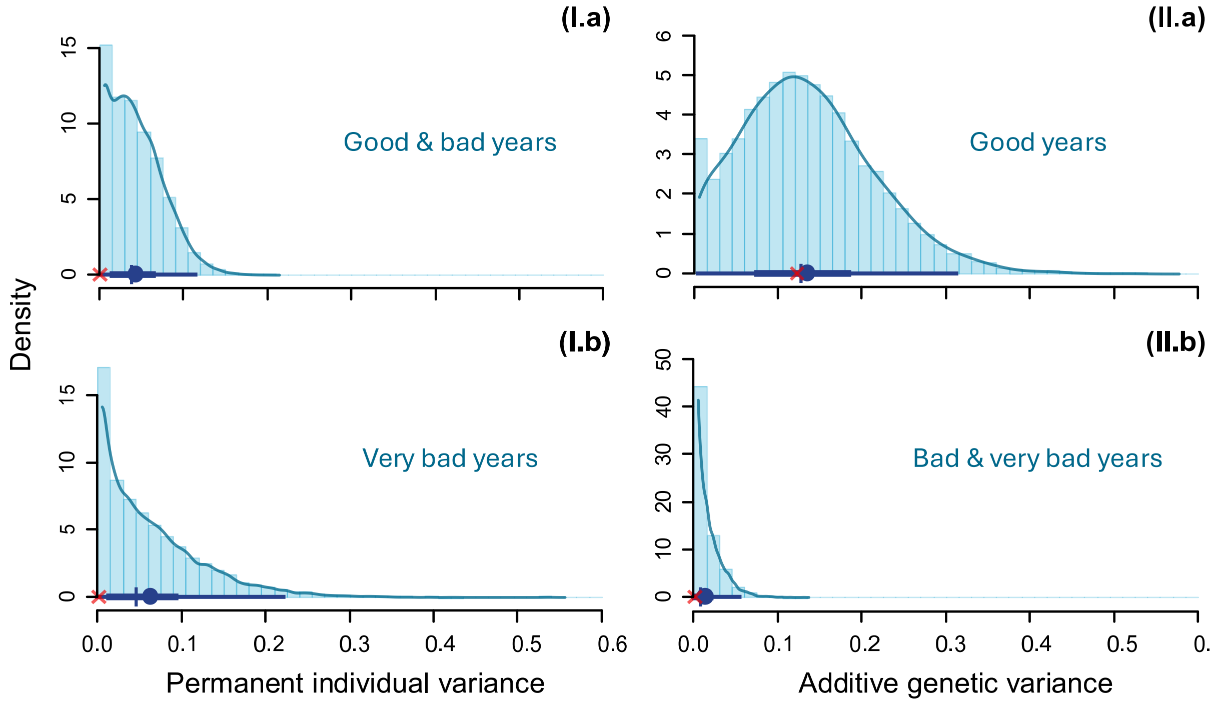


**Figure S15**. Posterior distributions of the environment-specific additive genetic variances underlying annual adult survival ($V_{\mathrm{A},k}$) under (I.a) good & bad vs (I.b) very bad environmental conditions, or (II.b) good vs (II.b) bad & very bad environmental conditions. Histogram bars show the distribution of the posterior samples, with corresponding kernel density estimation in solid line. The superimposed thin and thick horizontal lines show the 95% and 50% credible intervals, respectively. Along these horizontal lines, the plain circle, vertical tick and red cross show the mean, median, and mode, respectively.

In the model where good and bad years were grouped together (vs very bad years only), we found $V_{\mathrm{A}}^{good\,\&\,bad} = 0.043\ [0.000, 0.114]$ and $V_{\mathrm{A}}^{vbad} = 0.063\ [0.000, 0.222]$. Hence, the estimate for $V_{\mathrm{A}}^{good\,\&\,bad}$ was similar to the estimate for $V_A$ in the overall animal model (Equation 2 of main text); and the estimate of $V_{\mathrm{A}}^{vbad}$ was unchanged as expected. In the other model, where bad and very bad years were grouped together (vs good years only), we found $V_{\mathrm{A}}^{good} = 0.135\ [0.005, 0.312]$ and $V_{\mathrm{A}}^{vbad} = 0.014\ [0.000, 0.055]$. Hence, for $V_{\mathrm{A}}^{bad\,\&\,vbad}$ was similar to the estimate for $V_{\mathrm{A}}^{bad}$ in the original version of the main animal model (Equation 3 of main text); and the estimate of $V_{\mathrm{A}}^{good}$ was unchanged as expected.

*Models with cross-environment genetic correlations*

We also tested for a model similar to our main model (Equation 3 of main text), but that further included explicit correlations between pairs of environments $\{m, n\}$, $\rho_{m,n}$. More specifically, this implies expressing the joint distribution of the additive genetic effects in the three environments as follows:

$$\vec{\boldsymbol{a}}_{\mathbf{1:3}} \sim \mathcal{N}(0, \boldsymbol{G} \otimes \boldsymbol{A} V_{\mathrm{A}}^{k}). \tag{S47}$$

Here, $\vec{\boldsymbol{a}}_{\mathbf{1:3}}$ is the vertically stacked vector of vectors each environment-specific vector of breeding values $\boldsymbol{a_k}$ (with indexes 1, 2, and 3 referring to the good, bad, and very bad environment, respectively) and $\boldsymbol{G}$ is the cross-environment genetic covariance matrix (and $\otimes$ denotes the Kronecker product). More precisely:

$$\vec{\boldsymbol{a}}_{\mathbf{1:3}} = \begin{bmatrix} \boldsymbol{a_1} \\ \boldsymbol{a_2} \\ \boldsymbol{a_3} \end{bmatrix}, \tag{S48}$$

$$\boldsymbol{G} = \begin{bmatrix} V_{\mathrm{A}}^{good} & \rho_{1,2}\sqrt{V_{\mathrm{A}}^{good} V_{\mathrm{A}}^{bad}} & \rho_{1,3}\sqrt{V_{\mathrm{A}}^{good} V_{\mathrm{A}}^{vbad}} \\ \rho_{1,2}\sqrt{V_{\mathrm{A}}^{good} V_{\mathrm{A}}^{bad}} & V_{\mathrm{A}}^{bad} & \rho_{2,3}\sqrt{V_{\mathrm{A}}^{bad} V_{\mathrm{A}}^{vbad}} \\ \rho_{1,3}\sqrt{V_{\mathrm{A}}^{good} V_{\mathrm{A}}^{vbad}} & \rho_{2,3}\sqrt{V_{\mathrm{A}}^{bad} V_{\mathrm{A}}^{vbad}} & V_{\mathrm{A}}^{vbad} \end{bmatrix}. \tag{S49}$$

This alternative model revealed no evidence of any specific correlation between the three pairs of environments (Figure S16). The estimates for the cross-environment additive genetic correlations were 0.087 [-0.821, 0.880] for $\rho_{1,2}$ (between good and bad environments), 0.426 [-0.524, 0.937] for $\rho_{1,3}$ (between good and very bad environments), and -0.066 [-0.872, 0.841] for $\rho_{2,3}$ (between bad and very bad environments). The substantial uncertainties in these correlations are unsurprising given the estimated negligible $V_{\mathrm{A}}^{bad}$ and $V_{\mathrm{A}}^{vbad}$. Indeed, low variances in breeding values in the bad and very bad environments imply very little information for estimating associations between them or with breeding values in the good environment.

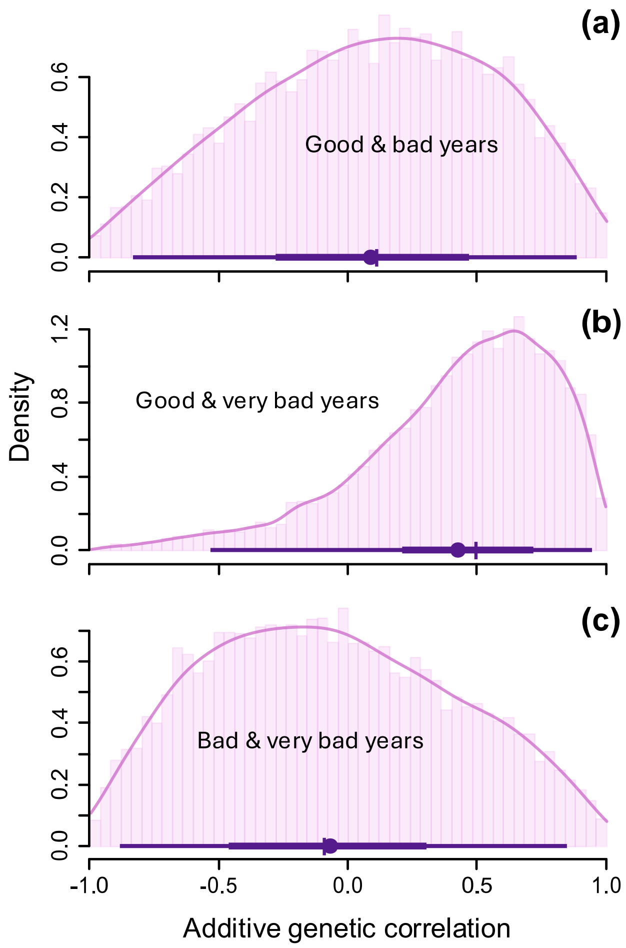


**Figure S16**. Posterior distributions of the additive genetic correlation underlying annual adult survival between (a) good & bad, (b) good & very bad, and (c) bad & very bad environmental conditions ($\rho_{1,2}$, $\rho_{1,3}$, and $\rho_{2,3}$, respectively). Histogram bars show the distribution of the posterior samples, with corresponding kernel density estimation in solid line. The superimposed thin and thick horizontal lines show the 95% and 50% credible intervals, respectively. Along these horizontal lines, the plain circle and vertical tick show the mean and median, respectively.

Yet, if anything, there was no evidence for a strongly negative correlation between the good and very bad environments (Figure S16b). Moreover, posterior distributions of $V_{\mathrm{A}}^{k}$ in each environmental category was similar to those in the main model (Figure S17); with $V_{\mathrm{A}}^{good} = 0.139\ [0.011, 0.304]$, $V_{\mathrm{A}}^{bad} = 0.034\ [0.000, 0.116]$, and $V_{\mathrm{A}}^{vbad} = 0.075\ [0.000, 0.234]$.

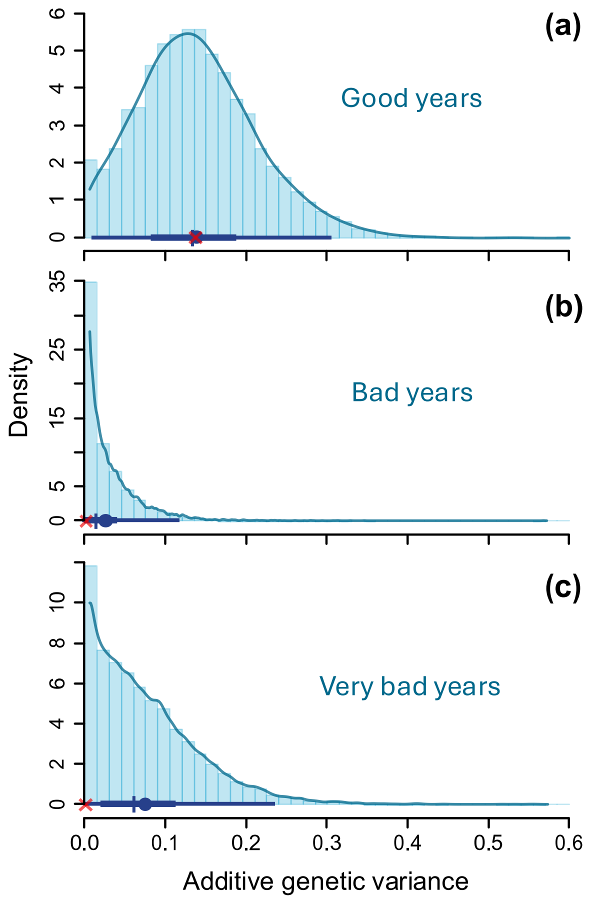


**Figure S17**. Posterior distributions of the additive genetic variances underlying annual adult survival ($V_{\mathrm{A},k}$) under (a) good, (b) bad, and (c) very bad environmental conditions, from the model with cross-environment genetic correlations. Histogram bars show the distribution of the posterior samples, with kernel density estimation in solid line. The superimposed thin and thick horizontal lines show the 95% and 50% credible intervals, respectively. Along these horizontal lines, the plain circle, vertical tick and red cross show the mean, median, and mode, respectively.

*Sex-specific quantitative genetic models*

We also tested for models expanding Equations 2 and 3 of the main text to include sex-specific additive genetic variance $V^k_{\mathrm{A}♀}$ and $V^k_{\mathrm{A}♂}$ and a(n environment-specific) cross-sex correlation in the additive genetic effects, $\rho_{♀,♂}$ or $\rho^k_{♀,♂}$. More specifically, this implies rewriting Equations 2-3 as:

$$\text{probit}(\phi_{i,t}) = \mu + \gamma_{s(i),t} + \beta_1 x_{i,t} + \beta_2 {x_{i,t}}^2 + b_i + a_{i,s(i)}, \quad \text{(S50)}$$

$$\text{probit}(\phi_{i,t}) = \mu + \beta_1 x_{i,t} + \beta_2 {x_{i,t}}^2 + \gamma_{s(i),t} + b_i + a_{k(t),i,s(i)}, \quad \text{(S51)}$$

where parameter specifications are as in the main text, except for the distribution of $a_{i,s(i)}$ and $a_{k(t),i,s(i)}$. Similarly as for the model with cross-environment genetic correlations, we express the joint distribution of the additive genetic effects in the two sexes as $\vec{\boldsymbol{a}}_{\{♀,♂\}} \sim \mathcal{N}(0, \boldsymbol{\Omega} \otimes \boldsymbol{A} V_{\mathrm{A}}^{k})$ and $\vec{\boldsymbol{a}}_{\boldsymbol{k}\{♀,♂\}} \sim \mathcal{N}(0, \boldsymbol{\Omega}_{\boldsymbol{k}} \otimes \boldsymbol{A} V_{\mathrm{A}}^{k})$, for Equation S50 and S51 respectively. Here, $\vec{\boldsymbol{a}}_{\{♀,♂\}}$ or $\vec{\boldsymbol{a}}_{\boldsymbol{k}\{♀,♂\}}$ is the vertically stacked vector of each sex-specific vector of breeding values and $\boldsymbol{\Omega}$ or $\boldsymbol{\Omega}_{\boldsymbol{k}}$ is the cross-sex genetic covariance matrix. More precisely:

$$\vec{\boldsymbol{a}}_{\{♀,♂\}} = \begin{bmatrix} \boldsymbol{a}_{♀} \\ \boldsymbol{a}_{♂} \end{bmatrix} \text{ and } \vec{\boldsymbol{a}}_{\boldsymbol{k}\{♀,♂\}} = \begin{bmatrix} \boldsymbol{a}_{\boldsymbol{k},♀} \\ \boldsymbol{a}_{\boldsymbol{k},♂} \end{bmatrix}, \tag{S52}$$

$$\boldsymbol{\Omega} = \begin{bmatrix} V_{\mathrm{A}♀} & \rho_{♀,♂}\sqrt{V_{\mathrm{A}♀}V_{\mathrm{A}♂}} \\ \rho_{♀,♂}\sqrt{V_{\mathrm{A}♀}V_{\mathrm{A}♂}} & V_{\mathrm{A}♂} \end{bmatrix} \text{ and } \boldsymbol{\Omega}_{\boldsymbol{k}} = \begin{bmatrix} V_{\mathrm{A}♀}^{k} & \rho_{♀,♂}^{k}\sqrt{V_{\mathrm{A}♀}^{k}V_{\mathrm{A}♂}^{k}} \\ \rho_{♀,♂}^{k}\sqrt{V_{\mathrm{A}♀}^{k}V_{\mathrm{A}♂}^{k}} & V_{\mathrm{A}♂}^{k} \end{bmatrix}. \tag{S53}$$

In practice, this implies envisaging survival in males and females as two distinct traits defined for all individuals but expressed only by those of the corresponding sex (see archived code: Acker et al., 2026).

Here, we found no strong evidence of a cross-sex genetic correlation neither across all environments, nor in any of each environment (Figure S18). Indeed, the estimates were largely uncertain and widely overlapping negative and positive values: 0.279 [-0.850, 0.972] for $\rho_{♀,♂}$ (overall model), 0.395 [-0.758, 0.978] for $\rho_{♀,♂}^{good}$, 0.001 [-0.940, 0.945] for $\rho_{♀,♂}^{bad}$, and -0.066 [-0.917, 0.965] for $\rho_{♀,♂}^{vbad}$. Such substantial uncertainties are again unsurprising given the negligible estimated variances in all environments (see below), which reflect both genuinely small variances and inevitably reduced statistical power, since sample sizes are approximately halved compared with analyses combining both sexes. If anything, there was little evidence for a strongly negative correlation between sexes in good years (Figure S18b).

There was also no clear evidence of any difference in additive genetic variance between the sexes. Likely due to the reduced statistical power, estimates of $V_{\mathrm{A}♀}$ and $V_{\mathrm{A}♂}$ (from the overall model) did not reveal any non-zero variance (0.039 [0.000, 0.107] and 0.038 [0.000, 0.115], respectively; Figure S19). The estimate of $V_{\mathrm{A}♂}^{good}$ was also equivocal (0.086 [0.000, 0.311]), but the estimate of $V_{\mathrm{A}♀}^{good}$ clearly revealed non-zero variance (0.206 [0.006, 0.555]; Figure 22). Yet the latter estimate of $V_{\mathrm{A}♀}^{good}$ was highly uncertain, and the posterior distributions of the two sexes largely overlapped, yielding $P\left(V_{\mathrm{A}♀}^{good} > V_{\mathrm{A}♂}^{good}\right) = 0.79$. Meanwhile, other estimates still revealed only low or no additive genetic variance (Figure S19); $V_{\mathrm{A}♀}^{bad}$: 0.046 [0.000, 0.212], $V_{\mathrm{A}♂}^{bad}$: 0.050 [0.000, 0.224], and $V_{\mathrm{A}♀}^{vbad}$: 0.120 [0.000, 0.544], $V_{\mathrm{A}♂}^{vbad}$: 0.050 [0.000, 0.222].

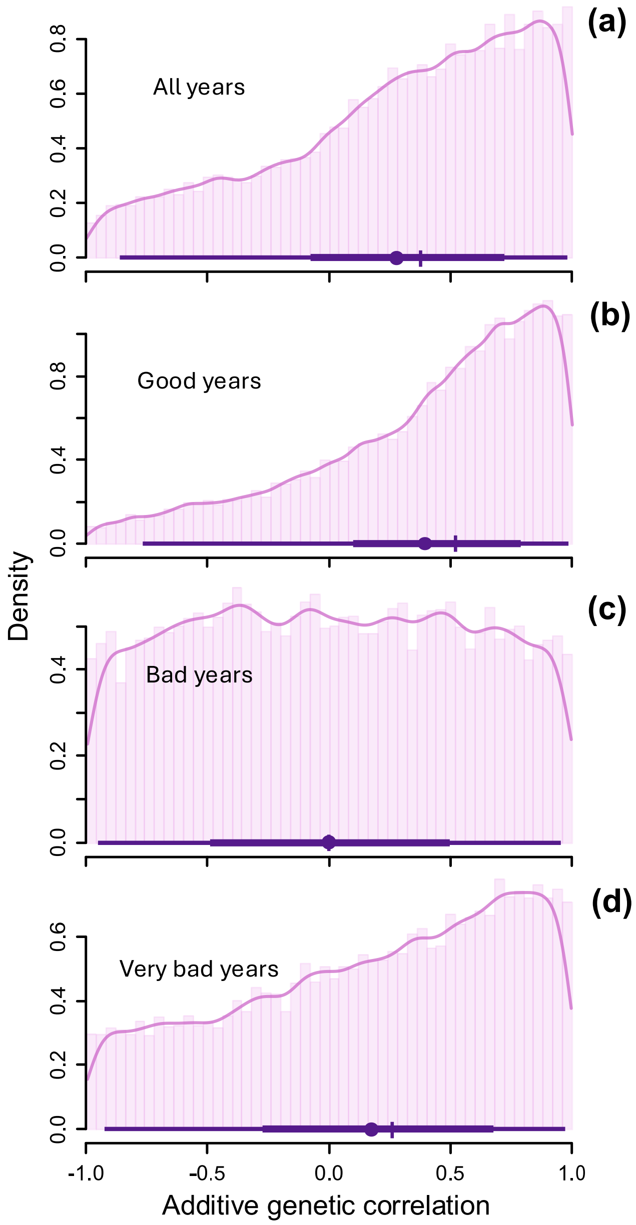


**Figure S18**. Posterior distributions of the cross-sex additive genetic correlation underlying annual adult survival (a) across all years ($V_{\mathrm{A}}$) and separately in years with (b) good, (c) bad, and (c) very bad environmental conditions ($\rho_{♀,♂}$, $\rho_{♀,♂}^{good}$, $\rho_{♀,♂}^{bad}$, and $\rho_{♀,♂}^{vbad}$, respectively). Histogram bars show the distribution of the posterior samples, with corresponding kernel density estimation in solid line. The superimposed thin and thick horizontal lines show the 95% and 50% credible intervals, respectively. Along these horizontal lines, the plain circle and vertical tick show the mean and median, respectively.

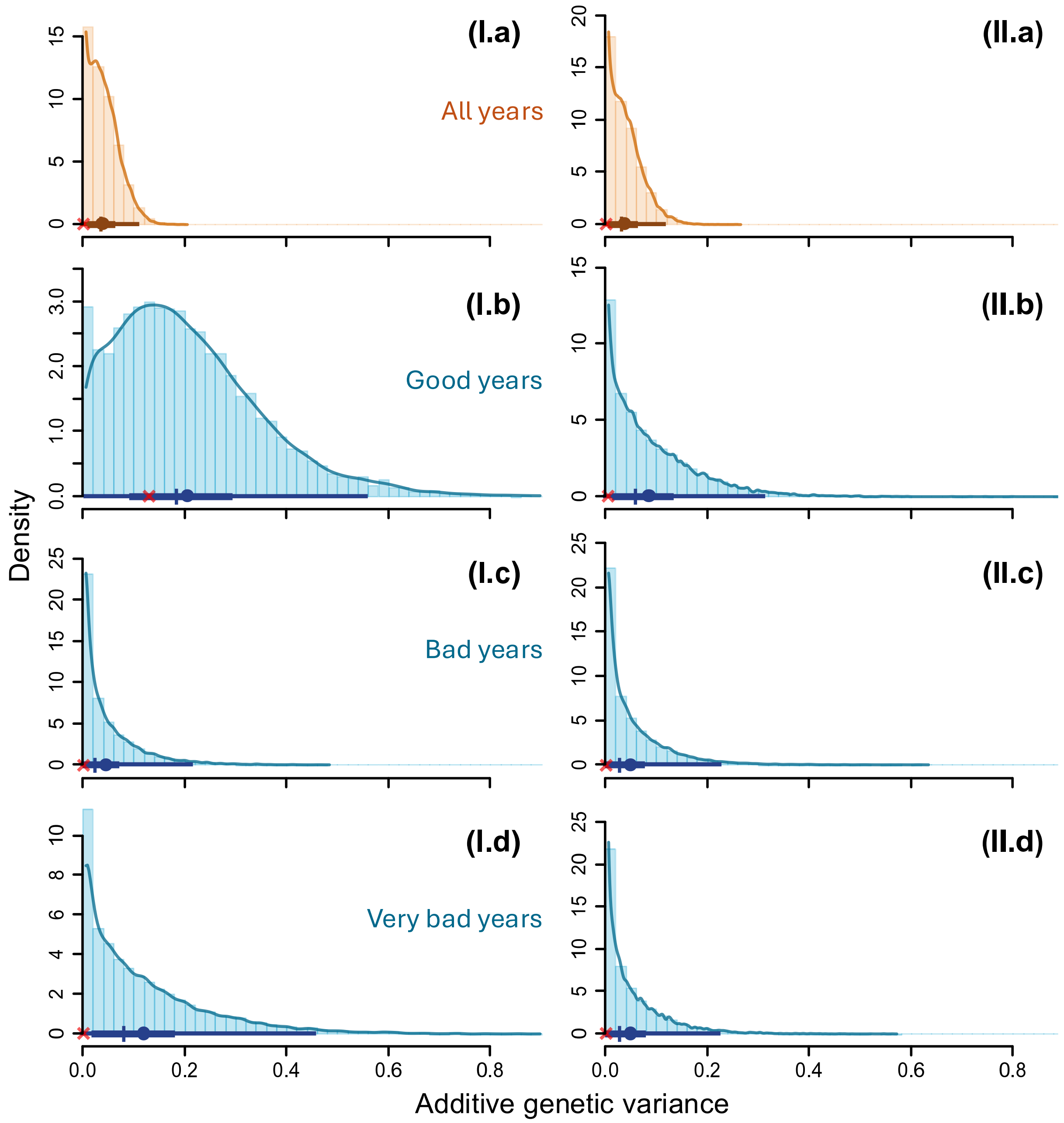


**Figure S19.** Posterior distributions of sex-specific additive genetic variance underlying annual adult survival estimated (I) in females and (II) in males, (a) across all years ($V_{\mathrm{A}}$) and separately in years with (b) good, (c) bad, and (c) very bad environmental conditions ($V_{\mathrm{A}}^{k}$). Histogram bars show the distribution of the posterior samples, with corresponding kernel density estimation in solid line. The superimposed thin and thick horizontal lines show the 95% and 50% credible intervals, respectively. Along these horizontal lines, the plain circle, vertical tick and red cross show the mean, median, and mode, respectively.

### Appendix S5
